# Two orthogonal MAP3K-driven pathways of NLRP1 inflammasome activation revealed by poisonous beetles

**DOI:** 10.64898/2026.01.23.701189

**Authors:** Rae Chua, Stephen Wearne, Shirley Ding, Toh Gee Ann, Muhammad Jasrie Firdaus, Lim Ying Shiang, Chai Yoke Tin, Seong Soo Lim, Ray Putra Prajnamitra, Foo Siu Wen, Melina Setiawan, Loh Yan Ping, Lee Sze Han, John E. A. Common, Wu Bin, Etienne Meunier, Franklin Zhong

## Abstract

Environmental toxins that cause irritant dermatitis remain poorly understood as activators of innate immune pathways. Here, we identify rove beetle (*Paederus*) and blister beetle (*Meloidae*) toxins as previously unrecognized triggers of the human NLRP1 inflammasome in keratinocytes. Rove beetles, likely through the ribosome inhibitor pederin, activate NLRP1 via translational stalling and the ZAKα-dependent ribotoxic stress response. In contrast, the phosphatase inhibitor cantharidin from blister beetles induces NLRP1 through TAK1-driven hyperphosphorylation of its linker region, independent of ZAKα. In their hyperactivated states, ZAKα and TAK1 share overlapping phosphosites on the NLRP1 disordered linker, including a common essential ‘TZ motif’. In addition, we show that TAK1 and ZAKα are jointly responsible for NLRP1 linker phosphorylation and activation caused by dsRNA and CHIKV infection. These findings reveal medically relevant insect toxins as activators of NLRP1, and uncover parallel MAP3 kinase pathways as converging upstream activating signals for the human NLRP1 inflammasome.

**GRAPHICAL ABSTRACT:** 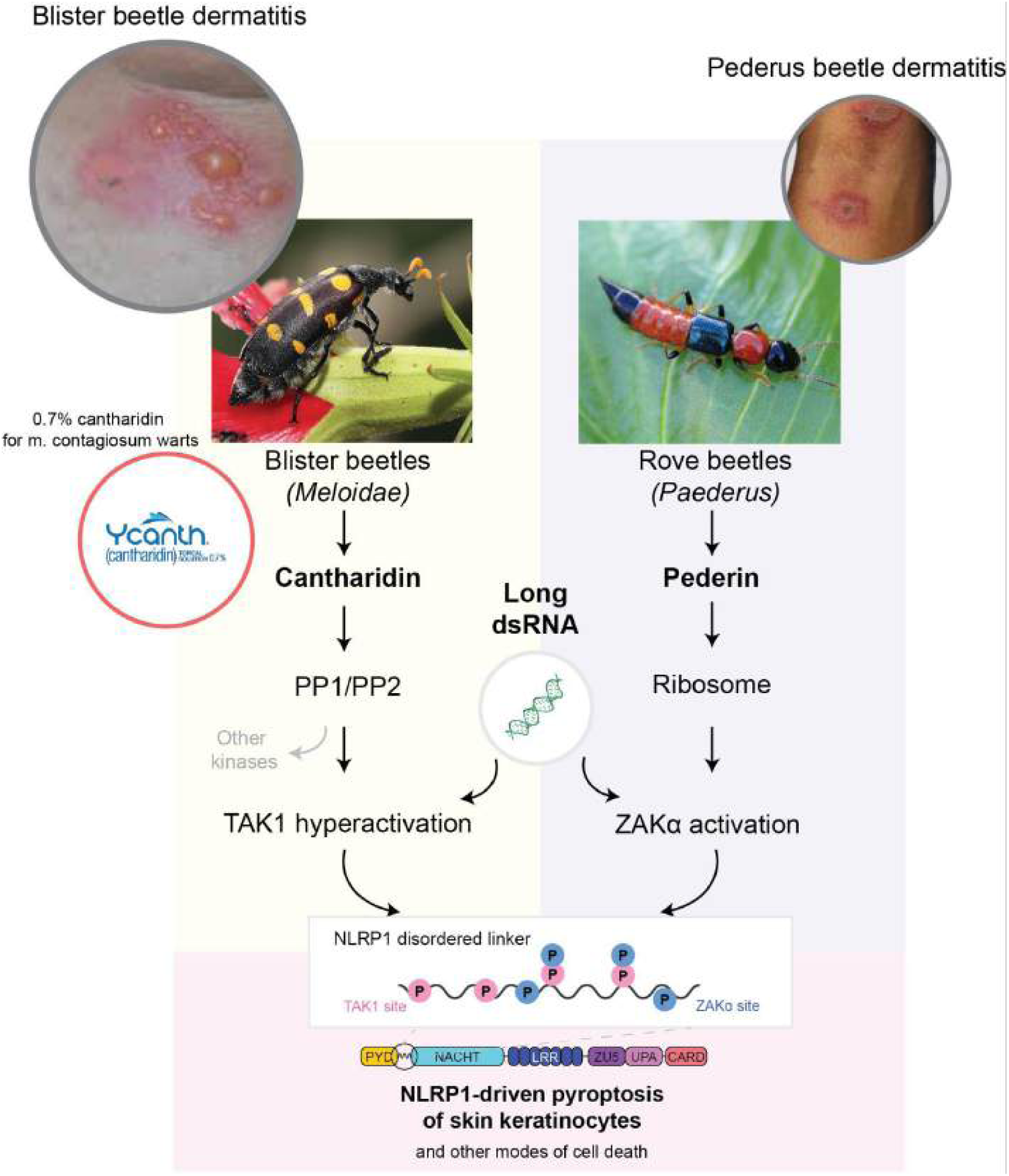

**KEY POINTS:** Two dermatitis-causing beetle species induce NLRP1-driven pyroptosis of human keratinocytes

Rove beetles, likely via pederin, activate NLRP1 via ribosome inhibition and ZAKɑ-driven RSR

Cantharidin from blister beetles activates human via TAK1-, instead of ZAKɑ-driven hyperphosphorylation of the NLRP1 linker region

Shared phosphosites by TAK1 and ZAKɑ on NLRP1 contribute to dsRNA-driven NLRP1 activation

## INTRODUCTION

Naturally occurring toxins have long served as powerful probes to study the innate immune system. Identifying the direct ligands/triggers of an innate immune sensor is often the most foundational step in the elucidation of novel immune signaling pathways. Most current studies on innate immune sensors, including those in the inflammasome pathway, have concentrated on a relatively narrow set of infectious triggers and danger signals, often overrepresented by ‘modern’ conditions encountered in the developed world. This bias overlooks the fact that the human innate immunity system interacts with a far broader array of external stimuli, both in the present day and in our evolutionary past. Much of the human ‘exposome’, including airborne pollutants, plant-derived irritants, and insect and marine toxins, can trigger or manipulate the human immune system in ways that remain poorly understood.

The NLRP1 inflammasome is an innate immune sensor highly expressed in human keratinocytes, where it can be activated by inhibition of certain cellular peptidases, viral proteases, dsRNA, reductive stress, and kinase signaling downstream of ribosome inhibition. Upon activation, NLRP1 assembles with the adaptor protein ASC and caspase-1 to drive IL-1β/IL-18 cytokine release and pyroptotic cell death (Barry et al. 2023; Yap et al. 2025; Bauernfried and Hornung 2022; Bachovchin 2021). Germline mutations in *NLRP1* result in Mendelian auto-inflammatory disorders such as MSPC and AIADK, all characterized by recurrent hyperplastic and inflammatory skin lesions (Zhong et al. 2016). In addition, common NLRP1 SNPs are genetic risk factors for skin diseases such as vitiligo and psoriasis (Fenini et al. 2022). These observations, together with its prominent expression in the skin, support the notion that NLRP1 functions as a key mediator of human skin immunity.

A number of naturally occurring toxins that activate NLRP1 have been described in the past few years (Robinson et al. 2022, 2023; Pinilla et al. 2023; Gorse et al. 2025); however, the full spectrum of NLRP1 triggers are still unknown and many questions remain regarding the upstream regulatory mechanisms governing NLRP1 activation. Identifying new NLRP1 triggers will thus shed further light on NLRP1 function and, by extension, how the human innate immune system defends against the broader skin exposome. In this study, we study two well-known insect species: blister beetles (genus: *Meloidae*) and rove beetles (genus: *Paederus*), both of which are well known to cause skin injury with ill-defined molecular mechanisms. Both beetle species are known pests and pose significant public health issues, particularly in tropical and subtropical regions where the beetles are found endemically and can swarm seasonally. For instance, local outbreaks of rove beetle dermatitis have been reported in many parts of Asia and Africa, and routinely trigger regional health alerts in affected countries during the summer months. Contact with these insects results in partially overlapping skin symptoms, ranging from erythematous plaques and vesicles to linear “whiplash”-like lesions. In both syndromes (known as blister beetle dermatitis and pederus dermatitis, respectively), keratinocyte death and inflammation are hallmarks of pathology. Although not usually life-threatening, both conditions lead to significant discomfort, lost productivity, and secondary infections. Current treatments are supportive and non-curative, to the best of our knowledge.

The causative toxins are known for both types of beetle dermatitis. Blister beetles release cantharidin (Bologna et al. 2008; Carrel et al. 1993), a vesicant terpenoid and potent inhibitor of the protein phosphatases PP1 and PP2A, enzymes involved in cell cycle regulation, apoptosis, and stress kinase signaling (Chen et al. 2002; Janssens and Goris 2001; Peng et al. 2002; Knapp et al. 1999; Williams et al. 2003; Lee et al. 2003; Zhang et al. 2025). Cantharidin has also been developed into an FDA-approved topical drug (YCANTH™) for the treatment of molluscum contagiosum, especially in pediatric patients, where destructive therapies are undesirable. The precise mode of action of cantharidin in therapeutic contexts—and how it drives both cytotoxicity and inflammation—also remains incompletely understood.

Rove beetles deliver pederin, a symbiont-derived polyketide that is known to block protein synthesis as well as transcription in mammalian cells (Kador et al. 2011; Piel et al. 2004; Brega et al. 1968). It is likely released from the beetle hemolymph and deposited onto the skin upon contact. Pederin has been postulated to be directly responsible for necrosis of keratinocytes and subsequent inflammation (Cressey et al. 2013; Assaf et al. 2010; Kalkman et al. 2024; Banney et al. 2000; Armstrong and Winfield 1969; Palaniappan and Karthikeyan 2023; Veraldi et al. 2013), but the mechanism remains unproven in skin-relevant assays. Pederin and its derivatives are also potently cytotoxic to tumor cell lines *in vitro*, sparking interest in their use as potential anti-cancer drugs (Narquizian and Kocienski 2000).

Here we report that exposure to both blister beetles and rove beetles results in NLRP1-driven pyroptosis of human keratinocytes, with each beetle species entailing a distinct upstream pathway. Rove beetle extract, likely via pederin, is an exquisitely potent inducer of ZAKɑ-dependent ribotoxic stress response (RSR), activating NLRP1 in a ZAKɑ-dependent manner with EC50 <0.01% of the body volume of an individual beetle (vol/vol) *in vitro*. By contrast, cantharidin found within blister beetles activates NLRP1 independently of ZAKɑ and p38 kinases, but instead via the hyperactivation of TAK1 following PP1/PP2 inhibition. In the hyperactivated state, TAK1 phosphorylates multiple sites in the NLRP1 linker region, including those targeted by ZAKɑ. Hence, TAK1 inhibition, KO or mutating the shared phosphorylation sites on NLRP1 prevents blister beetle or cantharidin-triggered pyroptosis of skin keratinocytes *in vitro* and in 3D skin organotypic cultures. Furthermore, we report that both ZAKɑ and TAK1 simultaneously contribute to dsRNA- and Chikungunya (CHIKV)-induced NLRP1 activation in keratinocytes via a common phosphorylation motif in the NLRP1 linker (TZ motif). Our study provides insight into the pathophysiological basis of beetle-induced dermatitis and uncovers additional regulatory mechanisms governing the activation of the human NLRP1 inflammasome. It also illustrates that underexplored insect-derived toxins are useful probes to dissect human innate immunity.

## RESULTS

### *Paederus* beetle extract induces ribotoxic stress and causes NLRP1-dependent pyroptosis

Pederin, along with closely related pseudopederin, is synthesized by the endosymbiotic *Pseudomonas spp* in *Paederus* beetles. More than 50 years ago, pederin was found to inhibit the translocation step of semi-purified yeast ribosomes (Barbacid et al. 1975). Related molecules in the pederin family, including mycalamide and onnamide, were subsequently shown to also inhibit the ribosome (Burgers and Fürst 2021). We prepared extracts of rove beetles (Figure 1A, Figure S1A) obtained from multiple geographical locations worldwide and tested their effects on protein synthesis and RSR-driven NLRP1 activation in a reporter system (A549-NLRP1-ASC-GFP). In this reporter cell line, ZAKɑ-driven NLRP1 activation can be quantitatively detected by the formation of ASC-GFP specks. De novo protein synthesis could be simultaneously measured using the BONCAT method employing AHA (Carlisle et al. 2023). Each beetle was dissolved in 100 µL of solvent, approximately the body volume of an individual beetle (Figure S1A). All 15 beetles tested were able to induce NLRP1 inflammasome assembly and inhibit protein synthesis in a dose dependent manner (Figure 1B-C). Some beetles were extraordinarily potent, with EC50 (ASC speck formation) and IC50 (translation inhibition) lower than 1/10,000 vol/vol; the differences in potency between individual beetles seem to be due to the storage duration between capture and extraction. We used mass spectrometry to quantify the relative pederin and pseudopederin concentrations in four beetles with the widest spread of biological activities (Figure 1C, S2A-G). A strong linear correlation was observed between translation inhibition, NLRP1 inflammasome assembly, and the concentration of pederin/pseudopederin (Figure 1D, S2H). These results extend previous published results on purified ribosomes to cultured human cells, demonstrating that rove beetle extracts indeed inhibit protein synthesis via the ribosome, thereby activating RSR and the NLRP1 inflammasome.

**Figure 1:**
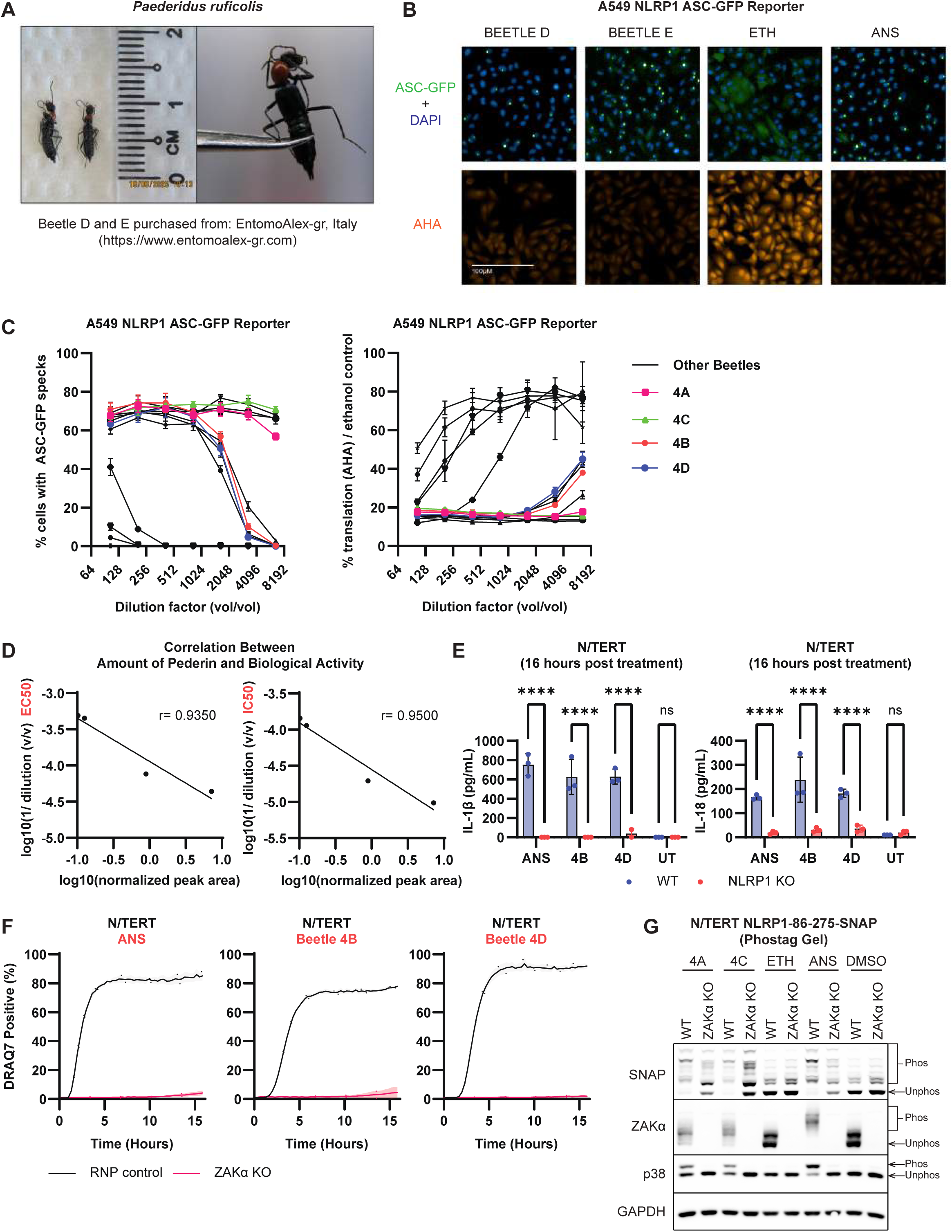
*Paederus* Beetle Extract Induces ZAKα-dependent Pyroptosis in Epithelial Cells. **A.** *Paederidus ruficolis* beetles obtained from Italy and used in the following experiments. **B.** Representative fluorescence images of A549 NLRP1 ASC-GFP reporter cells following treatment with 52X dilution of beetle D extract, 104X dilution of beetle E extract, ethanol (vehicle control for beetle extracts), and 1 μM anisomycin. Cells were fixed and stained with DAPI after 3 hours of treatment. AHA-labeling was done with AZDye™ 568 Alkyne. Scale bar, 100 μM. **C.** Percentage of ASC-GFP speck forming cells (left) and percentage of AHA incorporation/ vehicle control, as a proxy for translation inhibition (right), was quantified for all beetles. A total of 15 beetles were crushed and titrated on A549 NLRP1 ASC-GFP reporter cells. Highlighted beetles 4A-4D were used in the following functional experiments. **D.** Correlation between the amount of pederin detected by mass spectrometry and EC50 or IC50 of beetles 4A-4D, as proxies for biological activity. Simple linear regression was performed and Pearson correlation (r value) was calculated in GraphPad Prism. **E.** IL-1β and IL-18 ELISA from WT and NLRP1 KO N/TERTs following treatment with 1 μM anisomycin, or X1500 dilution of beetle 4B and beetle 4D extracts. Cell culture media was harvested after 16 hours of treatment. **F.** Quantification of the percentage of DRAQ7-positive ZAKα KO and RNP control N/TERT cells following treatment with 1μM anisomycin, X1500 dilution of beetle 4B and beetle 4D extracts. **G.** Rhodamine labelled SNAP-tagged NLRP1-DR and immunoblots of ZAK, p38, and GAPDH following PhosTag-SDS-Agarose-PAGE. ZAKα KO and RNP control N/TERT cells expressing NLRP1-86-275-SNAP were pre-treated with 5 μM emricasan (pan-caspase inhibitor) before stimulation with beetle 4A and beetle 4C extracts, ethanol (vehicle control for beetle extracts), 1μM anisomycin, DMSO (vehicle control). Cells were harvested after 3 hours of treatment. Phosphorylated and unphosphorylated species of each protein are indicated. Error bars represent SEM from three technical replicates, where one replicate refers to an independent sample. Significance values were calculated based on two-way ANOVA followed by Sidak’s test for multiple pairwise comparisons in (E). ns, nonsignificant; *P < 0.05; ****P < 0.0001.

We then tested the effect of rove beetle extracts on immortalized human keratinocytes, N/TERT-1 (hereby referred to as N/TERT cells). All beetle extracts caused bona fide inflammasome activation and pyroptosis in N/TERT cells, as measured by IL-1β/IL-18 secretion, GSDMD cleavage and rapid membrane lysis. Genetic deletion of either NLRP1 (Figure 1E, S1B-F) or ZAKɑ abrogated these effects (Figure 1F, S3A-F), but did not affect non-inflammasome-related cell death as shown by GSDME cleavage (Figure S1B and E, Figure S3A and D). Furthermore, harringtonine run-off assays performed directly on N/TERT cells confirmed that the beetle extracts caused ribosome stalling, similarly to the well-known A site inhibitor, anisomycin (Figure S3G-H). Beetle extracts also caused ZAKɑ autophosphorylation and p38 activation, as well as hyperphosphorylation of the NLRP1 disordered linker region-hallmark events in RSR-driven NLRP1 activation (Figure 1G). Taken together, these results demonstrate that rove beetles, likely via pederin and pseudopederin, cause ZAKɑ- and NLRP1-driven pyroptosis of human keratinocytes, which likely accounts for their toxic effects on human skin.

Despite repeated attempts, we have been unable to isolate pederin directly from rove beetles or obtain purified pederin from commercial and academic sources. Therefore we cannot currently rule out the possibility that molecules other than pederin/pseudopederin contribute to the ability of rove beetles to inhibit the ribosome and activate NLRP1. That said, the strong correlation between the activity of the extract and pederin levels, along with older literature on pederin, suggests that this is the case. Based on published pederin concentrations per beetle (∼2 µM) and assuming that the pederin/pseudopederin is the sole pyroptosis-inducing toxin in these beetles, our results imply that the IC50 for ribosome inhibition and EC50 of NLRP1 activation of pederin is less than 0.2 nM, making it one of the strongest ribosome inhibitors and RSR inducers. This is consistent with the fact that simply brushing the beetle on exposed skin is sufficient to cause Paederus dermatitis. This observation is consistent with the finding that simple contact, such as brushing the beetle against bare skin, is enough to trigger Paederus dermatitis.

### Cantharidin from Blister Beetles induces mixed modes of cell death involving NLRP1-driven pyroptosis

We next studied the effects of blister beetles of the genus Mylaris. The vesicant, or skin blistering properties of these beetles are attributed to cantharidin and its hydrolysis product cantharidic acid, both of which inactivate the human PP1 and PP2 phosphatase complexes with IC50 in the low micromolar ranges (Figure 2A) (Honkanen 1993; McCluskey et al. 2001; Sakoff et al. 2002). PP1 and PP2a/b are conserved macromolecular assemblies that dephosphorylate phospho-serine and phospho-threonine residues on a large number of substrate proteins, thus regulating diverse cellular processes such as cell cycle, transcription and cell motility (Shi 2009; Cohen 1989; Brautigan and Shenolikar 2018; Ingebritsen and Cohen 1983). Cantharidin and other PP1/PP2 targeting natural toxins, including okadaic acid, calyculin and nodularin are routinely used as tool compounds to investigate the functions of PP1/PP2 in vitro and in vivo. In addition, cantharidin has been developed as a topical drug to remove skin warts caused by molluscum contagiosum infection (brand name Y-canth) (Gupta et al. 2023). Other naturally occurring PP1 and PP2 inhibitors, such as okadaic acid are also being investigated for medical use (Stanford and Bottini 2023; Naz et al. 2020).

**Figure 2.**
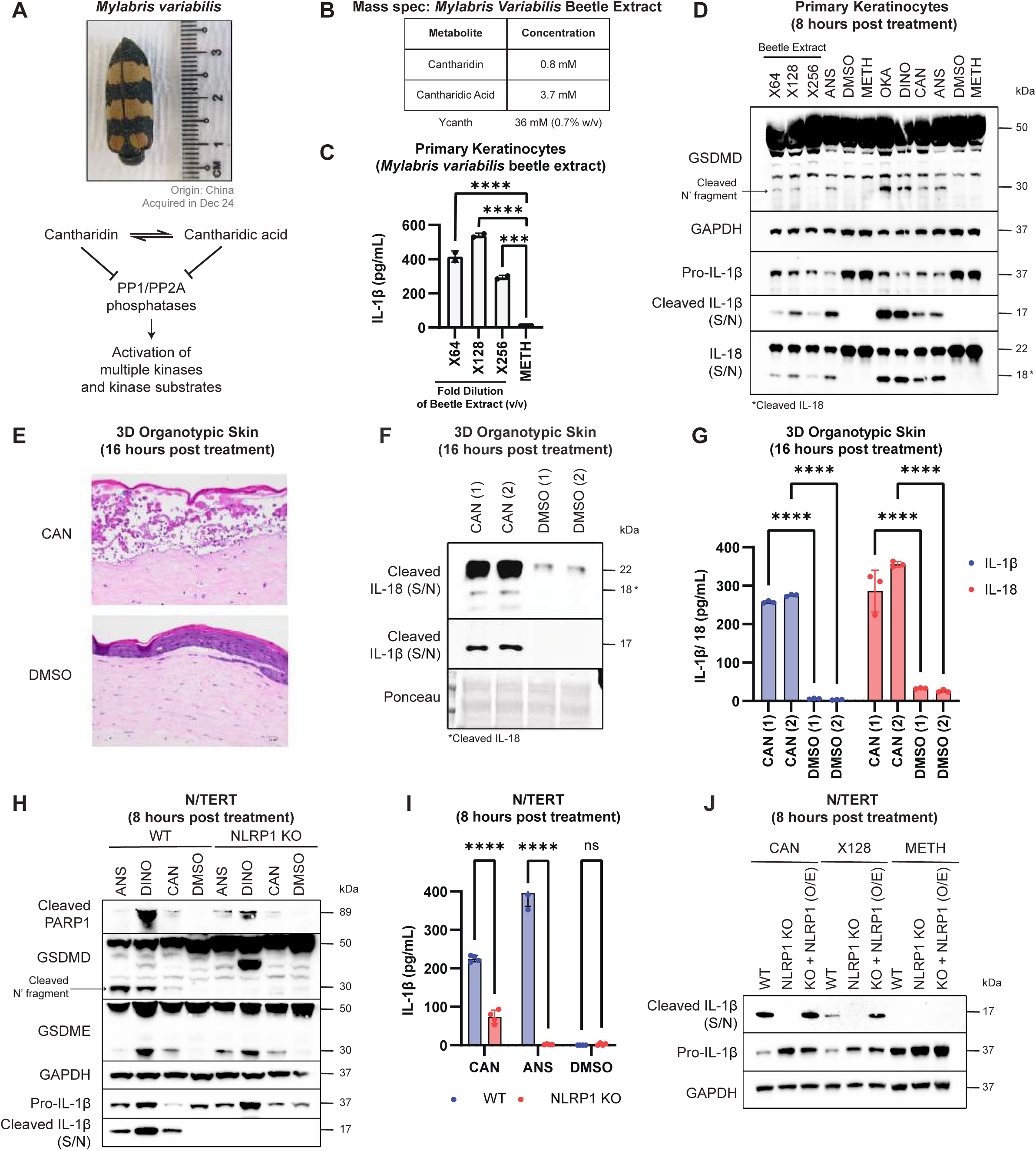
Mylabris beetle extract induces mixed mode of cell death involving NLRP1 in the skin. **A.** *Mylabris variabilis* beetle obtained from China and used in the following experiments. Cantharidin exists in equilibrium with its natural derivative, cantharidic acid. Both cantharidin and cantharidic acid inhibit protein phosphatases (PP1 and PP2A), likely leading to activation of cellular kinases. **B.** Concentration of cantharidin and cantharidic acid from beetle extracts detected by mass spectrometry. Ycanth, a topical therapeutic for molluscum contagiosum, contains 36 mM of Cantharidin. **C.** IL-1β ELISA from primary keratinocytes following treatment with several dilutions of *Mylabris variabilis* beetle extract and methanol (vehicle control for beetle extract). Cell culture media was harvested after 16 hours of treatment. **D.** Immunoblot of GSDMD (full length and cleaved), GAPDH, Pro-IL-1β, cleaved IL-1β, and IL-18 (full length and cleaved) from primary keratinocytes following treatment with several dilutions of *Mylabris variabilis* beetle extract, 1 μM anisomycin, DMSO (vehicle control), methanol (vehicle control for beetle extract), 62.5 nM okadaic acid, 12.5 nM dinophysis, and 50 μM cantharidin. Cells, floaters and cell culture media were harvested after 8 hours of treatment. **E.** Representative images of 3D organotypic skin sections stained with H&E, **F.** immunoblot of IL-18 (full length and cleaved) and cleaved IL-1β, or **G.** IL-1β and IL-18 ELISA following treatment with 50 μM cantharidin and DMSO (vehicle control). Two technical replicates are presented. 3D organotypic skin was fixed and cell culture media was harvested after 16 hours of treatment. **H.** Immunoblot of cleaved PARP1, GSDMD (full length and cleaved), GSDME (full length and cleaved), Pro-IL-1β, cleaved IL-1β, and GAPDH from WT and NLRP1 KO N/TERTs following treatment with 1 μM anisomycin, 12.5 nM dinophysis, 50 μM cantharidin, and DMSO (vehicle control). Cells, floaters, and cell culture media were harvested after 8 hours of treatment. **I.** IL-1β ELISA of WT and NLRP1 KO N/TERTs following treatment with 50 μM cantharidin, 1 μM anisomycin, and DMSO (vehicle control). Cell culture media was harvested after 8 hours of treatment. **J.** Immunoblots of cleaved IL-1β, Pro-IL-1β, and GAPDH from WT, NLRP1 KO, and NLRP1 KO N/TERTs overexpressing GFP_NLRP1-86-C_FLAG (WT) following treatment with 50 μM cantharidin, X128 dilution of *Mylabris variabilis* beetle extract, and methanol (vehicle control for beetle extract). Cells and cell culture media were harvested after 8 hours of treatment. Error bars represent SEM from three technical replicates, where one replicate refers to an independent sample. Significance values were calculated based on one-way ANOVA followed by Dunnett’s test (C) or two-way ANOVA followed by Sidak’s test for multiple pairwise comparisons (G and I). ns, nonsignificant; *P < 0.05; ****P < 0.0001.

We prepared crude extracts from preserved blister beetles (Mylabris variabilis) procured from a Chinese medicine store (Figure S4A-D). Mass spectrometry analysis of extract using authentic standards confirmed that both cantharidin and cantharidic acid are present in the millimolar range (Figure 2B and Figure S4E-F). Treatment of primary human keratinocytes as well as N/TERT cells with blister beetle extract caused rapid cell death within hours, characterized by mixed apoptotic and pyroptotic features, with a subset of cells becoming DRAQ7-permeable within 3 hours (Figure S5A-B). Exposure to blister beetle extract also led to caspase 1-dependent maturation and secretion of IL-1β and IL-18, as well as GSDMD cleavage in a dose-dependent manner (Figure 2C-D, S5C-D, S6A-C). A similar pro-pyroptotic effect was also observed with PP1/PP2-inhibiting dinoflagellate toxins okadaic acid (OKA), dinophysistoxin (DINO) and cyanobacterial toxin calyculin A, whereas extract from a related, non-blistering beetle in the Tenebrionidae family (darkling beetles, Blaps gibba) did not cause overt cytotoxicity in cultured keratinocytes (Figure S7A-F and Figure S8A-F). The concentration of cantharidin and cantharidic acid required to elicit keratinocyte pyroptosis is estimated to be ∼3-15 µM (∼1/250 of body volume of an individual blistering beetle), more than 1000 fold less than the concentration of cantharidin used in Y-canth (Figure 2B-C, S6A-B). To more faithfully model the effect of blister beetle exposure to skin, we treated 3D reconstructed skin models with purified cantharidin. While a lower concentration (∼3 µM) did not cause visible histological changes (Figure S5J), 50 µM or 12.5 µM cantharidin caused massive keratinocyte cell death, resulting in dramatic disruption and detachment of the epidermal layer from the dermal scaffold in multiple biological replicates (Figure 2E and S5J). This was accompanied by the release of mature IL-1β and IL-18 into the culture media (Figure 2F-G and Figure S5G-I). Thus, similar to rove beetles, blister beetles also cause pyroptosis of human skin keratinocytes.

Next, we investigated the mechanisms of cantharidin-induced pyroptosis. Deletion of NLRP1, ASC or chemical inhibition of caspase-1 abrogated N/TERT pyroptosis caused by not only cantharidin (Figure 2H-J, Figure S5C-F, Figure S6C-K, and Figure S7F), but also other PP1/PP2 inhibitors, including OKA, DINO and calyculin (Figure S8B-F). Re-expressing wild-type NLRP1 in NLRP1 KO N/TERTs restored cantharidin-induced IL-1β cleavage and secretion (Figure 2J). Consistent with the essential nature of PP1/PP2 phosphatase complexes, non-pyroptotic death-characterized by cell rounding as well as GSDME and PARP1 cleavage occurred independently of NLRP1 following cantharidin treatment (Figure 2H, Figure S5E, Figure S6E, Figure S7D, and Figure S8B). Thus, PP1/PP2 inhibitors are a new class of NLRP1 activators that trigger mixed modes of cell death encompassing inflammasome-driven pyroptosis in human keratinocytes. This may account for the unique vesicant properties of blister beetles, as well as the efficacy of Y-canth as a topical wart removal agent.

### ZAKɑ and p38 are not involved in cantharidin-induced NLRP1 DR linker phosphorylation and subsequent pyroptosis

As shown with pederin (this study), UVB irradiation and certain bacterial exotoxins, ribotoxic stress response (RSR) constitutes one of the major parallel pathways that drive human NLRP1 inflammasome activation in human skin. Mechanistically, RSR leads to rapid hyperphosphorylation of the NLRP1 disordered linker region by MAP3K20/ZAKɑ and its downstream effector, p38. As PP1/PP2 phosphatase complexes are known negative regulators of MAPK pathways, we thus tested if ZAKɑ, p38 and NLRP1 linker phosphorylation are also required for cantharidin-induced NLRP1 activation. Deletion of the N-terminal non-oligomerizing PYD did not impact the ability of cantharidin to activate the NLRP1 inflammasome, but further deletion of the linker region (a.a. 86-254) (Figure 3A and S9C) rendered NLRP1 insensitive to blister beetle extract, cantharidin and other PP1/PP2 inhibitors (Figure 3C and S10D), but not VbP (Figure S9C-G, and Figure S10D). Similarly, mutating all the serine and threonine residues to alanine in the linker region (STless) (Figure S9D) also selectively blocked cantharidin-triggered NLRP1 activation (Figure 3D-E and Figure S9H-I).

**Figure 3:**
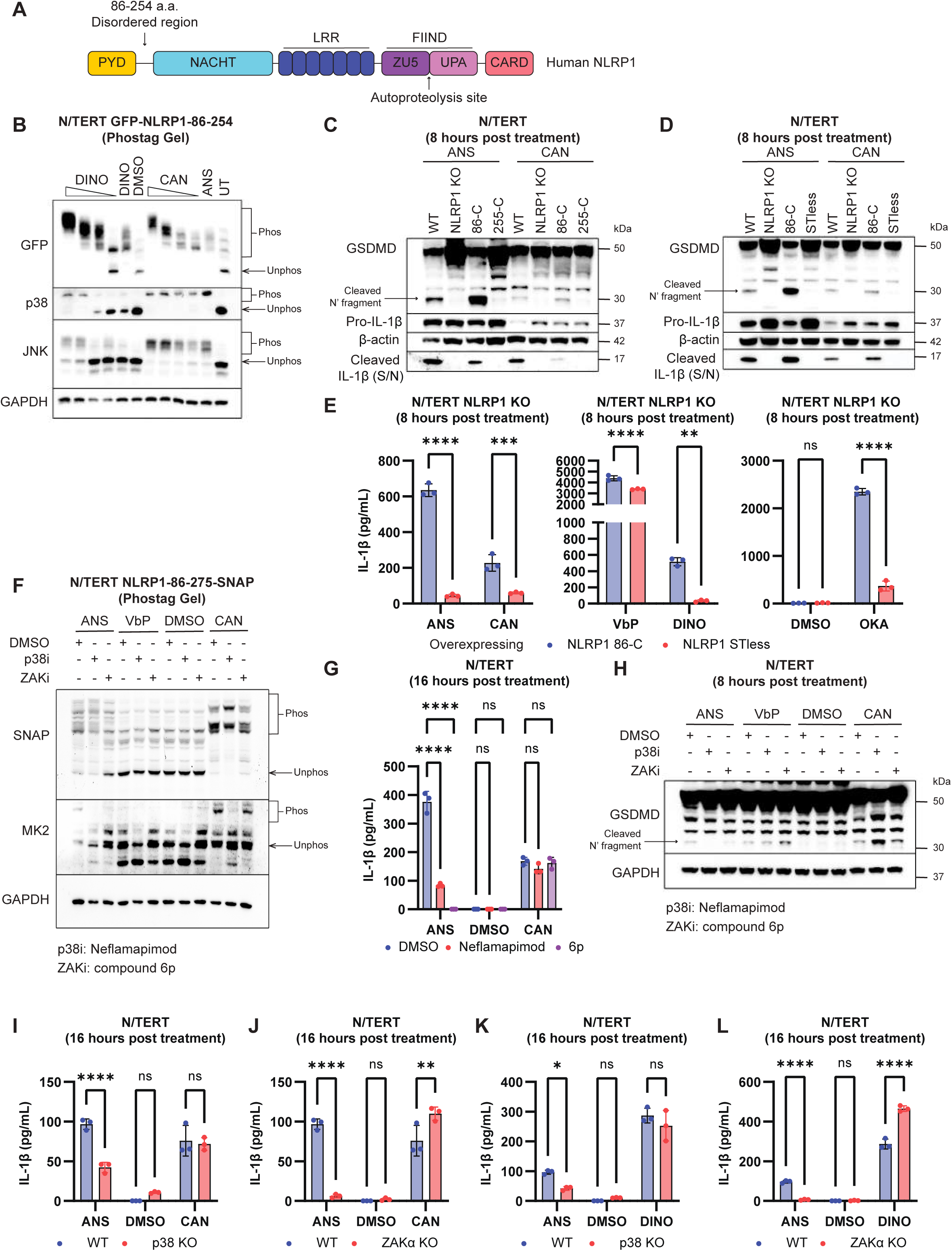
Cantharidin-dependent phosphorylation of the NLRP1 DR and activation of the NLRP1 inflammasome is independent of ZAKα and p38. **A.** Domain architecture of human NLRP1. The disordered region (NLRP1-DR residues 86-254 a.a.) lies after the PYD. **B.** Immunoblot of GFP-tagged NLRP1-DR, p38, JNK, and GAPDH following PhosTag-SDS-Agarose-PAGE. NLRP1 KO N/TERTs overexpressing NLRP1-86-254_GFP were pre-treated with 5μM emricasan (pan-caspase inhibitor) before stimulation with a titration of dinophysis, cantharidin, DMSO (vehicle control), and 1 μM anisomycin. Cells were harvested after 3 hours. Phosphorylated and unphosphorylated species of each protein are indicated. **C.** Immunoblot of GSDMD (full length and cleaved), Pro-IL-1β, cleaved IL-1β, and β-actin from NLRP1 KO N/TERTs overexpressing GFP_NLRP1-86-C_FLAG (WT), a truncated GFP_NLRP1-255-C_FLAG, or **D.** the GFP_NLRP1-86-C_FLAG (STless) mutant following treatment with 1 μM anisomycin and 50 μM cantharidin. Cells, floaters, and cell culture media were harvested after 8 hours of treatment. **E.** IL-1β ELISA from NLRP1 KO N/TERTs overexpressing GFP_NLRP1-86-C_FLAG (WT) or the GFP_NLRP1-86-C_FLAG (STless) mutant following treatment with 1 μM anisomycin, 50 μM cantharidin, 6 μM VbP, 12.5 nM dinophysis, DMSO (vehicle control), and 62.5 nM okadaic acid. Cell culture media was harvested after 8 hours of treatment. **F.** Rhodamine labelled SNAP-tagged NLRP1-DR and immunoblot of MK2 and GAPDH following PhosTag-SDS-Agarose-PAGE. N/TERTs overexpressing NLRP1-86-275_SNAP were pre-treated with 5μM emricasan (pan-caspase inhibitor) and DMSO (vehicle control), 1 μM of Neflamapimod (p38 inhibitor), or 6p (ZAK inhibitor) before stimulation with 1 μM anisomycin, 3 μM VbP, DMSO (vehicle control), and 50 μM cantharidin. Cell lysates were harvested after 3 hours of treatment. Phosphorylated and unphosphorylated species of each protein are indicated. **G.** IL-1β ELISA from N/TERTs pre-treated with 1 μM 6p (ZAK inhibitor), 1 μM of neflamapimod (p38 inhibitor), or DMSO (vehicle control) before stimulation with 1 μM anisomycin, DMSO (vehicle control), and 50 μM cantharidin. Cell culture media was harvested after 16 hours of treatment. **H.** Immunoblot of GSDMD and GAPDH from N/TERTs pre-treated with DMSO (vehicle control), 1 μM of Neflamapimod (p38 inhibitor), or 6p (ZAK inhibitor) before stimulation with 1 μM anisomycin, 3 μM VbP, DMSO (vehicle control), and 50μM cantharidin. Cells and floaters were harvested after 8 hours of treatment. **I-J.** IL-1β ELISA from p38 KO or ZAKα KO N/TERTs following treatment with 1 μM anisomycin, DMSO (vehicle control), 50 μM cantharidin, or **K-L.** 12.5 nM dinophysis. Cell culture media was harvested after 16 hours of treatment. Error bars represent SEM from three technical replicates, where one replicate refers to an independent sample. Significance values were calculated based on two-way ANOVA followed by Sidak’s test for multiple pairwise comparisons (E, G, and I-L). ns, nonsignificant; *P < 0.05; ****P < 0.0001.

PhosTag analysis confirmed that the NLRP1 linker, when expressed in isolation as a tagged fusion protein (Figure S9A-B), undergoes dramatic phosphorylation in response to blister beetle extract, purified cantharidin and other PP1/PP2 inhibitors. This is accompanied by phosphorylation of SAPK members of the MAPK pathway, p38 and JNK (Figure 3B and Figure S10A-C), in agreement with broad substrate specificity of PP1/PP2.

These results initially led us to hypothesize that cantharidin also activates NLRP1 via ZAKɑ- and p38-mediated NLRP1 phosphorylation likely by removing the endogenous negative regulation of these kinases imposed by PP1/PP2. However, ZAKɑ and p38 inhibitors, and genetic deletion of ZAK and p38 caused negligible changes to the degree of NLRP1 activation triggered by cantharidin or other PP1/PP2 inhibitors, as measured by IL-1β secretion, GSDMD cleavage (Figure 3G-L and S10E-F), or NLRP1 linker phosphorylation (Figure 3F). In some instances, ZAKɑ and p38 inhibition even potentiated cantharidin-induced pyroptosis (Figure 3F-L and Figure S10E-F). These results indicate that, in contrast to pederin and other ribotoxins, cantharidin and other PP1/2 inhibitors engage a different pathway to phosphorylate and activate NLRP1 independently of ZAKɑ and p38. We ruled out JNK and ERK as the upstream kinases driving NLRP1 activation by cantharidin (Figure S10G-K).

We devised a chemical screening approach to identify the responsible upstream kinase(s). A custom panel of kinase inhibitors were tested for their effects on dinophysistoxin/cantharidin-induced IL-1β secretion and NLRP1 linker hyperphosphorylation in N/TERT cells expressing an NLRP1-SNAP linker fusion protein (Figure 4A). All compounds were screened initially at 5 µM, above the reported IC50 against the intended kinase targets. Five inhibitors scored as hits in both assays: 5z-7-oxozeanol, dabrafenib, vemurafenib, NG25, as well as the pan-kinase inhibitor staurosporine (Figure 4A, S11A-C), with all retaining activity at 1 µM. As three molecules all target multiple kinases, we selected 6 well-characterized kinase inhibitors, Flavopiridol, BMS-345541, Nilotinib, SB230580, Torin 2, and Tofacitinib that had no effects on NLRP1 activation and NLRP1 linker phosphorylation (Figure S11D-G). Sorafenib was excluded in subsequent analyses due to its acute cellular toxicity. By intersecting the annotated targets of 4 positive inhibitors (5z-7-oxozeanol, dabrafenib Vemurafenib and NG25) and excluding the targets of the 6 negative compounds as defined in the KINOMEscan database (Keenan et al. 2018) (https://lincs.hms.harvard.edu/kinomescan/), we shortlisted 8 candidate kinases that could potentially contribute to NLRP1 activation following PP1/PP2 inhibition. Among the 8 kinases, the only serine/threonine kinase was MAP3K7/TAK1, which has well-established functions in the innate immune system, including cytokine signaling and cell death regulation (Figure 4A) (Xu and Lei 2020; Tan et al. 2017).

**Figure 4:**
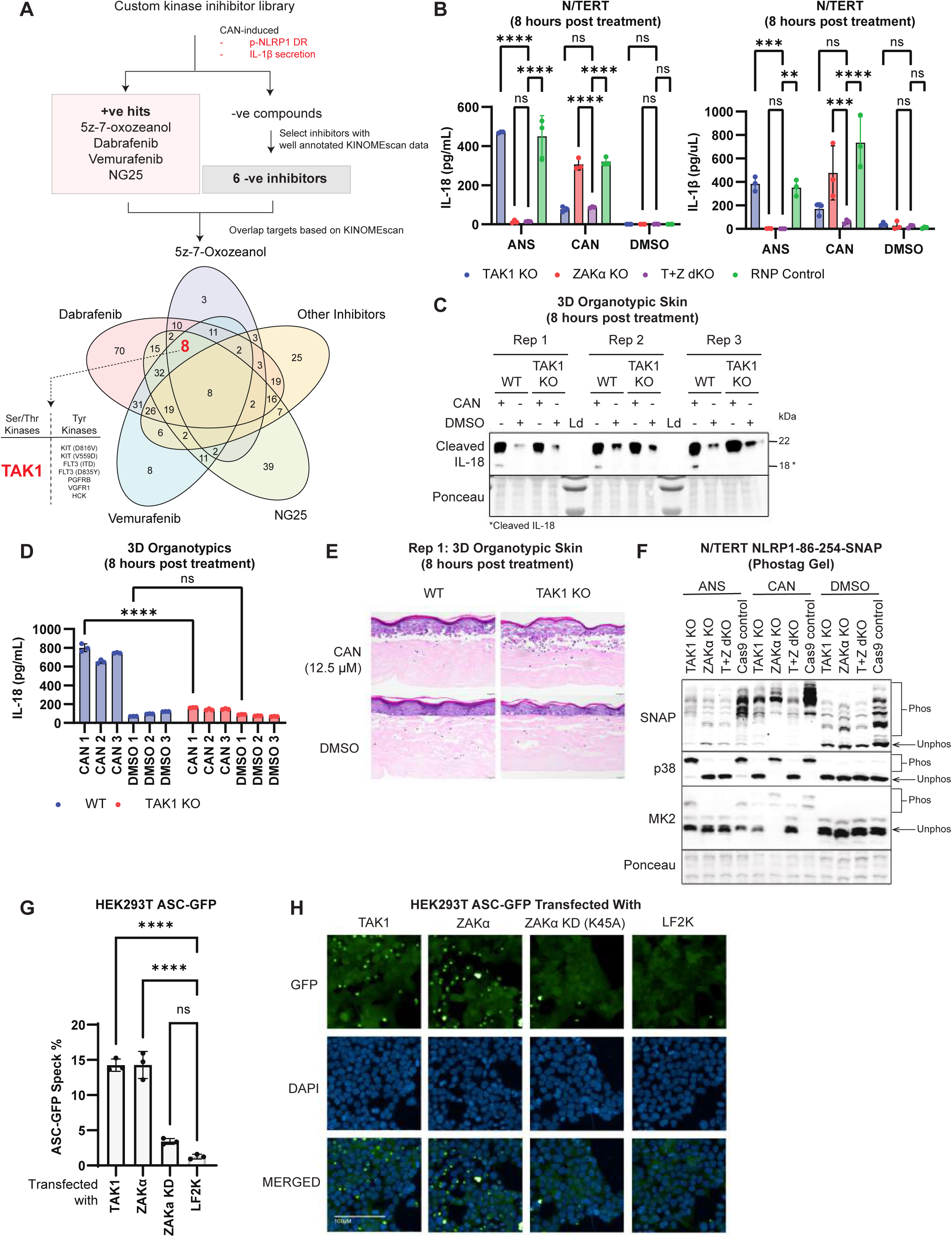
Chemical screen identifies TAK1 as the MAP3K responsible for cantharidin-dependent phosphorylation and activation of the NLRP1 inflammasome. **A.** Schematic describing the experimental workflow using well-characterized kinase inhibitors and the publicly available KINOMEscan database to identify TAK1 as the MAP3K responsible for cantharidin-dependent phosphorylation. **B.** IL-1β and IL-18 ELISA from TAK1 KO, ZAKα KO, T+Z dKO, and RNP control N/TERTs following treatment with 1 μM anisomycin, 50 μM cantharidin, and DMSO (vehicle control). Cell culture media was harvested after 8 hours of treatment. **C.** Immunoblot of IL-18 (cleaved and FL), **D.** IL-18 ELISA, or **E.** representative H&E images of WT and TAK1 KO 3D organotypic skin following treatment with 50 μM cantharidin and DMSO (vehicle control). 3D skin organotypics were fixed and culture media was harvested after 8 hours of treatment. **F.** Rhodamine labelled SNAP-tagged NLRP1-DR following PhosTag-SDS-Agarose-PAGE and immunoblot of p38 and MK2 from TAK1 KO, ZAKα KO, T+Z dKO, and RNP control N/TERTs following treatment with 1 μM anisomycin, 50 μM cantharidin, and DMSO (vehicle control). Cells were harvested after 3 hours of treatment. Phosphorylated and unphosphorylated species of each protein are indicated. **G.** Percentage of ASC-GFP speck forming or **H.** representative fluorescence images of HEK239T NLRP1 ASC-GFP reporter cells transfected with cDNA of TAK1, ZAKα, ZAKα kinase dead mutant (K45A), and LF2K (transfection control). Cells were fixed and stained with DAPI 24 hours after transfection. Error bars represent SEM from three technical replicates, where one replicate refers to an independent sample. Significance values were calculated based on one-way ANOVA followed by Dunnett’s test (G) or two-way ANOVA followed by Sidak’s test for multiple pairwise comparisons (B and D). ns, nonsignificant; *P < 0.05; ****P < 0.0001.

Among 5 kinase inhibitors that inhibit cantharidin-induced NLRP1 activation, 5z-7-oxozeanol (abbreviated as 5z7 hereafter) and NG25 are often used tool inhibitors for TAK1; however, both molecules also cross-inhibit ZAKɑ with IC50s in the submicromolar range (https://lincs.hms.harvard.edu/kinomescan/). Specifically, at 1 µM, 5z7 causes greater inhibition of ZAKɑ than TAK1 in vitro, as documented by the original report on its synthesis and characterization (Tan et al. 2017). We confirmed 5z7’s ability to inhibit ZAKɑ in N/TERT cells (Figure S11B-C). Another reported TAK1 inhibitor known as NG25 also cross-reacts with ZAKɑ (IC50 < 1µM) (https://lincs.hms.harvard.edu/kinomescan/). However, the reported ZAKɑ inhibitor, compound 6p, does demonstrate strong selectivity for ZAKɑ, and has no measurable effect on TAK1 in a KINOMEscan assay (Yang et al. 2020).

To extend our experiments beyond chemical inhibitors, we generated TAK1 KO, ZAKα KO and TAK1+ZAKα double KO (T+Z dKO) N/TERT cells (Figure S12A) and compared their response to PP1/PP2 inhibitors vs. ribosome inhibitors. A clear divergence was observed: while ZAK deletion selectively abrogated anisomycin-induced inflammasome activation, cantharidin-induced inflammasome activation was only lost in TAK1 KO cells, but not ZAK KO cells. T+Z dKO cells failed to respond to either anisomycin or cantharidin as measured by IL-1β/IL-18 secretion and GSDMD cleavage (Figure 4B and Figure S12B-C). The requirement for TAK1 was further validated in 3D skin cultures reconstructed with TAK1 KO primary human keratinocytes, as TAK1 KO 3D skin no longer secreted mature IL-18 after cantharidin exposure (Figure 4C-D). Nonetheless, TAK1 KO did not offer histologically significant protection against the disintegration of the epidermal layer (Figure 4E and Figure S12D). This is consistent with the observation that cantharidin-treated TAK1 KO N/TERT cells showed similar levels of apoptotic markers, including PARP1 and GSDME cleavage, and underwent similar morphological changes as control cells (Figure S12E-F). These results support the notion that PP1/PP2 inhibition not only leads to pyroptosis driven by the TAK1-NLRP1 axis, but also apoptotic cell death.

Using PhosTag analysis of the tagged NLRP1 disordered linker, we further demonstrated that TAK1 deletion preferentially reduced NLRP1 phosphorylation caused by cantharidin instead of anisomycin, whereas the opposite was true for ZAKɑ (Figure 4F). Interestingly, TAK1 KO also largely eliminated p38 and MK2 phosphorylation caused by cantharidin, dinophysistoxin and blister beetle extract (Figure 4F and Figure S12G-I). As p38 and MK2 are themselves not involved in cantharidin-triggered NLRP1 activation (Figure 3F-L), these results indicate that TAK1 likely controls a larger stress response encompassing p38, MK2 and the NLRP1 inflammasome downstream of PP1/PP2 inhibition. Next we tested if TAK1 was sufficient to drive NLRP1 activation in the absence of general PP1/PP2 inhibition. Indeed, when overexpressed in the commonly used HEK293T-NLRP1-ASC-GFP reporter system, TAK1 causes a similar level of ASC-GFP speck formation as ZAKɑ (Figure 4G-H).

TAK1 is a well-established signal transducer for proinflammatory cytokines such as IL-1β and TNFɑ, as well as various TLR ligands. However, none of these stimuli are sufficient to activate NLRP1. This may seem at odds with our results, which point to TAK1 as a critical upstream switch for NLRP1. To resolve this paradox, we performed a head-to-head comparison between cantharidin and TNFɑ, with the phosphorylation of TAK1, p38, JNK and ERK as a readout for the TAK1-driven MAPK output. This analysis reveals that the two stimuli cause vastly different levels and patterns of TAK1 and MAPK activation, with cantharidin resulting in a far greater TAK1, p38 and JNK activation (Figure S12J). This likely explains why ‘conventional’ and ‘physiological’ TAK1-activating cytokines, including TNFɑ, are not sufficient to cause NLRP1 linker hyperphosphorylation and subsubsequent NLRP1 activation. In light of these results, we refer to the effects of PP1/PP2 inhibition on TAK1 as ‘TAK1 hyperactivation’, to be distinguished from physiological TAK1 activation involved in cytokine signaling.

Taken together, our results establish TAK1 as a selective and key driver of NLRP1 activation induced by PP1/PP2 inhibitors such as cantharidin. This pathway functions independently of, but analogous to, the ribosome-ZAKɑ RSR pathway, both culminating in the hyperphosphorylation of the NLRP1 linker.

### TAK1 directly phosphorylates the NLRP1 linker

Employing a kinase assay with recombinant activated TAK1 (TAK1-TAB fusion) and ZAKɑ, we found that both MAP3Ks can directly phosphorylate SNAP-tagged NLRP1 linker, with at least five phosphorylated species that can be visualized on PhosTag SDS-PAGE (Figure 5A). Importantly, TAK1 and ZAKɑ caused a similar pattern of bandshift that is distinct from p38ɑ (Figure 5A). To narrow down the residues that TAK1 targets, we took a previously established approach by dividing the serine and threonine residues into four regions and performed alanine substitutions en bloc (S->A; T->A, abbreviated as SATA mutants 1-4) (Figure S13A). Each of the NLRP1 SATA mutants were expressed in NLRP1 KO N/TERT cells and compared to wild-type NLRP1 or the STless mutant (where all the serine and threonine residues are mutated to alanine) (Figure 5B and S13B). SATA1 and SATA4 mutants effectively restored canthardin-, dinophysis toxin-, okadaic acid- and anisomycin-triggered pyroptosis to wild-type levels, whereas SATA2 and SATA3 mutants behaved similarly to the STless mutant or vector control (Figure 5B and S13B). Importantly, none of the SATA mutants compromised VbP-induced NLRP1 activation (Figure S13B). These results suggest that certain serine and threonine residues mutated in the SATA2 and SATA3 constructs, ie. between a.a. 115-196, are crucial for both ZAKɑ- and TAK1-driven NLRP1 activation.

**Figure 5:**
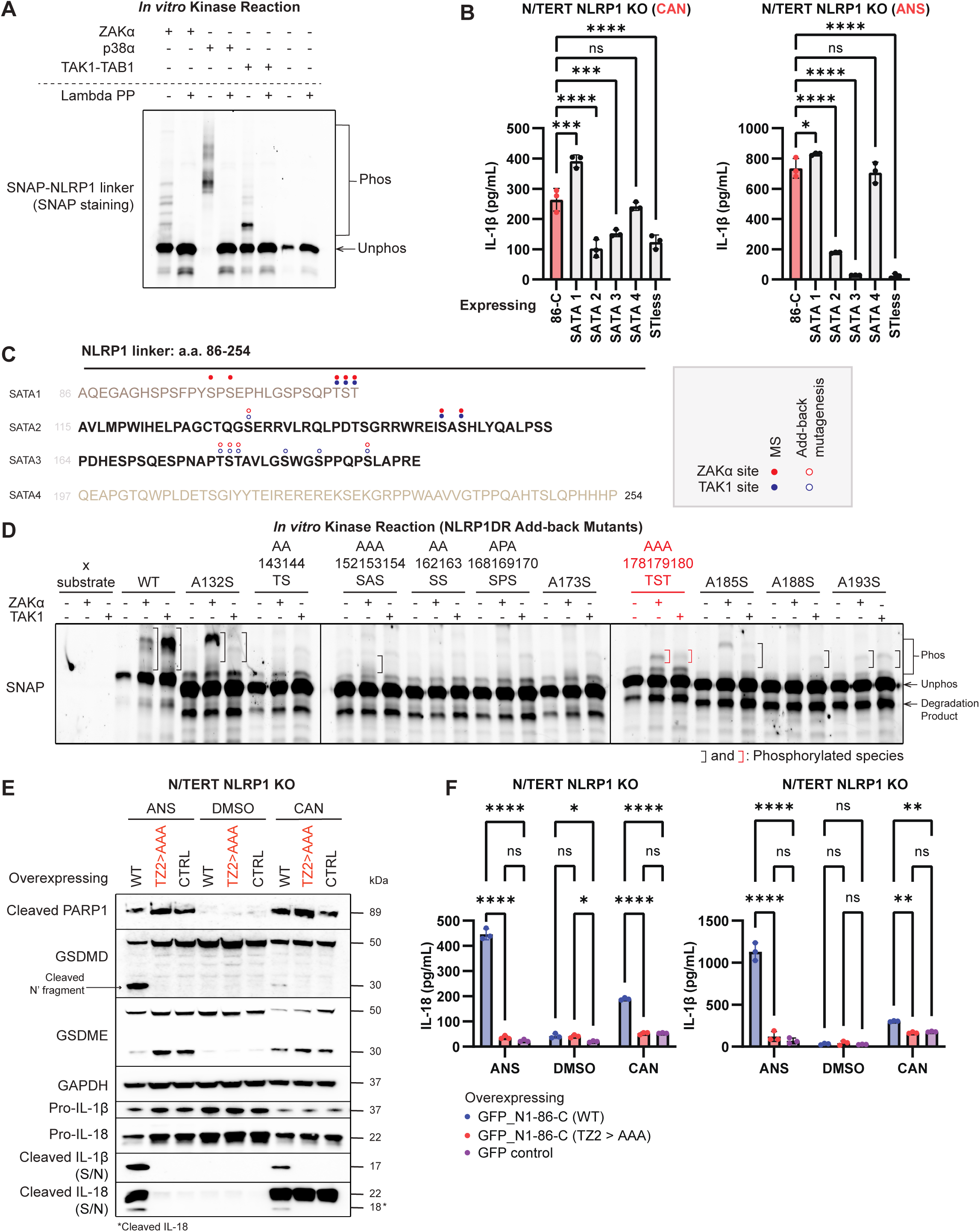
TAK1 directly phosphorylates the NLRP1 DR at specific sites and requires a specific TST motif for NLRP1 inflammasome activation. **A.** Rhodamine labelled recombinant SNAP-tagged NLRP1-DR following PhosTag-SDS-Agarose-PAGE. Recombinant NLRP1-DR was incubated with recombinant ZAKα, p38α, and TAK1-TAB1 fusion in a standard kinase reaction for 60 minutes and labelled with SNAP ligand fluorescence (TMR-STAR). Lambda protein phosphatase or a negative control was spiked into each reaction, as indicated. Phosphorylated vs unphosphorylated species are indicated. **B.** IL-1β ELISA of NLRP1 KO N/TERTs overexpressing GFP_NLRP1-86-C_FLAG (WT), SATA 1-4, or STless mutants following treatment with 50 μM cantharidin or 1 μM anisomycin. Cell culture media was harvested after 8 hours of treatment. **C.** Recombinant SNAP-tagged NLRP1-DR was incubated with recombinant TAK1-TAB1 fusion or ZAKα in a standard kinase reaction for 60 minutes and mass spectrometry was used to detect phosphorylated peptides. Phosphorylated residues detected by mass spectrometry are indicated with solid dots while residues identified from the following add-back analysis are indicated with clear dots. Residues phosphorylated by TAK1 and ZAKα are indicated with blue and red respectively. **D.** Rhodamine labelled SNAP-tagged NLRP1-DR ‘Add-back’ mutants following PhosTag-SDS-Agarose-PAGE. In the STless background, indicated alanine residues were mutated back to serine or threonine. These recombinantly expressed mutants were incubated with recombinant TAK1-TAB1 fusion or ZAKα in a standard *in vitro* kinase reaction for 60 minutes. Phosphorylated vs unphosphorylated species for each protein is indicated. The crucial “TZ2” motif is highlighted in red. **E.** Immunoblot of cleaved PARP1, GSDMD (full length and cleaved), GSDME (full length and cleaved), GAPDH, Pro-IL-1β, cleaved IL-1β, and IL-18 (full length and cleaved) or **F.** IL-18 and IL-1β ELISA from NLRP1 KO N/TERTs overexpressing GFP_NLRP1-86-C_FLAG (WT), TZ2 mutant, or GFP control following treatment with 1 μM anisomycin, DMSO (vehicle control), and 50 μM cantharidin. Cells, floaters and cell culture media were harvested after 8 hours of treatment. Error bars represent SEM from three technical replicates, where one replicate refers to an independent sample. Significance values were calculated based on one-way ANOVA followed by Dunnett’s test for multiple pairwise comparisons (B) or two-way ANOVA followed by Sidak’s test for multiple pairwise comparisons (F). ns, nonsignificant; *P < 0.05; ****P < 0.0001.

We then used three methods to pinpoint the crucial serine and threonines targeted by TAK1. In the first, we subjected the recombinant kinase reaction mixtures to direct mass spectrometry analysis. Strikingly, ZAKɑ and TAK1 phosphorylate the NLRP1 linker at many overlapping sites (Figure 5C). In the second method, we performed targeted mutagenesis of candidate serine/threonine residues in the SATA2-3 region, which had incomplete coverage in the preceding mass spec analysis. Again, many mutants compromised both TAK1- and ZAKɑ mediated phosphorylation of the NLRP1 linker as measured by PhosTag SDS-PAGE (Figure S5C). Next we performed a more stringent ‘add-back’ analysis to differentiate direct phosphosites vs. ‘priming’ sites. In this experiment, each candidate phosphosite was restored to serine or threonine from the STless background. The resulting ‘add-back’ mutants were tested in the recombinant kinase assay with TAK1 and ZAKɑ (Figure 5D). This analysis identified additional phosphosites targeted by both TAK1 and ZAKɑ within the critical a.a. 115-196 region (Figure 5C). Among these, a.a. T178S179T180 is particularly noteworthy, as this site falls within the SATA2 region and has been identified in our previous work as essential (termed ZAKɑ motif #2 therein) for ZAKɑ-driven NLRP1 activation. Our current data now reveal that this motif can also be phosphorylated by TAK1, in the absence of any priming phosphosites. In light of this finding, we rename a.a. T178S179T180 and the surrounding residues TZ (TAK1+ZAKɑ) motif 2. Remarkably, an identical stretch of 7 amino acids present in the SATA1 region is also targeted by both TAK1 and ZAKɑ. This motif is thus renamed as TZ motif 1.

When overexpressed at an identical level relative to wild-type control in NLRP1 KO cells, a NLRP1 mutant bearing alanine substitutions in TZ motif 2 (TZ2>AAA) is defective in both ZAKɑ-induced (ANS, rove beetle extract) and TAK1-induced (cantharidin, dinophysistoxin) activation (Figure 5E-F and Figure S13D-E), as measured by IL-1β/IL-18 secretion and GSDMD cleavage. Thus, TZ motif 2 phosphorylation is the common dominator and converging point between two parallel activation pathways for the NLRP1 inflammasome, which employ two distinct MAP3Ks: TAK1 downstream of PP1/PP2 inhibition and ZAKɑ downstream of ribosome inhibition.

### TAK1 and ZAKα are jointly responsible for dsRNA and CHIKV-driven NLRP1 activation

The extent of TAK1 activation evoked by PP1/PP2 inhibitors, such as cantharidin, is orders of magnitude higher than that of more physiological inflammatory cues, such as TNFɑ (Figure S12J). It thus raises the question whether the TAK1-driven NLRP1 activation pathway uncovered here is an evolutionary edge case, i.e. restricted to isolated toxins, or represents a more general pathogen defense mechanism. Recently, human NLRP1 has been shown to detect a ‘universal’ PAMP: cytosolic long double-stranded RNA (dsRNA) (Bauernfried et al. 2021; Jenster et al. 2023). Employing synthetic dsRNA such as poly(I:C) or viral infection, multiple groups demonstrated that two biochemical events are necessary: physical binding of NLRP1 LRR domain to dsRNA and the activation of ZAKɑ (Jenster et al. 2023; Zhou et al. 2023). Subsequent work showed that the latter is most likely triggered by ribosome stalling caused by widespread mRNA cleavage by RNase L (Xi et al. 2024; Karasik et al. 2024, 2025). Curiously, the requirement of ZAKɑ is partial, as complete ZAKɑ knockout or inhibition does not fully inhibit dsRNA-mediated NLRP1 activation (Jenster et al. 2023; Zhou et al. 2023). Thus, additional regulators are likely involved.

We confirmed that poly(I:C) transfection led to NLRP1-dependent IL-1β secretion as well as p38 phosphorylation and hyperphosphorylation of NLRP1 linker in N/TERT cells in a dose- and time-dependent manner (Figure S14A-C). Neither NLRP1-driven IL-1β/IL-18 secretion (Figure 6A-B) nor NLRP1 linker phosphorylation (Figure 6D) could be fully inhibited by the specific ZAKɑ inhibitor, 6p, even though the auto-phosphorylation of ZAK itself is fully abrogated (Figure 6D). Furthermore, poly(I:C)-induced p38 and JNK phosphorylation are also only marginally diminished by 6p (Figure 6D). This is consistent with the notion that a non-ZAKɑ MAPK mediates dsRNA-triggered NLRP1 and SAPK activation.

**Figure 6:**
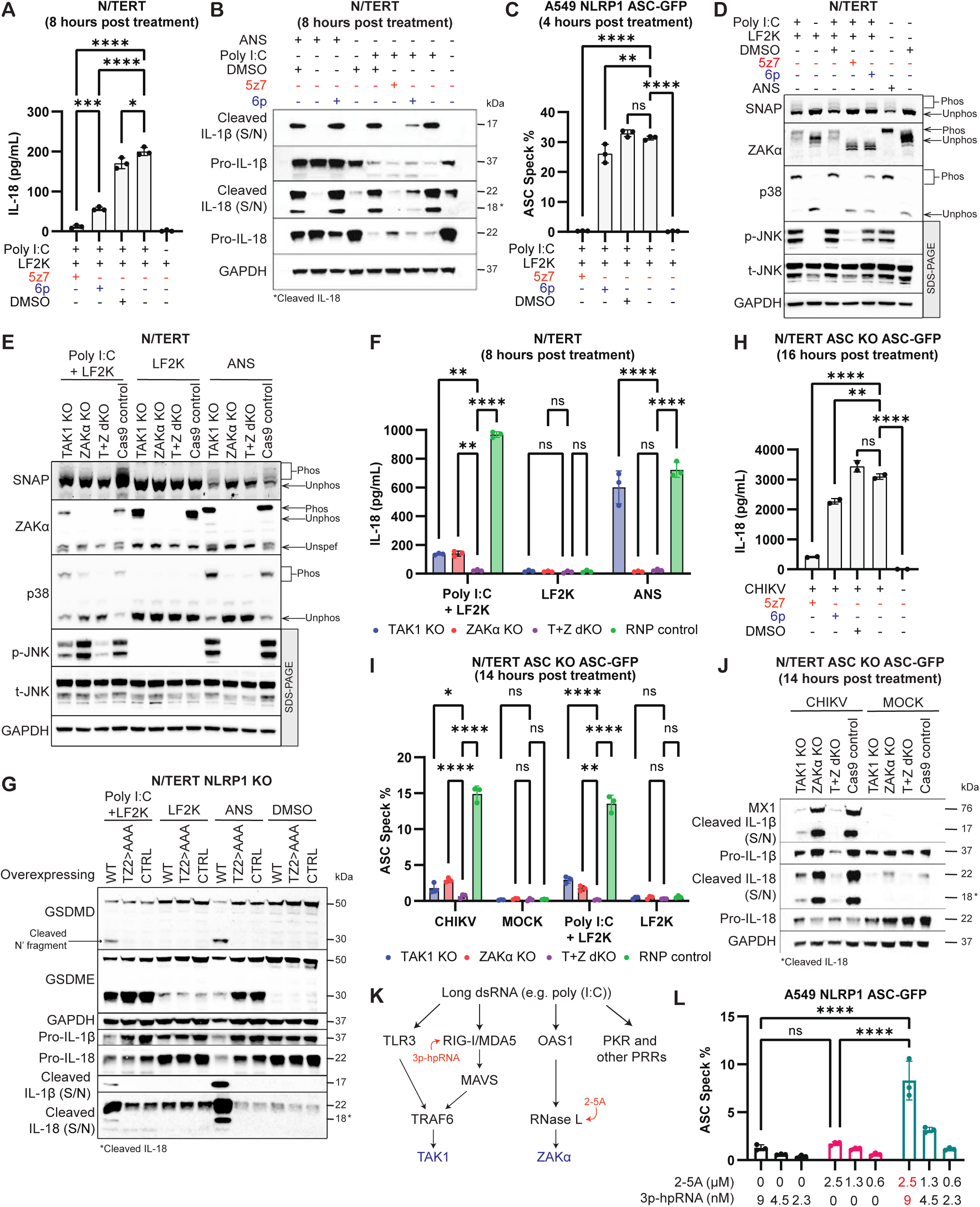
Poly I:C and CHIKV activates the NLRP1 inflammasome via combined actions of TAK1 and ZAKα. **A.** IL-18 ELISA or **B.** Immunoblots of cleaved IL-1β, Pro-IL-1β, IL-18 (full length and cleaved), and GAPDH from N/TERTs pre-treated with 1 μM 5-z7-oxozeanol (pan-kinase inhibitor), 1 μM 6p (ZAK inhibitor), or DMSO (vehicle control) before stimulation with 200ng/mL Poly I:C. Cells and cell culture media were harvested after 8 hours of treatment. **C.** Percentage of ASC-GFP speck forming A549 NLRP1 ASC-GFP reporter cells pre-treated with 1 μM 5-z7-oxoxeanol (pan-kinase inhibitor), 1 μM 6p (ZAK inhibitor), or DMSO (vehicle control) before stimulation with 200 ng/mL Poly I:C. Cells were fixed and stained with DAPI after 4 hours of treatment. **D.** Rhodamine labelled SNAP-tagged NLRP1-DR, immunoblot of ZAK and p38 following PhosTag-SDS-Agarose-PAGE, and immunoblot of p-JNK, t-JNK, and GAPDH following SDS-PAGE from N/TERTs pre-treated with 5 μM emricasan (pan-caspase inhibitor), 1 μM 5-z7-oxoxeanol (pan-kinase inhibitor), 1 μM 6p (ZAK inhibitor), or DMSO (vehicle control) before stimulation with 1 μM anisomycin, DMSO (vehicle control), 200 ng/mL Poly I:C, and LFK2 (transfection control). Cell lysates were harvested after 6 hours of treatment. **E.** Rhodamine labelled SNAP-tagged NLRP1-DR, immunoblot of ZAK and p38 following PhosTag-SDS-Agarose-PAGE, and immunoblot of p-JNK, t-JNK, and GAPDH following SDS-PAGE from TAK1 KO, ZAKα KO, T+Z dKO, and RNP control N/TERTs expressing NLRP1-86-275-SNAP stimulated with 200 ng/mL Poly I:C, LFK2 (transfection control), and 1 μM anisomycin. Cells were harvested after 6 hours of treatment. **F.** IL-18 ELISA from TAK1 KO, ZAKα KO, T+Z dKO, and RNP control N/TERTs following treatment with 200 ng/mL Poly I:C, LFK2 (transfection control), and 1 μM anisomycin. Cell culture media was harvested after 8 hours of treatment. **G.** Immunoblot of GSDMD (full length and cleaved), GSDME (full length and cleaved), GAPDH, cleaved IL-1β, Pro-IL-1β, and IL-18 (full length and cleaved) of NLRP1 KO N/TERTs overexpressing GFP_NLRP1_86-C_FLAG (WT), TZ2 mutant, or GFP control. Cells, floaters and cell culture media were harvested after 8 hours of treatment. **H.** IL-18 ELISA of ASC KO N/TERTs overexpressing ASC-GFP (N/TERT ASC-GFP reporter cells) pre-treated with 1 μM 5-z7-oxozeanol (pan-kinase inhibitor), 1 μM 6p (ZAK inhibitor), or DMSO (vehicle control) before stimulation with CHIKV (MOI 10). Cell culture media was harvested after 16 hours of treatment. **I.** Percentage of ASC-GFP speck forming TAK1 KO, ZAKα KO, T+Z dKO, and RNP control N/TERT ASC-GFP reporter cells following treatment with CHIKV (MOI 10), mock (vehicle control for CHIKV), 200 ng/mL Poly I:C, and LF2K (transfection control). Cells were fixed and stained with DAPI after 14 hours of treatment. **J.** Immunoblot of MX1 (IFN-stimulated gene), Pro-IL-1β, cleaved IL-1β, IL-18 (full length and cleaved), and GAPDH of TAK1 KO, ZAKα KO, T+Z dKO, and RNP control N/TERTs following treatment CHIKV (MOI 10) and mock (vehicle control for CHIKV). Cells and cell culture media were harvested after 14 hours of treatment. **K.** Schematic of known dsRNA sensors and downstream signalling cascades. 3p-hpRNA is a RIG-I agonist while 2-5A activates RNaseL. RIG-I and RNaseL are upstream of TAK1 and ZAKα respectively. **L.** Percentage of ASC-GFP speck forming A549 NLRP1 ASC-GFP reporter cells following transfection of 3p-hpRNA and 2-5A individually or in combination. Cells were fixed and stained with DAPI after 3 hours of treatment. Error bars represent SEM from three technical replicates, where one replicate refers to an independent sample. Significance values were calculated based on one-way ANOVA followed by Dunnett’s test (A, C, and H) or two-way ANOVA followed by Sidak’s test for multiple pairwise comparisons (F, I, and L). ns, nonsignificant; *P < 0.05; ****P < 0.0001.

Additional investigation confirms that TAK1 is the missing link. First, dual inhibition of TAK1 and ZAKɑ by 5z7 completely abrogated poly(I:C) induced pyroptosis in N/TERT cells (Figure 6A-B and S14D) and the formation of ASC-GFP specks in an A549 NLRP1-ASC-GFP reporter cell line (Figure 6C). Further, we examined NLRP1 linker phosphorylation in poly(I:C) transfected ZAKɑ KO, TAK1 KO and T+Z dKO N/TERT cells (Figure 6E and S14F). In agreement with the results obtained using 5z7, poly(I:C)-driven NLRP1 linker phosphorylation was partially reduced in single KO cells and was fully abrogated in T+Z dKO cells (Figure 6D-E). A similar trend was observed for p38 and JNK phosphorylation (Figure 6E), whose activation was fully suppressed in T+Z dKOs, but not in single KOs. These results demonstrate that ZAKɑ and TAK1 are jointly responsible for poly(I:C)-triggered NLRP1 linker phosphorylation and SAPK activation. As constitutive TAK1 KO (and T+Z dKO) reduced pro-IL-1β levels, IL-1β ELISA and cleavage is not a faithful readout to measure pyroptosis in these cells. Thus, we opted to examine IL-18 secretion and cleavage in subsequent experiments employing TAK1 KO, as pro-IL-18 is not under the control of TAK1 signaling. We further generated an analogous series of KO cells where the endogenous ASC locus was replaced with ASC-GFP (Figure S14E). Simultaneous removal of both TAK1 and ZAKɑ is required to fully eliminate poly(I:C)-driven IL-18 cleavage/secretion, ASC-GFP speck formation and GSDMD cleavage, which were only partially reduced in single KO cells (Figure 6F and Figure S14G-K). Mutating the shared TZ motif 2 also completely abrogated poly(I:C)-driven NLRP1 activation, phenocopying TAK1 and ZAKɑ dKO (Figure 6G and S14L-M). These results further corroborate the joint requirement of both TAK1 and ZAKɑ. Interestingly, the level of GSDME and caspase-3 cleavage is not consistently reduced in any of the KO cells or by the mutation of TZ motif 2, indicating that the role of TAK1 and ZAKɑ in poly(I:C) induced cell death is restricted pyroptosis, but not apoptosis (Figure 6G and S14K).

We then examined NLRP1 inflammasome activation during chikungunya virus (CHIKV) infection. CHIKV is a mosquito-borne positive-sense, single-stranded RNA (ssRNA) virus. Although keratinocytes are typically not considered a major player in viral dissemination in vivo, human keratinocytes, including N/TERT cells, are permissive for CHIKV and likely experience frequent viral challenge in the real world through mosquito bites. Replication-associated dsRNA intermediates have previously been shown to activate the NLRP1 inflammasome in a partially ZAKɑ-dependent manner, similar to poly(I:C) (Jenster et al. 2023). Using IL-1β/IL-18 secretion, ASC speck formation and GSDMD cleavage as proxies for inflammasome activation, we confirmed that CHIKV caused MOI-dependent NLRP1 activation in N/TERT cells (Figure S15A-E). All three inflammasome readouts were fully abrogated by the 5z7, but only partially reduced by the ZAKɑ-specific inhibitor 6p (Figure 6H and Figure S15F-H). Similar to intracellular poly(I:C), CHIKV-induced IL-18 cleavage/secretion and ASC speck formation were most affected by double knockout of TAK1 and ZAKɑ (TZ dKO) relative to control cells, but only partially reduced in single KOs (Figure 6I-J and S15I). Mutating the TZ motif 2 fully abrogated CHIKV-induced IL-1β/IL-18 secretion (Figure S15J). Thus, just like poly(I:C), CHIKV also requires both TAK1 and ZAKɑ to activate the NLRP1 inflammasome in N/TERT cells.

We also performed a complementary experiment to test the potential synergy between ZAKɑ and TAK1 in the context of NLRP1 activation. Many poly(I:C)-responsive PRRs require TAK1 and/or ZAKɑ for downstream signaling. For instance, TAK1 can be activated by RIG-I (Mikkelsen et al. 2009), MDA5 (Rui et al. 2017) and TLR3 (Jiang et al. 2004), whereas ZAKɑ can be activated by RNase L (Figure 6K) (Xi et al. 2024; Karasik et al. 2024). We thus used specific PRR ligands to more precisely control TAK1 and ZAKɑ activation. In A549-NLRP1-GFP reporter cells, combined intracellular delivery of the RIG-I ligand, 3p-hpRNA and RNase L ligand, 2-5A led to a prominent increase in ASC-GFP specks, but each ligand alone had minimal effect (Figure 6L and Figure S14N). These results further corroborate the loss-of-function experiments and demonstrate that TAK1 and ZAKɑ synergistically drive dsRNA-mediated NLRP1 activation. In this reductive system, the maximum level of ASC-GFP specks achieved was significantly less than poly(I:C) itself. This can be explained by the fact that direct binding of NLRP1 LRR to long dsRNA, which is not modeled by 3p-hpRNA, is also necessary to unleash full-scale NLRP1 activation (Bauernfried et al. 2021). Additional TLR3 activation might also be required to attain a high level of TAK1 activation that is commensurate with poly(I:C).

## DISCUSSION

In this study, we investigated the effects of two dermatitis-causing pests, blistering beetles and rove beetles, on human keratinocytes and *in vitro* 3D reconstructed skin cultures. Our results demonstrate that crude extracts from both beetle species can induce rapid pyroptotic death of human skin keratinocytes via the NLRP1 inflammasome, with each beetle species requiring a distinct upstream MAP3K. Pederin/pseudopederin in rove beetles acts as a potent ribosome inhibitor, activating NLRP1 via the RSR kinase ZAKɑ. Cantharidin in blister beetles inhibits PP1/PP2 phosphatase complexes and activates the NLRP1 inflammasome via TAK1. These results provide plausible mechanisms of pathogenesis for both forms of beetle-associated dermatitis, which have been underexplored. *Paederus* dermatitis caused by rove beetles, in particular, has been increasingly reported in China and Southeast Asia, posing significant threats to human health (Zhang 2023; Wang 2025; Wu et al. 2025; The Star Online 2024; Nga 2022; Shaw 2020; Khalil 2022). Reports of blister beetle dermatitis are less frequent, likely due to the restricted distribution of blister beetle species. Nonetheless, our characterization sheds light on the efficacy of cantharidin as a wart removal agent. We propose that the same mechanism, i.e. TAK1-driven NLRP1 activation, might also account for the well-known, but unexplained skin blistering properties of other PP1/PP2 targeting environmental toxins, particularly those found in the aquatic environment, such as okadaic acid, dinophysistoxin, calyculin A and microcystin. Topical inhibition of the NLRP1 inflammasome, or IL-1β/IL-18-driven inflammation, might help alleviate reported disease symptoms.

Mechanistically, our study establishes that TAK1 and ZAKɑ both directly phosphorylate the NLRP1 disordered linker region on multiple overlapping, but non-identical sites. At least one of the shared sites (TZ motif 2) is essential for both ZAKɑ- and TAK1-driven NLRP1 activation. This repositions human NLRP1 as an innate immune sensor for over-stimulation of stress-responsive MAPK signaling. We speculate that additional MAP3Ks, other than TAK1 and ZAKɑ could function as upstream triggers for the NLRP1 inflammasome. Future studies will be required to test this conjecture in various stress and/or infection contexts.

Our study also underscores the value of uncommon natural immunostimulatory molecules as probes to dissect human innate immune signaling. Similar to how rare monogenic disorders can illuminate the pathogenesis of common diseases through stronger phenotypes and more direct genotype–phenotype links, we postulated that natural PP1/PP2-inhibiting toxins such as cantharidin represent an ‘edge case’ of a broader and more general pathogen defense pathway in which TAK1 acts as a critical switch for the NLRP1 inflammasome. Indeed, we discovered a role for TAK1 in dsRNA-induced NLRP1 activation. In this context, TAK1 and ZAKɑ jointly drive NLRP1 activation, and neither kinase can substitute for the other. This is supported by experiments showing that dsRNA-induced NLRP1 activation is only fully abrogated by the genetic deletion or pharmacologic inhibition of both MAP3Ks, and only partially reduced in single KO cells. Based on these data and recent reports (Bauernfried et al. 2021; Jenster et al. 2023), we propose that dsRNA-mediated NLRP1 activation requires distinct signals: 1) phosphorylation of TZ motifs by TAK1 and ZAKɑ, and 2) dsRNA binding to the LRR domain.

Why does dsRNA require both TAK1 and ZAKɑ, whereas ribotoxins and the PP1/PP2 inhibitors require only one MAP3K? We speculate this is likely because dsRNA causes much weaker MAP3K activation, and the accompanying NLRP1 phosphorylation, than ribotoxic stress agents, such as rove beetle extract, UV, anisomycin and PP1/PP2 inhibitors such as cantharidin and okadaic acid (Figure 6). However, intracellular dsRNA has the unique ability of triggering multiple PRRs and activating both TAK1 and ZAKɑ simultaneously, which can then act additively and/or synergistically to achieve sufficient phosphorylation for the NLRP1 linker region. If proven, these results also imply that the NLRP1 functions as a ‘signal integrator’ that can compute the strength of the MAP3K input. In other words, NLRP1 activation will ensue as long as a certain threshold level of phosphorylation of the linker region has been reached, which could be caused by the hyperactivation of a single MAP3K (as in the case of ribotoxins and PP1/PP2 inhibitors) or weaker activation of two MAP3Ks (e.g. dsRNA). It is plausible that the phosphorylation of multiple sites in the linker region induces a progressive conformational change or subtly decreases the stability of the NLRP1 N-terminal fragment to relieve its repressive effect on the C-terminal fragment. Future studies are needed to test these hypotheses.

In conclusion, by investigating two poisonous beetle species, our study expands the repertoire of NLRP1 activators and uncovers a TAK1-driven pathway that activates the human NLRP1 inflammasome in parallel to ZAKɑ-driven RSR. Furthermore, we demonstrate the involvement of both pathways in dsRNA-triggered, NLRP1-mediated pyroptosis. These discoveries will likely find applications in the treatment of beetle-associated dermatitis and viral infections, as well as the design of mRNA therapeutics that can better evade adverse reactions triggered by dsRNA. However, there are certain limitations and caveats to our results:

1. Despite numerous attempts, we were not able to obtain pure pederin from commercial or academic sources. Thus, the attribution of rove beetles’ ability to inhibit the ribosome and activate NLRP1 to pederin/pseudopederin was based on correlation to relative concentrations of pederin/pseudopederin. Future experiments with purified pederin will be required to fully validate these results.
2. Our experiments with blister beetles and cantharidin imply that endogenous TAK1 signaling is constitutively suppressed by PP1 and PP2. The exact molecular mechanisms are beyond the scope of this study. As PP1/PP2 complexes consist of many adaptor proteins, it would be interesting to identify the dedicated adaptor that connects TAK1 to PP1/PP2. Furthermore, none of the PP1/PP2 targeting toxins used in this study, including cantharidin, strongly differentiates PP1 vs. PP2 complexes. Therefore, it is not clear which complex plays a predominant role in regulating the TAK1-NLRP1 axis.
3. The NLRP1 linker region is not conserved in laboratory rodents. In addition, mouse skin, in stark contrast to human skin, does not express components of the inflammasome pathway (Sand et al. 2018). Thus, it is not feasible to model the effect of blistering beetles and rove beetles in mice. Other non-standard animal species, such as primates, might be required to study the role of NLRP1 inflammasome in beetle-associated dermatitis and other skin disorders.
4. Even though our study identifies additional chemical activators and genetic regulators for the NLRP1 inflammasome, we have not fully addressed a key question: how does phosphorylation lead to NLRP1 activation? To answer this question thoroughly will most likely require substantially more work that goes beyond the scope of this study. Nonetheless, the experiments presented here can provide certain enabling tools. For instance, it should be possible to reconstitute NLRP1 phosphorylation and activation *in vitro* using recombinant TAK1 and/or ZAKɑ and dsRNA. This would enable future structural studies to capture the phosphorylated, ligand-bound transitional states of NLRP1.

## ACKNOWLEDGEMENTS

The authors would like to acknowledge Dr. ZHENG Yu-xin (Zhejiang University, China), Dr. YANG Liang (SUSTech, China) and numerous helpful strangers on RedNote, WeChat and Instagram who answered our appeal and helped us collect rove beetles from all across China. We also thank Calvin LEUNG (ASE, NTU, Singapore), Sean YAP (CNCS, NUS, Singapore) and James KHOO (The Curious Pangolin) for sharing their knowledge and passion about beetles. The authors dedicate this study to colleagues devoted to fundamental, non-applied research, whose work, though not always yielding immediate economic returns, is vital to advancing our understanding of the natural world and human health. The authors acknowledge funding from the National Medical Research Council, Singapore (MOH-001499), Ministry of Education Tier 1 grant (RT23/23), and Ministry of Education Tier 2 grant (MOE-T2EP30222-0008). This work was supported by Asian Skin Microbiome Programme 2.0 IAF-PP grant (H22J1a0040) and Atopic Dermatitis Research Programme for Patients (ADEPT) project (NMRC OF-LCG: MOH001635).

## MATERIALS AND METHODS

### Study design

The aim of this study was to understand how phosphorylation activates the NLRP1 inflammasome. A naturally occurring beetle toxin, Pederin, was used to treat epithelial cells and found to activate the NLRP1 inflammasome in a ZAKα-dependent manner. Cantharidin, a naturally occurring phosphatase inhibitor secreted by beetles, was used to identify that an alternative MAP3K, TAK1, is sufficient and necessary to phosphorylate and activate the NLRP1 inflammasome. In addition, genetic knockouts demonstrated a division of labour between TAK1 and ZAKα to phosphorylate and activate NLRP1 in response to dsRNA and viral sensing.

### Cell culture

293Ts (ATCC #CRL-3216) and A549s (InvivoGen a549-ascg-nlrp1) were cultured according to manufacturer’s protocols. Immortalized human keratinocytes (N/TERT) were provided by H. Rheinwald (MTA) and cultured in Keratinocyte Serum-Free Media (Gibco, 17005042) supplemented with final concentration of 25 μg/ml of bovine pituitary extract (Gibco, 13028-014),

294.4 pg/ml of recombinant human epidermal growth factor (Gibco, 10450-013), and 300 μM Calcium Chloride (Kanto Chemicals, 07058-00). Primary human keratinocytes were derived from the skin of healthy donors and obtained with informed consent from the Asian Skin Biobank (ASB) (https://www.a-star.edu.sg/sris/technology-platforms/asian-skin-biobank). All cell lines underwent routine mycoplasma testing with MycoGuard (Genecopoeia #LT07-118).

### CRISPR-Cas9 or lentiviral knockout cell lines

HEK293T-*NLRP1-ASC-GFP*, N/TERT *ASC* KO, N/TERT *ASC* KO overexpressing *ASC-GFP*, N/TERT *NLRP1* KO, N/TERT *p38α/β* dKO, and N/TERT *ZAKα* KO cells were previously described <u>(Robinson et al. 2022)</u>. Lentiviral Cas9 and guide RNA plasmid (LentiCRISPR-V2, Addgene plasmid #52961) or RNP (ribonucleoprotein) CRISPR electroporation was used to create stable deletions in N/TERT keratinocytes.

The sgRNAs target sequences (5′ to 3′) for N/TERT keratinocytes are: *NLRP1* (GATAGCCCGAGTGCATCGG), *ASC* (GCTAACGTGCTGCGCGACAT), *MAP3K20* (*ZAKα*) sg1 (TGTATGGTTATGGAACCGAG), *MAP3K20* (*ZAKα*) sg4 (TGCATGGACCGGAAGACGATG), *MAPK14* (*p38α*) sg1 (TGATGAAATGACAGGCTACG), *MAPK11* (*p38β*) sg1 (CGACGAGCACGTTCAATTCC), *MAPK11* (*p38β*) sg2 (ACTCGGCCGGGATCATCCAC), *MAP3K7* (*TAK1*) sg1 (CTCACCGGCCGAAGACGAGG), *MAP3K7* (*TAK1*) sg2 (GATCGACTACAAGGAGATCG), and Cas9 control (GGATTATATCCGGAAGACCC).

*TAK1* or *ZAKα* KO N/TERTs were generated by nucleofecting with CRISPR/ Cas9 RNPs with crRNAs encoding *MAP3K7* (*TAK1*) sg1 and sg2 or *MAP3K20* (*ZAKα*) sg1 and sg4 respectively. The detailed protocol for RNP assembly and nucleofection were described in detail previously. The same sgRNA target sequences, sg1 and sg2, were used to generate *TAK1*-KO primary keratinocytes. Knockout efficiency was validated by immunoblot and/ or sanger sequencing of extracted genomic DNA. Overall Cas9 editing efficiency was determined using the Synthego ICE tool (Synthego Performance Analysis, ICE Analysis. 2019. v2.0. Synthego, https://ice.synthego.com/#/). *TAK1 + ZAK* double knockout (*T+Z* dKO) N/TERTs were generated by nucleofecting *ZAK* KO N/TERTs with CRISPR/ Cas9 RNPs with crRNAs encoding *MAPK7* (*TAK1*) sg1 and sg2.

### Plasmids and preparation of lentiviral stocks

Constitutive lentiviral expression was performed using pCDH vector constructs (System Biosciences) and packaged using third-generation packaging plasmids. Primers (IDT Technologies) were designed for site-directed mutagenesis to generate *NLRP1* mutants. Mutagenesis was performed in pCS2+_NLRP1-FL plasmids before the cloning of residues 86-1473 a.a. (referred to as 86-C) into pCDH vector constructs, between an N’ terminal GFP and a C’ terminal FLAG tags, for lentiviral expression. All expression plasmids for transient expression were cloned into the pCS2+ vector backbone using InFusion HD (Clontech) or 2X CE (Vazyme). *TAK1* plasmids were obtained from GenScript and ZAKα plasmids were gifts from S. Bekker-Jensen (Addgene plasmids #141193, #141194, #141195, #141196, and #141197; http://n2t.net/addgene:141193 ; RRID:Addgene_141193).

### Drugs or chemicals

List of drugs and chemicals used in this study can be found in Supplementary Table 1. All drugs and chemicals were dissolved and stored according to manufacturer’s protocols.

### Beetle Extract

Beetles were obtained from several sources. To obtain beetle extract, beetles of the *Meloida* family were crushed in a mortar and pestle until a fine powder was achieved. The fine powder was resuspended analytical grade methanol (Sigma-Aldrich, 67-56-1) with 1% hydrochloric acid (Sigma-Aldrich, 7647-01-0). Suspension was incubated at room temperature overnight and filtered through a 70 μm cell strainer before evaporation to dryness. Dried beetle extract was reconstituted in analytical grade methanol and filtered through a 0.22 μm filter before treatment of cells. Methanol was used as a vehicle control in these experiments.

To obtain beetle extracts, beetles of the *Paederus* family were crushed in a mortar and pestle until a fine powder was achieved. The fine powder was resuspended in analytical grade ethanol (Sigma-Aldrich, 64-17-5). Suspension was incubated at 37 °C, 200 rpm for 4 hours. After incubation, suspension was centrifuged at 21,000g for 5 minutes to pellet beetle debris. Supernatant was transferred to a clean Eppendorf tube and covered with parafilm. Holes were punctured in the parafilm cover before lyophilization overnight at 30 °C, 1400 rpm. Lyophilized beetle extract was resuspended in 50 μL analytical grade ethanol. Ethanol was used as a vehicle control in these experiments.

### Transfection of Poly (I:C), 3p-hp RNA, and compound 2-5A

Cells were transfected with HMW Poly(I:C) (InvivoGen), 3php (InvivoGen), and compound 2-5A using Lipofectamine 2000 (Thermo Fisher Scientific, 11668019) diluted in Opti-MEM. Cell culture media was used to top up the final volume to 500 μL. Cells were then incubated for various durations before ASC-GFP specks were quantified or cell lysates and cell culture media were harvested. Compound 2-5A was a gift from S. Bekker Jensen (University of Copenhagen).

### CHIKV Generation and infection

Chikungunya viruses were a gift from Dahai Lab (Nanyang Technological University, Lee Kong Chian School of Medicine). Viruses were propagated in Vero cells (ATCC, CCL-81) cultured in DMEM (Gibco, 11995065), supplemented with 10% FBS (Gibco, 16140071). Plaque assays were done to determine MOI of propagated viruses. Viruses were stored at -80 °C and freeze-thaw cycles were avoided. DMEM supplemented with 10% FBS was used as mock/vehicle control in these experiments.

Cells were infected with viruses by incubating a thin layer of viruses diluted in Opti-MEM™ (Gibco, 31985070) at 37 °C, for 1 hour. Gentle agitation of plate was required every 10 minutes for more efficient transduction of viruses. Viral media was discarded and fresh cell culture media was added. Cells were incubated for 16 hours before cell lysates and cell culture media were harvested. All viral waste was disposed in 10% bleach.

### Analysis of Pederin and Pseudopederin

Beetle extracts were diluted 100x in methanol containing 300 nM verapamil as internal standard and used for UHPLC-MS/MS analysis. The system consisted of a Vanquish™ Duo UHPLC, coupled to TSQ Quantis™ Triple Quadrupole Mass Spectrometer with a heated electrospray ionization (H-ESI) source, operated in positive ion mode. Chromatographic separation was achieved using a Waters UPLC^®^ BEH C18 1.7 μm (2.1 x 50 mm) column protected by a guard column. The column temperature was maintained at 40 °C. Mobile phase A was 0.1% formic acid in water, while mobile phase B was 0.1% formic acid in methanol. The flow rate was maintained at 0.2 mL/min throughout the run. The gradient program was as follows: 50% B at 0 min, linearly increased to 85% B from 0 to 5 min, ramped to 100% B at 5.1 min, held until 7.0 min, then returned to 50% B at 7.1 min and re-equilibrated until 11.0 min. The injection volume was 1 μL. MS/MS data were acquired in multiple reaction monitoring (MRM) mode. Spray voltage was set at 3.5 kV, ion transfer tube temperature at 325 °C, vaporizer temperature at 230

°C, sheath gas at 50 (arb), auxiliary gas 18 (arb), and sweep gas at 0 (arb). Data acquisition and processing were carried out using Thermo Xcalibur™ software (version 4.2.47) and Skyline software (version 25.1.0.142) was used for analysis and quantification. The MRM table for the detection of pederin, pseudopederin and verapamil can be found in Supplementary Table 3.

### Standard Addition for the Analysis of Cantharidin and Cantharidic Acid

The beetle extract was diluted 200x in methanol. Aliquots of the beetle extract (20 μL) were taken into separate tubes and spiked with either cantharidin or cantharidic acid standards (80

μL). For no matrix control, methanol (20 μL) was taken into separate tubes and spiked with the same cantharidin or cantharidic acid standards (80 μL). The final concentrations of the cantharidin series were 0, 1, 2, 4, 8 and 10 μM. The final concentrations of the cantharidic acid series were 0, 4, 8, 16, 32 and 40 μM. All samples were subjected to UHPLC-MS/MS analysis. The system consisted of a Vanquish™ Duo UHPLC, coupled to TSQ Quantis™ Triple Quadrupole Mass Spectrometer with a heated electrospray ionization (H-ESI) source, operated in positive ion mode. Chromatographic separation was achieved using a Waters UPLC^®^ BEH C18 1.7 μm (2.1 x 50 mm) column protected by a guard column. The column temperature was maintained at 40 °C. Mobile phase A was 0.1% formic acid in water, while mobile phase B was 0.1% formic acid in methanol. The flow rate was maintained at 0.2 mL/min throughout the run. The gradient program was as follows: 2.5% B at 0 min, increased to 90% B from 0 to 5.5 min (curve 7), held until 7.4 min, then returned to 2.5% B at 7.5 min and re-equilibrated until 13.5 min. The injection volume was 1 μL. MS/MS data were acquired in multiple reaction monitoring (MRM) mode. Spray voltage was set at 3.5 kV, ion transfer tube temperature at 325 °C, vaporizer temperature at 230 °C, sheath gas at 50 (arb), auxiliary gas 18 (arb), and sweep gas at 0 (arb). Data acquisition and processing were carried out using Thermo Xcalibur™ software (version 4.2.47) and Skyline software (version 25.1.0.142) was used for analysis and quantification. The MRM table for the detection of cantharidin and cantharidic acid can be found in Supplementary Table 4.

### Immunoblotting

For SDS-PAGE using whole cell lysates, cells and floaters were separately harvested in 1X Laemmli buffer, heated at 95 °C for 5 min, and loaded into SDS-PAGE gels to run. For analysis of GSDMD and GSDME by immunoblotting, cells and floaters were combined. For analysis of secreted IL-1β and IL-18 cytokines by immunoblotting, harvested cell culture media was concentrated using filtered centrifugation (Merck, Amicon Ultra, #UFC500396). Protein samples were run on SurePAGE™, Bis-Tris, 10x8, 4-20% gels (Genscript, M00656/7) and blotted onto 0.22 μm Nitrocellulose membranes (Bio-Rad, 1620112). Membranes were blocked with 3% milk (Biobasic, NB0669) at room temperature for 1 hour and incubated with primary antibodies at 4°C overnight with gentle agitation. Corresponding secondary antibodies were then added at room temperature for 1 hour with gentle agitation. All primary and secondary antibodies used in this study are described in Supplementary Table 2. Various enhanced chemiluminescent (ECL) substrates were used and blots were visualized using a ChemiDoc Imaging system (Bio-Rad).

For Phos-tag analysis using whole cell lysates, cells were directly harvested in 1X Laemmli buffer, heated at 95 °C for 5 minutes, and loaded into Phos-tag gels to run. Phos-tag SDS-PAGE was carried out using homemade 8% SDS-PAGE gel, with addition of Phos-tag Acrylamide (Wako Chemicals, AAL-107) to a final concentration of 15 μM or 30 μM and manganese chloride(II) (Sigma-Aldrich, #63535) to a final concentration of 30 μM or 60 μM. Once the run was completed, Phostag gels were washed twice in 1X transfer buffer containing 10 mM EDTA. Subsequently, the gel was washed once in 1X transfer buffer without EDTA and immunoblotting was carried out as described above.

### Cytokine analysis

To quantify secreted IL-1β and IL-18 cytokines, human IL-1β enzyme linked immunosorbent assay (ELISA) kit (BD, #557953) and human IL-18 ELISA (R&D Systems, DY318-05) were used according to manufacturer’s protocols. Cell culture media from various experiments were diluted appropriately and used for ELISA.

### AHA/ OPP Assays

A549 NLRP1 ASC-GFP reporter cells were seeded at a cell density of 8K cells/well in a black 96-well plate (PerkinElmer, CellCarrier-96 Ultra, #6055300). Cells were treated in methionine-free cell culture media. To quantify translation inhibition, nascent polypeptides were labelled with 100 μM Click-IT™ AHA (L-Azidohomoalanine) (Vector Laboratories, CCT-1066) over the course of the treatment duration. Cells were fixed with 4% PFA and permeabilized with 0.3% Triton-X. Incorporated AHA was labelled with 2 μM AZDye™ 568 Alkyne (Vector Laboratories, CCT-1294). Cells were stained with 300 nM DAPI.

Images of cells were acquired in five random fields with 4′,6-diaminidino-2-phenylindole (DAPI, 358 nm/461 nm), GFP (469 nm/525 nm), AZDye (578 nm/602 nm), and brightfield channels under 10X magnification using the Operetta high content screening microscope (Perkin Elmer Operetta CLS imaging system, NTU Optical Bio-Imaging Centre in Nanyang Technological University, Singapore). Images were stored and analysed using the Harmony software (Version 6). For five fields of view per well, with three wells per treatment, the mean intensity of 568 nm fluorescence of cells was measured. Mean intensity was then calculated as a percentage of the mean intensity measured from vehicle control treated but stained cells, this is the percentage of AHA incorporation/ vehicle control as a proxy for translation inhibition.

### Puromycin incorporation and Harringtonine Run-off Assays

To visualise translation inhibition, treated cells were pulsed with 10 μg/mL puromycin for 10 minutes before harvest. In Harringtonine Run-off Assays, cells were treated with 2 μg/mL Harringtonine (translating initiation inhibitor) before run-off times of 10, 5, and 0 minutes. Cells were then pulsed with 10 μg/mL puromycin for 10 minutes before harvest. Cells were harvested in 1X Laemmli buffer, heated at 95 °C for 5 min, and loaded into SDS-PAGE gels to run. Immunoblotting with an anti-puromycin antibody was used to visualize puromycin-labelled actively translating peptides at the time of puromycin pulse.

### DRAQ7 inclusion assay

N/TERT cells were seeded at a cell density of 10K cells/well in a black 96-well plate (PerkinElmer, CellCarrier-96 Ultra, #6055300). For certain experiments, cells were seeded at a cell density of 80K cells/well in black 24-well plates (Cellvis, P24-1.5P). The next day, cells were treated and stained with a final concentration of 0.3 μM DRAQ7™ Dye (Invitrogen, D15106) or Propidium iodide ().

Images of cells were acquired in five random fields with DRAQ5 (647 nm/681 nm), and brightfield channels at 10X magnification using the Operetta high-content screening microscope (Perkin Elmer Operetta CLS imaging system, NTU Optical Bio-Imaging Centre in Nanyang Technological University, Singapore). Images were taken over 14-18 hours, at 20 minute intervals. Images were stored and analysed using the Harmony software (Version 6). For five fields of view per well, with three wells per treatment, the number of live cells was counted using digital phase contrast images which can identify cell borders. The number of DRAQ7-stained cells, identified through the DRAQ7 channel (488 nm/647 nm), were counted as DRAQ7 positive cells. The number of DRAQ7 positive cells was then calculated as a percentage of the total number of cells, this is the percentage of cell death for a given timepoint. Percentage of cell death was plotted against the time course.

### Overexpression of Kinase cDNA in HEK239T NLRP1 ASC-GFP Reporter Cells

HEK293T NLRP1 ASC-GFP reporter cells were seeded at a cell density of 80K cells/well in a 24-well plate format. Cells were transfected with 500 ng/mL of cDNA encoding each kinase (ZAKα, TAK1, and p38α) using Lipofectamine 2000 (Thermo Fisher Scientific, 11668019) diluted in Opti-MEM. Cells were incubated for 24 hours. Cells were fixed in 4% PFA, permeabilized with 0.3% Triton-X, and stained with 300 nM DAPI.

Images of cells were acquired in eight random fields with 4′,6-diaminidino-2-phenylindole (DAPI, 358 nm/461 nm), GFP (469 nm/525 nm), and brightfield channels under 20X magnification using the Operetta high content screening microscope (Perkin Elmer Operetta CLS imaging system, NTU Optical Bio-Imaging Centre in Nanyang Technological University, Singapore). Images were stored and analysed using the Harmony software (Version 6). For five fields of view per well, with three wells per treatment, the number of ASC-GFP speck forming cells was calculated. The number of live cells was identified through the DAPI channel and the number of ASC-GFP speck forming cells was identified through the GFP channel (488 nm/647 nm). The number of ASC-GFP speck forming cells was then calculated as a percentage of the total number of cells, this is the percentage of ASC-GFP speck forming cells.

### ASC-GFP Speck Assays

A549 NLRP1 ASC-GFP reporter cells were primed with 10 ng/mL TNFα overnight before treatment. Treated cells were fixed in 4% PFA, permeabilized with 0.3% Triton-X, and stained with 300 nM DAPI. Cells were imaged on the Operetta high content screening microscope (Perkin Elmer Operetta CLS imaging system, NTU Optical Bio-Imaging Centre in Nanyang Technological University, Singapore). For five fields of view per well, with three wells per treatment, the number of ASC-GFP speck forming cells was calculated. The number of live cells was identified through the DAPI channel and the number of ASC-GFP speck forming cells was identified through the GFP channel (488 nm/647 nm). The number of ASC-GFP speck forming cells was then calculated as a percentage of the total number of cells, this is the percentage of ASC-GFP speck forming cells.

### Organotypic 3D reconstructed skin culture

Fibroblast layer was generated with 2 mL collagen I (Corning, #354249) with 7.5 × 10^5^ human fibroblasts and polymerized over 1 mL of acellular collagen I in six-well culture inserts (Falcon, #353102) placed in 6-well deep well plates (Falcon, #355467). After 24 hours, 1 × 10^6^ primary human keratinocytes were seeded into the inserts and kept submerged in a 3:1 DMEM (Hyclone, #SH30243.01) and F12 (Gibco, #31765035) mixture supplemented with 10% FBS (Hyclone, #SV30160.03), 100 U/mL of penicillin–streptomycin (Gibco, #15140122), 10 µM Y-27632 (Tocris, #1254), 10 ng/mL of EGF (Sigma-Aldrich, #E9644), 100 pM cholera toxin (Enzo, #BML-G117-001), 0.4 µg/mL of hydrocortisone (Sigma-Aldrich, #H0888), 0.0243 mg/mL adenine (Sigma-Aldrich, #A2786), 5 µg/mL of insulin (Sigma-Aldrich, #I2643), 5 µg/mL of transferrin (Sigma-Aldrich, #T2036), and 2 nM 3,3′,5′-triiodo-l-thyronine (Sigma-Aldrich, #T6397). After 24 hours, the 3D skin organotypic cultures were then air-lifted and cultured with the airlifting media (culture media described above, without Y-27632 and EGF) below the insert to induce epidermal differentiation. The airlifting medium was replaced every 2 days. After 14-16 days post-airlifting, treatments were carried out.

Cantharidin or DMSO was added to culture media, and treatments were performed with technical replicates. 3D skin organotypics were then harvested and formalin-fixed for 24 hours. Fixed tissues were embedded in wax before sectioning and staining with H&E protocol for histology imaging. Cell culture medium was harvested for cytokine analysis with ELISA and western blotting.

### KINOMescan Analysis

Well-characterized kinase inhibitors with available KINOMEscan data were selected. N/TERTs expressing NLRP1-86-275_SNAP were pre-treated with kinase inhibitors before stimulation with 50 μM cantharidin. Kinase inhibitors that reduced phosphorylation of the rhodamine labelled NLRP1 DR-SNAP were classified as positive compounds while kinase inhibitors that did not affect phosphorylation were classified as negative compounds. For each kinase inhibitor, a cutoff based on kinase inhibition data was assigned to obtain lists of kinases inhibited by each inhibitor. Kinases inhibited by positive compounds were intersected with inner join logic. This set of kinases were then intersected with kinases inhibited by negative compounds with anti join logic. Analyses were done in RStudio and an annotated code is available in the supplementary.

### Recombinant protein expression and purification

All plasmids for NLRP1DR (a.a. 86-254) recombinant proteins were cloned with a C-terminal SNAP tag (NEB, N9181) into a pET47b(+) vector with C-terminal SNAP tag between NdeI and EcoRI restriction sites. Primers (IDT Technologies) were designed for site-directed mutagenesis to replace Serine and Threonine residues in the NLRP1DR STless construct.

The plasmids constructed were transformed into BL21(DE3) (NEB, C2527) and grown in lysogeny broth at 37 °C for 16-18 hours, in a shaking incubator (220 rpm). The starting culture was then used to inoculate a larger volume of lysogeny broth, which was grown to OD600 of ∼0.6-0.8. Temperature was then lowered to 16 °C, protein expression was induced with 0.5 mM IPTG, and bacteria were cultured overnight. Bacteria were harvested at 4000g (Avanti JXN series, Beckman Coulter) for 15 min and then resuspended with lysis buffer (20 mM Tris-HCl pH 8.0, 300 mM NaCl, 20 mM imidazole, and 10% glycerol). Cells were then lysed by Emulsiflex-C3 Homogenizer and centrifuged at a speed of 30,000g (Avanti JXN series, Beckman Coulter) for 30 minutes at 4°C. The supernatant was passed through a pre-equilibrated column containing Ni-NTA agarose beads (BioBasic, SA005100) by gravity. The column was then washed with a lysis buffer containing 20 mM imidazole to remove any non-specific binding proteins. His-tagged proteins were then eluted with increasing imidazole concentration (100-300 mM, each step a 50 mM increment) and verified via SDS-PAGE with Coomassie blue staining.

### *In vitro* kinase assay

Per reaction, 500 ng of recombinant NLRP1DR-SNAP was incubated with 100 ng of recombinant ZAKα (Abcam, ab89852), p38α (Abcam, ab271606), or TAK1-TAB1 (Invitrogen, PV4394) and reaction buffer (20 mM HEPES-KOH pH 7.0, 5 mM MgCl_2_, 2 mM ATP, and 0.5 M DTT) at 30 °C for 1 hour. For lambda phosphatase treatments, a final concentration of 1X NEBuffer for Protein MetalloPhosphatases (PMP), 1 mM MnCl_2_, and 0.4 μL of Lambda Protein Phosphatase (NEB, P0753S) was added to the reaction mix and incubated at 30 °C for 30 minutes. To prepare proteins for visualization, a final concentration of 0.5 μM SNAP-Cell TMR-Star (NEB, S9105) was added to the reaction mix and incubated at room temperature for 5 minutes. A 5X Laemmli buffer was then added to a final concentration of 1X, and the reaction mix was run on an 8% Phos-tag SDS-PAGE acrylamide gel. The gel was visualized with a ChemiDoc MP (Bio-Rad) using a standard rhodamine filter.

### Liquid chromatography with tandem mass spectroscopy

Phos-tag SDS-PAGE gels with *in vitro* kinase reaction samples loaded were stained with Coomassie Brilliant Blue and destained. Protein bands corresponding to phosphorylated proteins were excised and digested with chymotrypsin (Roche, 11418467001) according to the manufacturer’s protocol. Conventional peptide extraction and clean-up from digestion were then performed. The peptides were separated and analyzed using a Dionex Ultimate 3000 RSLCnano system coupled to a Q Exactive instrument (Thermo Fisher Scientific, MA, USA). Separation was performed on a Dionex EASY-Spray 75 μm × 10 cm column packed with PepMap C18 3 μm, 100 Å (Thermo Fisher Scientific) using solvent A (0.1% formic acid) and solvent B (0.1% formic acid in 100% ACN) at flow rate of 300 nL/min with a 60-min gradient. Peptides were then analyzed on a Q Exactive apparatus with an EASY nanospray source (Thermo Fisher Scientific) at an electrospray potential of 1.5 kV. Raw data files were processed and searched using Proteome Discoverer 2.1 (Thermo Fisher Scientific). The Mascot algorithm was then used for data searching to identify proteins represented. All mass spec-related experiments were undertaken at the School of Biological Sciences Mass Spec Core Facility in Nanyang Technological University, Singapore.

### FACS Analysis

Single cell suspensions were transferred to 5mL tubes and the LSRFortessa X-20 (BD Biosciences) was used for sample acquisition. Gating, analysis, and data presentation was done with Flowjo (Treestar).

### Statistical analysis

All error bars represent S.E.M. Statistical analyses were performed using Prism 10 (GraphPad). The methods for statistical analysis are included in the figure legends.

**Figure S1:**
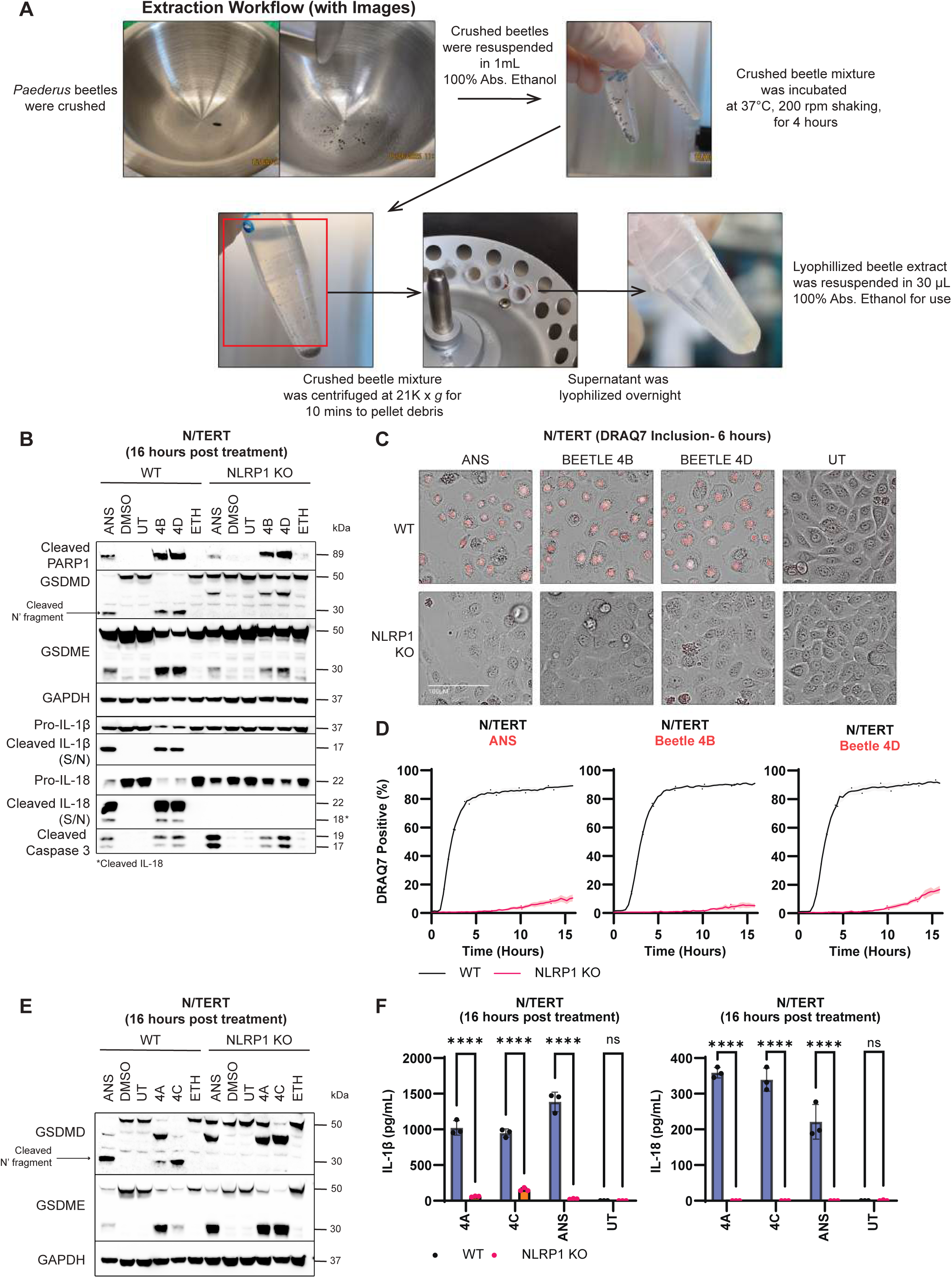
*Paederus* beetle extract induces NLRP1-dependent pyroptosis in epithelial cells. **A.** Workflow of how *Paederus* beetle extracts were obtained. **B.** Immunoblot of cleaved PARP1, GSDMD (full length and cleaved), GSDME (full length and cleaved), GAPDH, Pro-IL-1β, cleaved IL-1β, IL-18 (full length and cleaved), cleaved caspase-3 from WT and NLRP1 KO N/TERTs following treatment with 1 μM anisomycin, DMSO (vehicle control), X1500 dilution of beetle 4B and beetle 4D extracts, and ethanol (vehicle control for beetle extracts). Cells, floaters, and cell culture media were harvested after 16 hours of treatment. **C.** Representative images of DRAQ7-positive and **D.** quantification of the percentage of DRAQ7-positive WT and NLRP1 KO N/TERTs following treatment with 1 μM anisomycin, X1500 dilution of beetle 4B and beetle 4D extracts. Images were taken after 6 hours of treatment. **E.** Immunoblot of GSDMD (full length and cleaved), GSDME (full length and cleaved), and GAPDH from WT and NLRP1 KO N/TERTs following treatment with 1 μM anisomycin, DMSO (vehicle control), X1500 dilution of beetle 4A and beetle 4C extracts, ethanol (vehicle control for beetle extracts). Cells, floaters, and cell culture media were harvested after 16 hours of treatment. **F.** IL-1β and IL-18 ELISA of WT and NLRP1 KO N/TERTs following treatment with 1 μM anisomycin and X1500 dilution of beetle 4A and beetle 4C extracts. Cell culture media was harvested after 16 hours of treatment. Error bars represent SEM from three technical replicates, where one replicate refers to an independent sample. Significance values were calculated based on two-way ANOVA followed by Sidak’s test for multiple pairwise comparisons (F). ns, nonsignificant; *P < 0.05; ****P < 0.0001.

**Figure S2.**
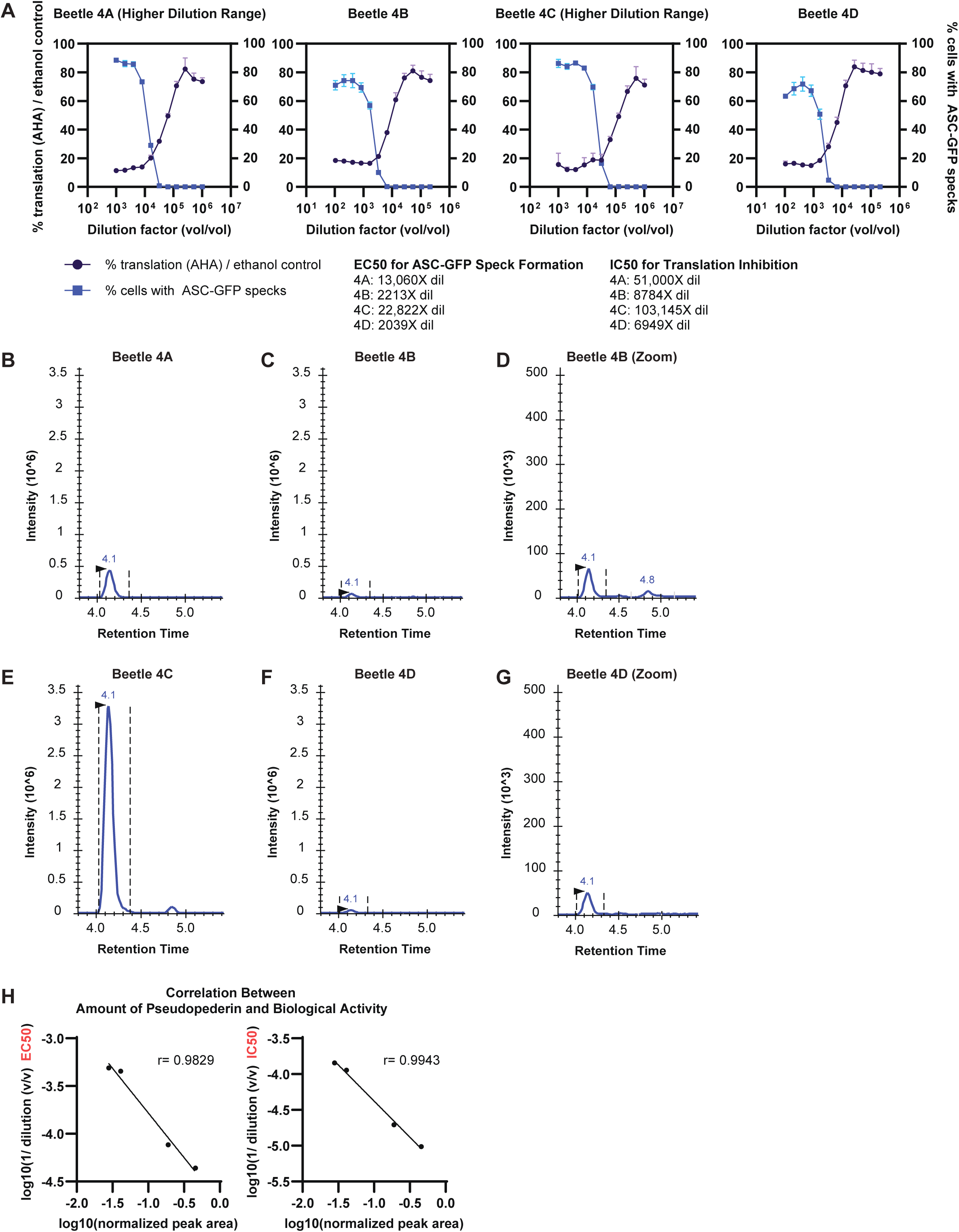
Biological activity of beetle extracts is correlated with the amount of pederin detected by mass spectrometry. **A.** Percentage of ASC-GFP speck forming A549 NLRP1 ASC-GFP reporter cells and percentage of AHA incorporation/ vehicle control, as a proxy for translation inhibition, plotted against dilution factor of respective beetle extracts. EC50 was calculated from the percentage of ASC-GFP speck forming cells. IC50 was calculated from the percentage of AHA incorporation/ vehicle control. **B-G.** Liquid chromatogram retention times of pederin detected in respective beetle extracts. **H.** Correlation between the amount of pseudopederin detected by mass spectrometry and EC50 or IC50 of beetles 4A-4D, as proxies for biological activity. Simple linear regression was performed and Pearson correlation (r value) was calculated in GraphPad Prism.

**Figure S3.**
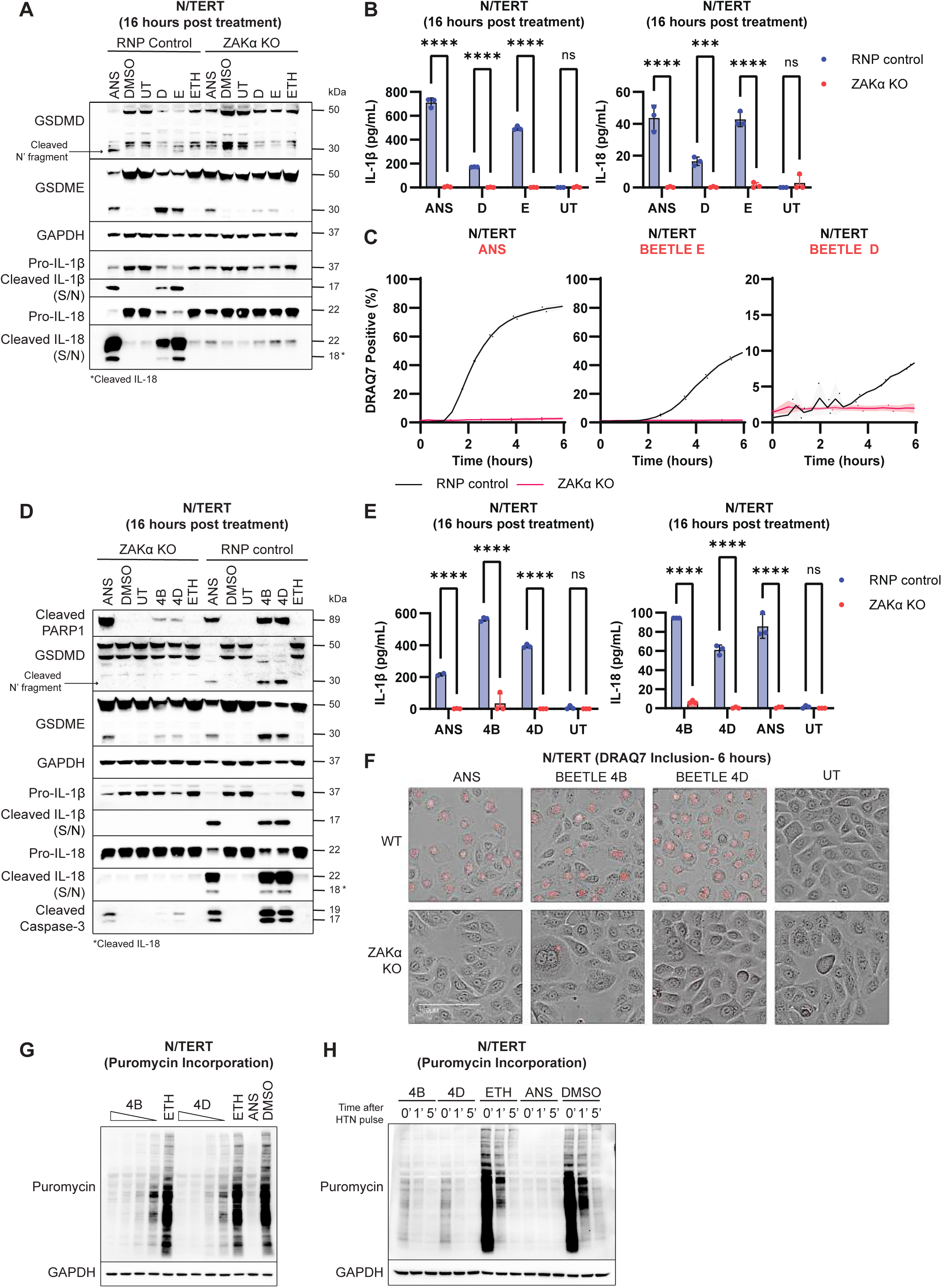
*Paederus* beetle extract induces ZAKα-dependent pyroptosis in epithelial cells. **A.** Immunoblot of GSDMD (full length and cleaved), GSDME (full length and cleaved), GAPDH, Pro-IL-1β, cleaved IL-1β, and IL-18 (full length and cleaved) from RNP control and ZAKα KO N/TERTs following treatment with 1 μM anisomycin, DMSO (vehicle control), 52X dilution of beetle D extract, 104X dilution of beetle E extract, and ethanol (vehicle control for beetle extracts). Cells, floaters, and cell culture media were harvested after 16 hours of treatment. **B.** IL-1β and IL-18 ELISA, and **C.** quantification of the percentage of DRAQ7-positive RNP control and ZAKα KO N/TERTs following treatment with 1 μM anisomycin, 52X dilution of beetle D extract and 104X dilution of beetle E extract. Cell culture media was harvested after 16 hours of treatment. **D.** Immunoblot of cleaved PARP1, GSDMD (full length and cleaved), GSDME (full length and cleaved), GAPDH, Pro-IL-1β, cleaved IL-1β, IL-18 (full length and cleaved), and cleaved caspase-3 from RNP control and ZAKα KO N/TERTs following treatment with 1 μM anisomycin, DMSO (vehicle control), X1500 dilution of beetle 4B and 4D extracts, and ethanol (vehicle control for beetle extracts). Cells, floaters, and cell culture media were harvested after 16 hours of treatment.**E.** IL-1β and IL-18 ELISA and **F.** representative images of DRAQ7-positive RNP control and ZAKα KO N/TERTs following treatment with 1 μM anisomycin, and X1500 dilution of beetle 4B and 4D extracts. Cells were imaged after 6 hours of treatment and cell culture media was harvested after 16 hours of treatment. **G.** Immunoblot of puromycin from N/TERTs following treatment with serial dilution of beetle 4B and 4D extracts, ethanol (vehicle control for beetle extracts), 1 μM anisomycin, and DMSO (vehicle control). Cells were pulsed with 10 μg/mL puromycin after 3 hours of treatment to label translating peptides. **H.** Immunoblot of puromycin from N/TERTs following treatment with X1500 dilution of beetle 4B and 4D extracts, ethanol (vehicle control for beetle extracts), 1 μM anisomycin, and DMSO (vehicle control). Translation initiation was inhibited by 2 μg/mL harringtonine for 5 or 10 minutes for actively translating ribosomes to run off. Cells were pulsed with 10 μg/mL puromycin after 3 hours of treatment to label remaining translating peptides.

**Figure S4:**
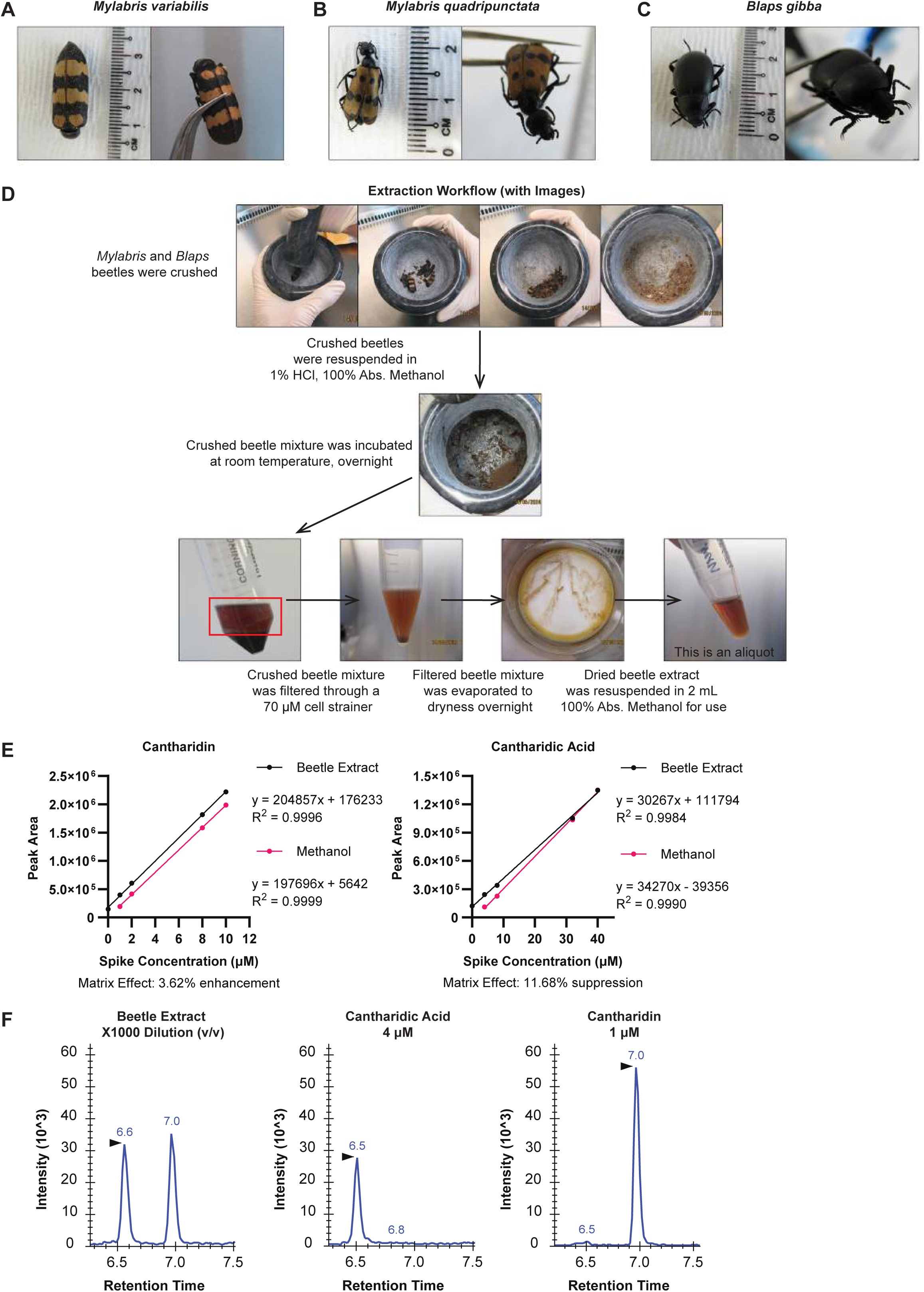
Cantharidin and cantharidic acid in *Mylabris variabilis* beetle extracts were detected by mass spectrometry. **A-D.** Workflow of how *Mylabris variabilis or quadripunctata* and *Blaps gibba* beetle extracts were obtained. **E.** Standard addition curves constructed to determine the concentrations of cantharidin and cantharidic acid in beetle extracts and matrix effects. **F.** Liquid chromatogram retention times of cantharidin and cantharidic acid detected in *Mylabris variabilis* beetle extracts, compared against commercial cantharidic acid, and cantharidin standards.

**Figure S5:**
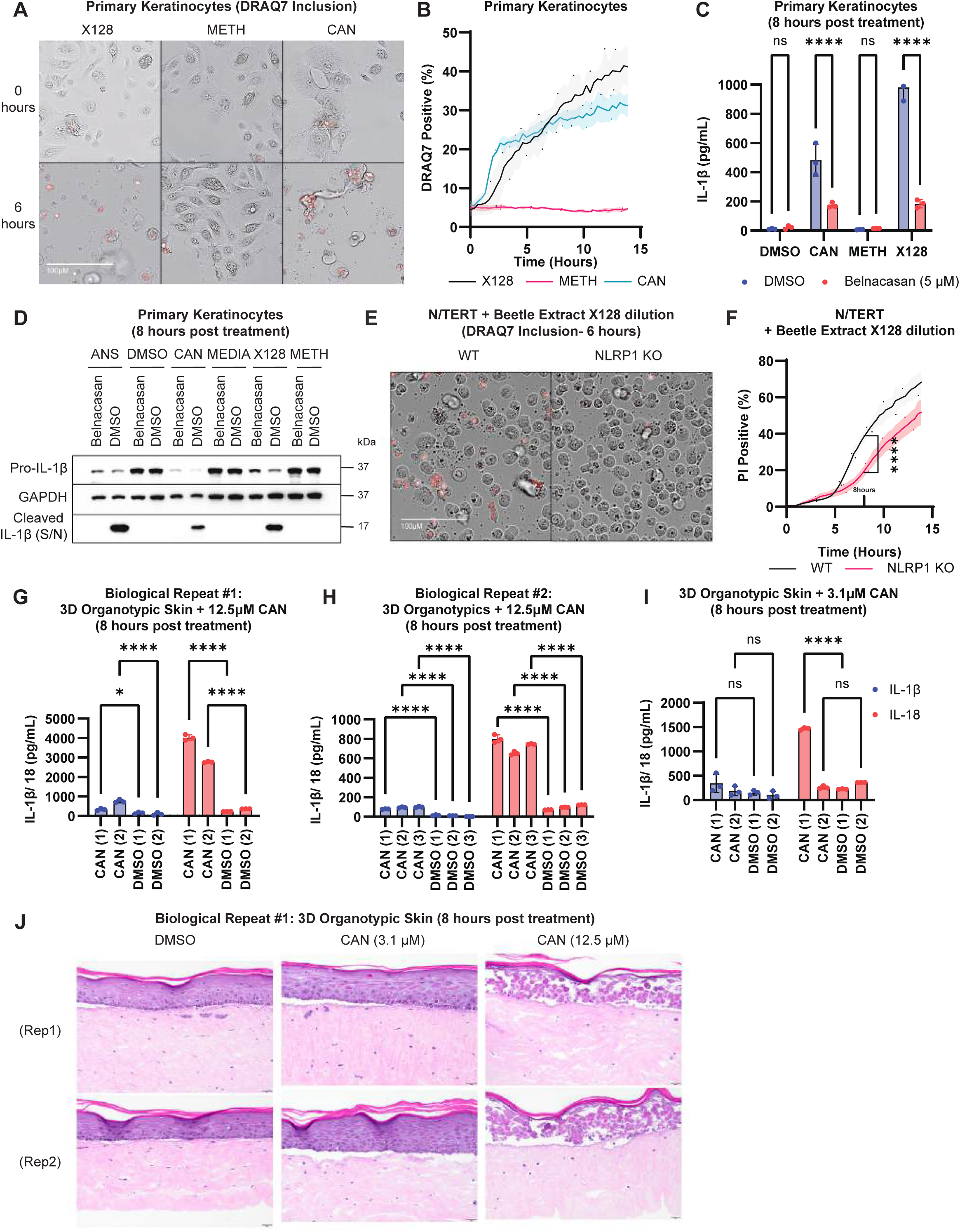
*Mylabris* beetle extract and cantharidin induce a mixed mode of cell death involving NLRP1. **A.** Representative images of DRAQ7-positive and **B.** quantification of the percentage of DRAQ7-positive primary keratinocytes following treatment with X128 dilution of *Mylabris variabilis* beetle extract, methanol (vehicle control for beetle extracts), and 50 μM cantharidin. Cells were imaged after 0 and 6 hours of treatment. **C.** IL-1β ELISA and **D.** immunoblot of Pro-IL-1β, GAPDH, and cleaved IL-1β from primary keratinocytes pre-treated with 5 μM belnacasan (caspase-1 inhibitor) before stimulation with DMSO (vehicle control), 50 μM cantharidin, methanol (vehicle control for beetle extracts), and X128 dilution of *Mylabris variabilis* beetle extract. Cells and cell culture media were harvested after 8 hours of treatment. **E.** Representative images of DRAQ7-positive and **F.** quantification of the percentage of DRAQ7-positive WT vs NLRP1 KO N/TERTs following treatment with X128 dilution of *Mylabris* beetle extract. Cells were imaged after 6 hours of treatment. **G-I.** IL-1β and IL-18 ELISA and **J.** representative images from 3D organotypic skin 3D organotypic skin sections stained with H&E following treatment with 12.5 μM or 3.1 μM cantharidin and DMSO (vehicle control). Cell culture media was harvested and 3D organotypic skin was fixed after 8 hours of treatment. Error bars represent SEM from three technical replicates, where one replicate refers to an independent sample. Significance values were calculated based on Wilcoxon matched-pairs signed rank test (F). ns, nonsignificant; *P < 0.05; ****P < 0.0001.

**Figure S6:**
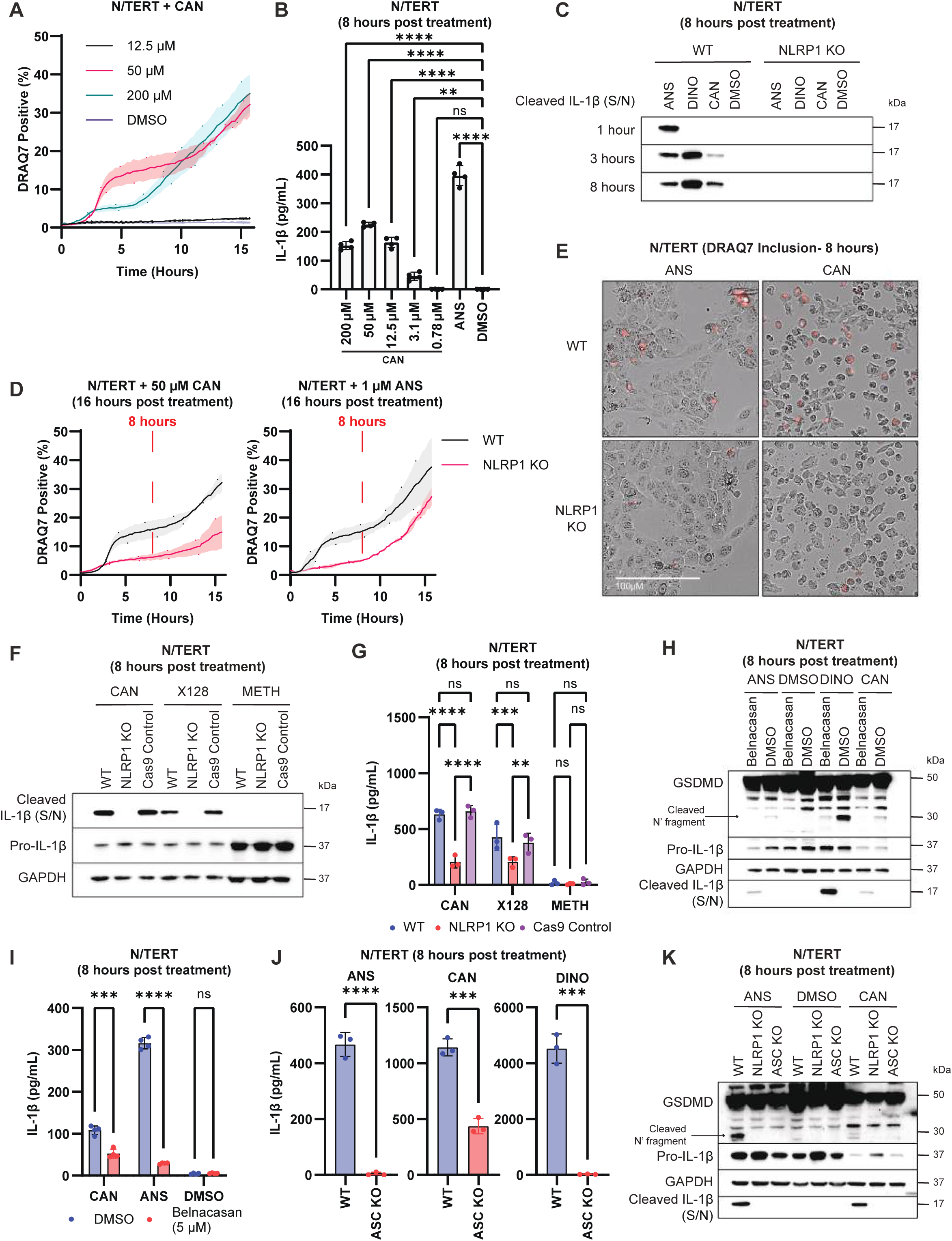
Cantharidin induces a mixed mode of cell death involving NLRP1 in immortalized keratinocytes. **A.** Quantification of the percentage of DRAQ7-positive N/TERTs following treatment with a titration of cantharidin and DMSO (vehicle control). **B.** IL-1β ELISA from N/TERTs following treatment with a titration of cantharidin, 1 μM anisomycin, and DMSO (vehicle control). Cell culture media was harvested after 8 hours of treatment. **C.** Cleaved IL-1β from WT and NLRP1 KO N/TERTs following treatment with 1 μM anisomycin, 6 μM VbP, 50 μM cantharidin, and DMSO (vehicle control). Cell culture media was harvested at various timepoints. **D.** Quantification of the percentage and **E.** representative images of DRAQ7-positive WT and NLRP1 KO N/TERTs following treatment with 50 μM cantharidin or 1 μM anisomycin. Cells were imaged 8 hours after treatment. **F.** Immunoblot of cleaved IL-1β, Pro-IL-1β, and GAPDH and **G.** IL-1β ELISA from WT, NLRP1 KO, and Cas9 control N/TERTs following treatment with 50μM cantharidin, X128 dilution of *Mylabris variabilis* beetle extract, and methanol (vehicle control for beetle extracts). Cells and cell culture media were harvested after 8 hours of treatment. **H.** Immunoblot of GSDMD (full length and cleaved), Pro-IL-1β, cleaved IL-1β, and GAPDH and **I.** IL-1β ELISA from N/TERTs following pre-treatment with 5 μM belnacasan (caspase-1 inhibitor) before stimulation with 1 μM anisomycin, DMSO (vehicle control), 12.5 nM dinophysis, and 50 μM cantharidin. Cells, floaters, and cell culture media were harvested after 8 hours of treatment. **J.** IL-1β ELISA and **K.** immunoblot of GSDMD (full length and cleaved), Pro-IL-1β, cleaved IL-1β, and GAPDH from WT and ASC KO N/TERTs following treatment with 1 μM anisomycin, 50 μM cantharidin, 12.5 nM dinophysis, and DMSO (vehicle control). Cells, floaters, and cell culture media were harvested after 8 hours of treatment. Error bars represent SEM from three technical replicates, where one replicate refers to an independent sample. Significance values were calculated based on one-way ANOVA followed by Dunnett’s test (B) or two-way ANOVA followed by Sidak’s test for multiple pairwise comparisons (G, I, and J). ns, nonsignificant; *P < 0.05; ****P < 0.0001.

**Figure S7:**
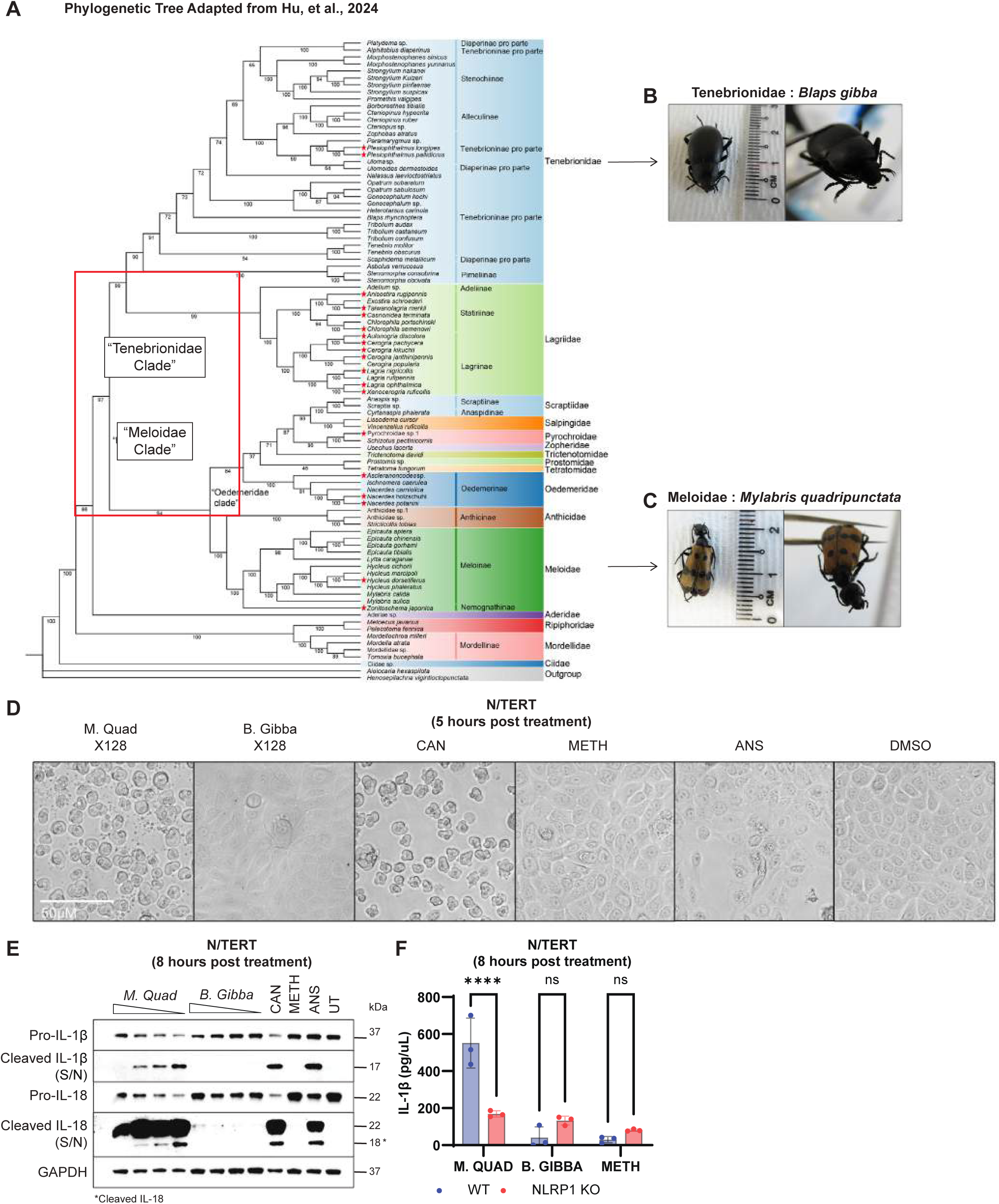
*Mylabris quadripunctata* beetle extract induces pyroptosis in immortalized keratinocytes. **A.** Phylogenetic tree adapted from <u>(Hu et al. 2024)</u>. **C.** *Mylabris quadripunctata* belongs to the *meloidae* clade of blistering beetles that contain cantharidin while **B.** *Blaps gibba* belongs to the *tenebrionidae* clade of closely related, non-blistering beetles. **D.** Representative brightfield images of N/TERTs following treatment with X128 dilution of *Mylabris quadripunctata* and *Blaps gibba* beetle extract, 50 μM cantharidin, methanol (vehicle control for beetle extracts), 1 μM anisomycin, and DMSO (vehicle control). Cells were imaged after 5 hours of treatment. **E.** Immunoblot of Pro-IL-1β, cleaved IL-1β, IL-18 (full length and cleaved), and GAPDH from N/TERTs following treatment with a titration of *Mylabris quadripunctata* and *Blaps gibba* beetle extract, 50 μM cantharidin, methanol (vehicle control for beetle extracts), and 1 μM anisomycin. Cells and cell culture media were harvested after 8 hours of treatment. **F.** IL-1β ELISA from WT and NLRP1 KO N/TERTs following treatment with X128 dilution of *Mylabris quadripunctata* and *Blaps gibba* beetle extract, and methanol (vehicle control for beetle extracts). Cell culture was harvested after 8 hours of treatment. Error bars represent SEM from three technical replicates, where one replicate refers to an independent sample. Significance values were calculated based on two-way ANOVA followed by Sidak’s test for multiple pairwise comparisons (F). ns, nonsignificant; *P < 0.05; ****P < 0.0001.

**Figure S8:**
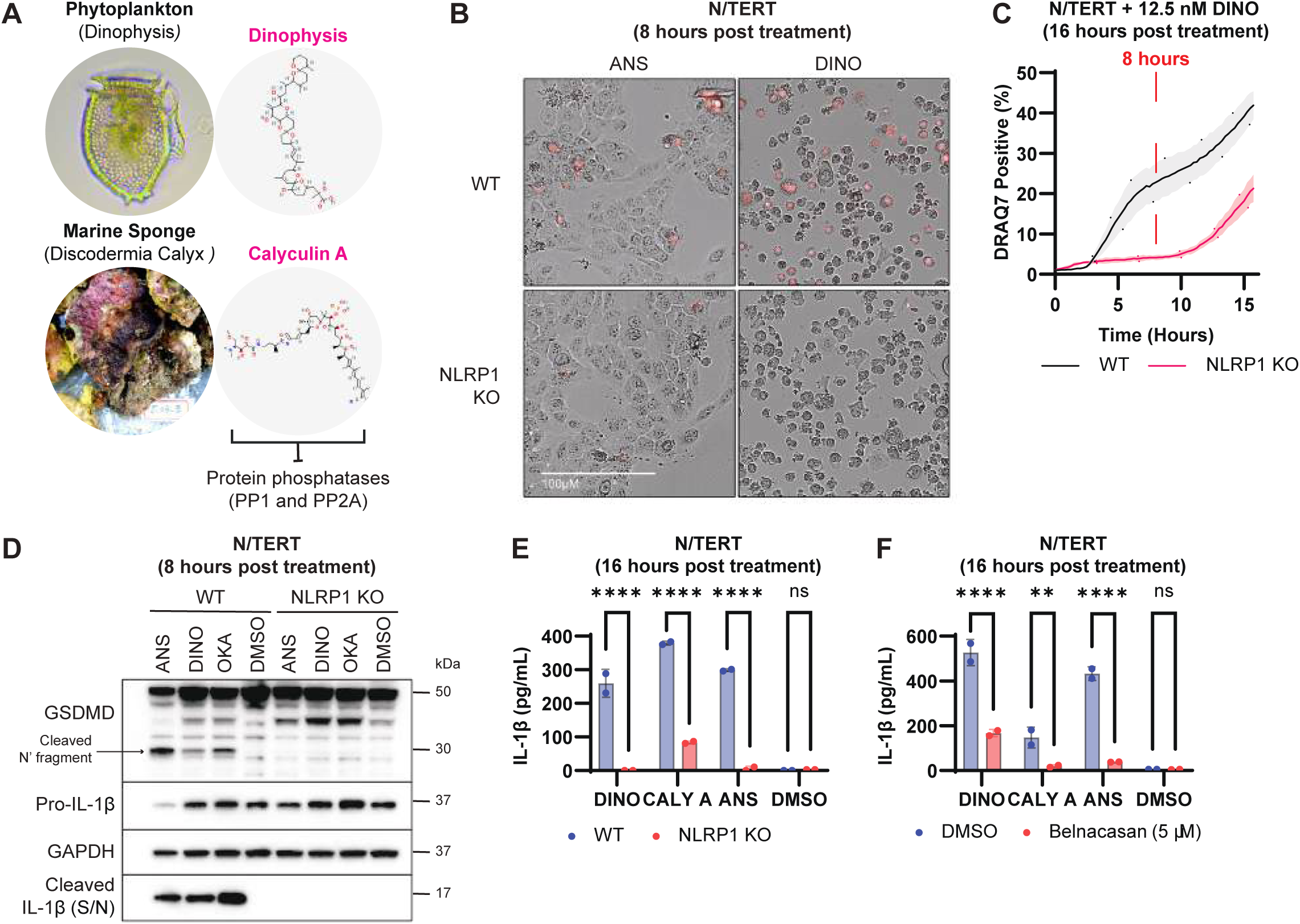
Marine toxins induce pyroptosis in immortalized keratinocytes. **A.** Dinophysis and calyculin A are naturally occurring PP1 and PP2A protein phosphatases. **B.** Representative images and **C.** quantification of the percentage of DRAQ7-positive WT and NLRP1 KO N/TERTs following treatment with 12.5 nM dinophysis or 1 μM anisomycin. Cells were imaged after 8 hours of treatment. **D.** Immunoblot of GSDMD (full length and cleaved), Pro-IL-1β, cleaved IL-1β, and GAPDH from WT and NLRP1 KO N/TERTs following treatment with 1 μM anisomycin, 12.5 nM dinophysis, 62.5 nM okadaic acid, and DMSO (vehicle control). Cells, floaters, and cell culture media were harvested after 8 hours of treatment. **E.** IL-1β ELISA from WT and NLRP1 KO N/TERTs and **F.** N/TERTs pre-treated with DMSO or 5 μM belnacasan (caspase-1 inhibitor) before stimulation with 12.5 nM dinophysis, 2 nM calyculin A, 1 μM anisomycin, and DMSO (vehicle control). Cell culture media was harvested after 16 hours of treatment. Error bars represent SEM from three technical replicates, where one replicate refers to an independent sample. Significance values were calculated based on two-way ANOVA followed by Sidak’s test for multiple pairwise comparisons (E and F). ns, nonsignificant; *P < 0.05; ****P < 0.0001.

**Figure S9:**
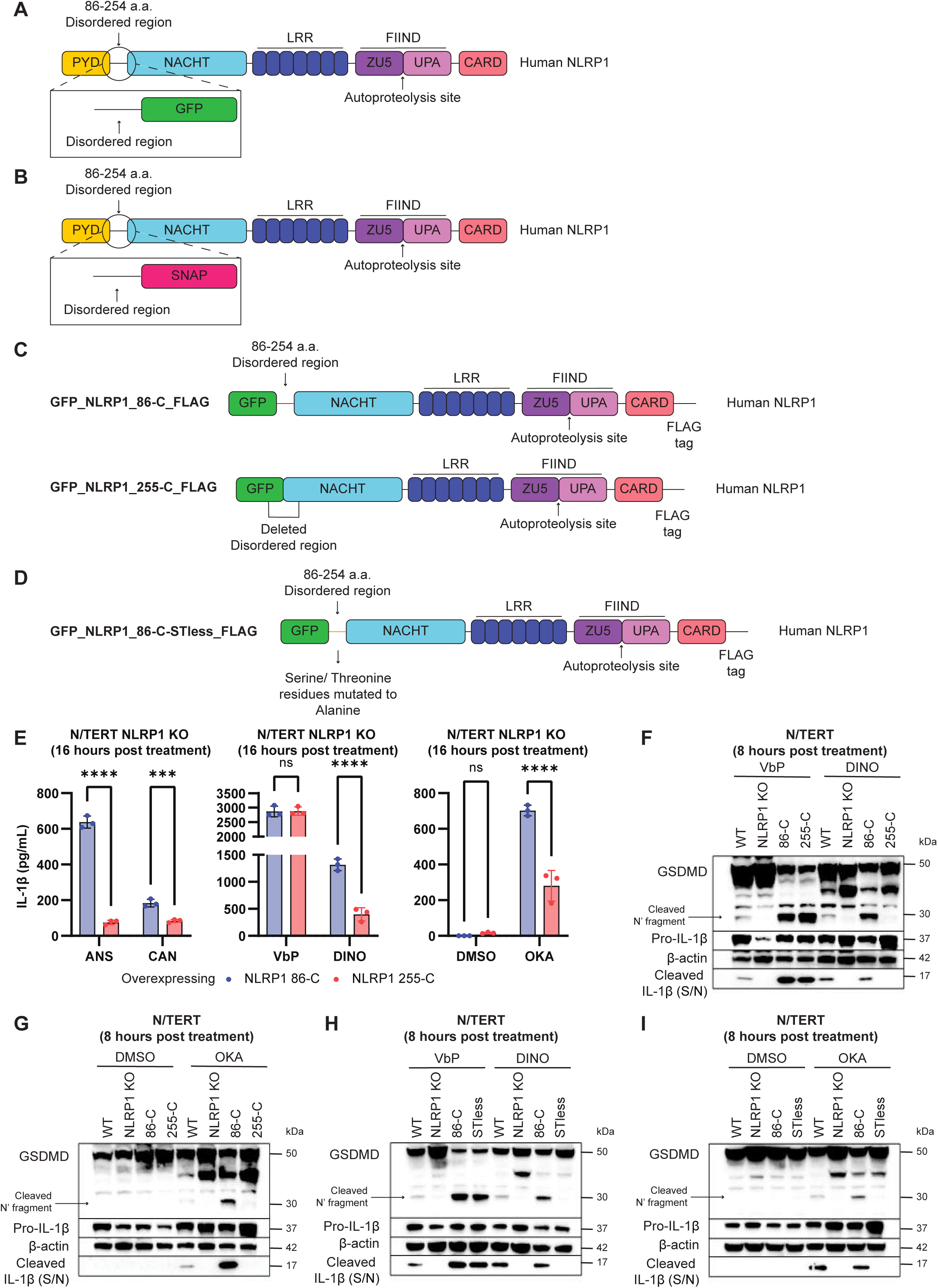
Phosphatase inhibitors require S/T residues in the disordered region for activation of the NLRP1 inflammasome. **A.** Domain architecture of human NLRP1. Disordered region between the PYD and NACHT domains (residues 86-254a.a.) was cloned into a C’ terminal GFP- or **B.** SNAP-tagged vector and overexpressed in N/TERTs. **C.** Disordered region was truncated in the GFP_NLRP1-255-C_FLAG construct or **D.** serine and threonine residues within the NLRP1-DR were mutated to alanine residues in the GFP_NLRP1-86-C_FLAG (STless) construct and overexpressed in N/TERTs. **E.** IL-1β ELISA from NLRP1 KO N/TERTs overexpressing GFP_NLRP1-86-C_FLAG (WT) and truncated GFP_NLRP1-255-C_FLAG following treatment with 1 μM anisomycin, 50 μM cantharidin, 6 μM VbP, 12.5 nM dinophysis, DMSO (vehicle control), and 62.5 nM okadaic acid. Cell culture media was harvested after 8 hours of treatment. **F-G.** Immunoblot of GSDMD (full length and cleaved), Pro-IL-1β, cleaved IL-1β, and GAPDH from NLRP1 KO N/TERTs overexpressing GFP_NLRP1-86-C_FLAG (WT) and truncated GFP_NLRP1-255-C_FLAG or **H-I.** GFP_NLRP1-86-C_FLAG (STless) following treatment with 6 μM VbP, 12.5 nM dinophysis, DMSO (vehicle control), and 62.5 nM okadaic acid. Cells, floaters and cell culture media were harvested after 8 hours of treatment. Error bars represent SEM from three technical replicates, where one replicate refers to an independent sample. Significance values were calculated based on two-way ANOVA followed by Sidak’s test for multiple pairwise comparisons (E). ns, nonsignificant; *P < 0.05; ****P < 0.0001.

**Figure S10:**
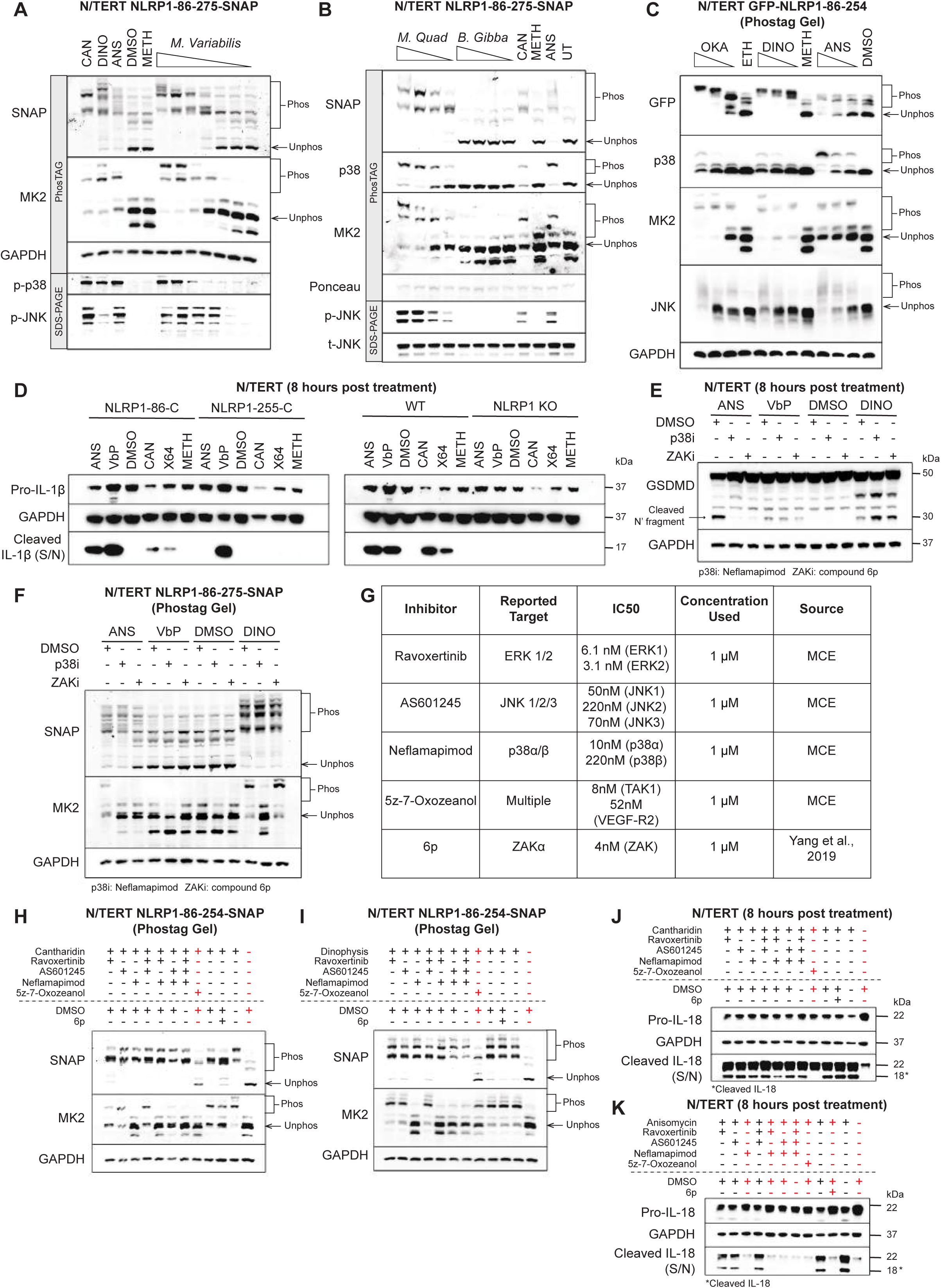
Phosphatase-dependent phosphorylation of the NLRP1 DR and activation of the NLRP1 inflammasome is independent of ERK and JNK. **A.** Rhodamine labelled SNAP-tagged NLRP1-DR or immunoblot of MK2, and GAPDH following PhosTag-SDS-Agarose-PAGE or p-p38 and p-JNK following SDS-PAGE from N/TERTs pre-treated with 5μM emricasan (pan-caspase inhibitor) before stimulation with 50 μM cantharidin, 12.5 nM dinophysis, 1 μM anisomycin, DMSO (vehicle control), methanol (vehicle control for beetle extract), and a dilution titration of *Mylabris variabilis* beetle extract. Cells were harvested after 3 hours of treatment. Phosphorylated and unphosphorylated species of each protein are indicated. **B.** Rhodamine labelled SNAP-tagged NLRP1-DR or immunoblot of p38, MK2, and GAPDH following PhosTag-SDS-Agarose-PAGE or t-JNK and p-JNK following SDS-PAGE from N/TERTs pre-treated with 5 μM emricasan (pan-caspase inhibitor) before stimulation with a dilution titration of *Mylabris quadripunctata* and *Blaps gibba* beetle extracts, 50 μM cantharidin, methanol (vehicle control for beetle extracts), and 1 μM anisomycin. Cells were harvested after 3 hours of treatment. Phosphorylated and unphosphorylated species of each protein are indicated. **C.** Immunoblot of GFP-tagged NLRP1-DR, p38, MK2, JNK, and GAPDH following PhosTag-SDS-Agarose-PAGE from N/TERTs pre-treated with 5 μM emricasan (pan-caspase inhibitor) before stimulation with a titration of okadaic acid, dinophysis, anisomycin, and ethanol/ methanol/ DMSO (vehicle controls). Cells were harvested after 3 hours of treatment. Phosphorylated and unphosphorylated species of each protein are indicated. **D.** Immunoblot of Pro-IL-1β, cleaved IL-1β, and GAPDH from WT, NLRP1 KO, and NLRP1 KO N/TERTs overexpressing GFP_NLRP1-86-C_FLAG (WT) or truncated GFP_NLRP1-255-C_FLAG following treatment with 1 μM anisomycin, 6 μM VbP, DMSO (vehicle control), 50 μM cantharidin, X64 dilution of *Mylabris variabilis* beetle extract, and methanol (vehicle control for beetle extracts). Cells and cell culture media were harvested after 8 hours of treatment. **E.** Immunoblot of GSDMD (full length and cleaved) and GAPDH from N/TERTs pre-treated with 5μM emricasan (pan-caspase inhibitor) and DMSO (vehicle control), 1 μM neflamapimod (p38 inhibitor), or 1 μM 6p (ZAK inhibitor) before stimulation with 1 μM anisomycin, 6 μM VbP, DMSO (vehicle control), and 12.5 nM dinophysis. Cells, floaters, and cell culture media were harvested after 8 hours of treatment. **F.** Rhodamine labelled SNAP-tagged NLRP1-DR or immunoblot of MK2 and GAPDH following PhosTag-SDS-Agarose-PAGE from N/TERTs pre-treated with 5 μM emricasan (pan-caspase inhibitor) and DMSO (vehicle control), 1 μM of Neflamapimod (p38 inhibitor), or 6p (ZAK inhibitor) before stimulation with 1 μM anisomycin, 3 μM VbP, DMSO (vehicle control), and 12.5 nM dinophysis. Cell lysates were harvested after 3 hours of treatment. Phosphorylated and unphosphorylated species of each protein are indicated. **G.** Summary of kinase inhibitors used in the following panels. **H.** Rhodamine labelled SNAP-tagged NLRP1-DR or immunoblot of MK2 and GAPDH following PhosTag-SDS-Agarose-PAGE from N/TERTs pre-treated with 5 μM emricasan (pan-caspase inhibitor) and DMSO (vehicle control), or 1 μM of various kinase inhibitors before stimulation with 50 μM cantharidin or **I.** 12.5 nM dinophysis. Cells were harvested after 3 hours of treatment. Phosphorylated and unphosphorylated species of each protein are indicated. **J.** Immunoblot of IL-18 (full length and cleaved) and GAPDH from N/TERTs pre-treated with DMSO (vehicle control) or 1 μM of various kinase inhibitors before stimulation with 50 μM cantharidin or **K.** 1 μM anisomysin. Cells and cell culture media were harvested after 8 hours of treatment.

**Figure S11:**
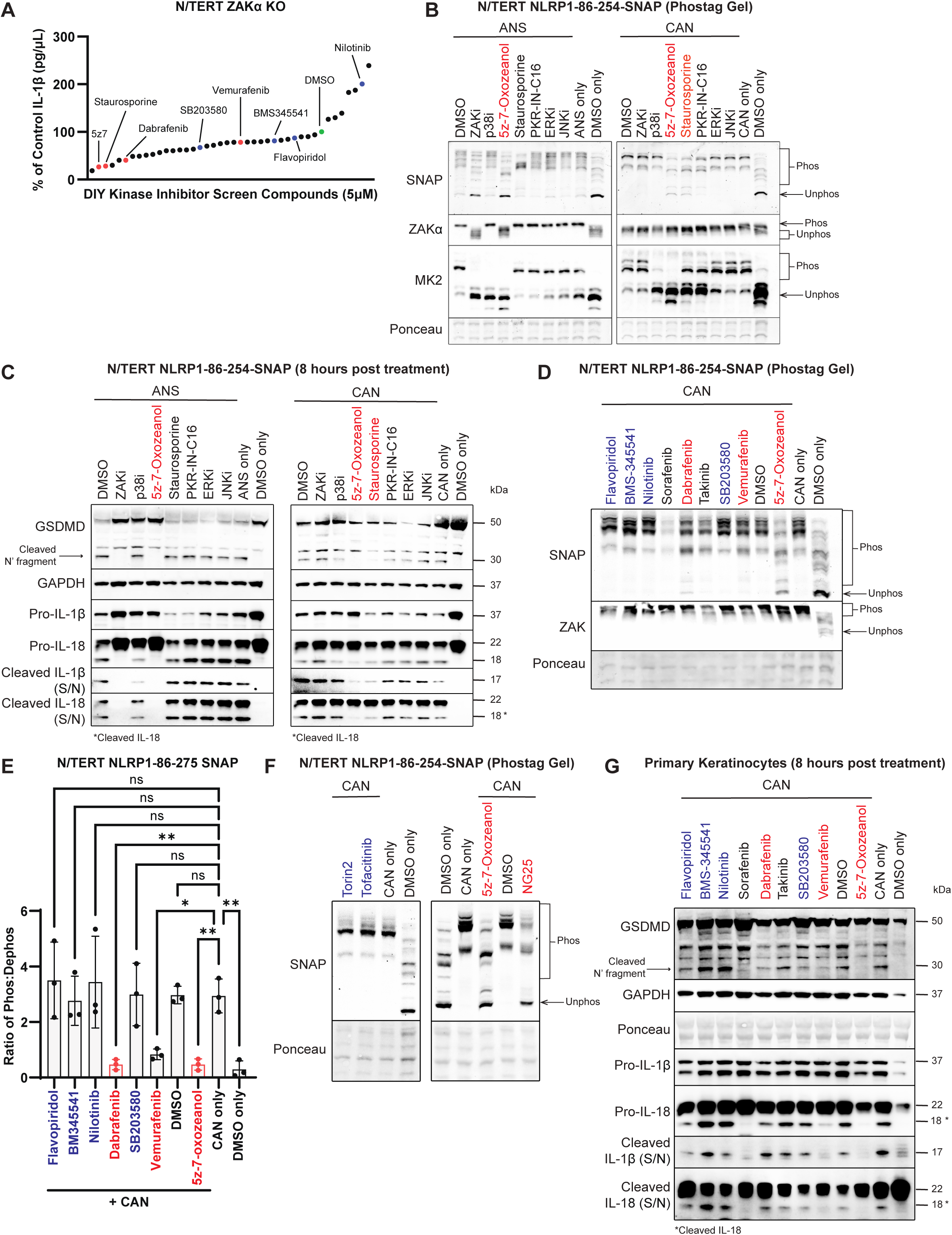
Phosphorylation and functional analyses done in the chemical screen. **A.** IL-1β release as a percentage of control (DMSO) from ZAKα KO N/TERTs pre-treated with 5 μM of various kinase inhibitors before stimulation with 12.5 nM dinophysis. ZAKα KO N/TERTs and dinophysis were chosen as this combination gave the strongest induction of NLRP1 activation, thus ensuring a healthy dynamic range to detect the effect of kinase inhibitors on phosphatase inhibitor-dependent NLRP1 activation. Cell culture media was harvested after 8 hours of treatment. Positive kinase inhibitors are indicated in red, DMSO control indicated in green, and negative kinase inhibitors are indicated in blue. **B.** Rhodamine labelled SNAP-tagged NLRP1-DR or immunoblot of ZAK, MK2, and GAPDH following PhosTag-SDS-Agarose-PAGE from N/TERTs pre-treated with 5 μM emricasan (pan-caspase inhibitor) and 1 μM of indicated kinase inhibitors before stimulation with 1 μM anisomycin (left) or 50 μM cantharidin (right). Cells were harvested after 3 hours of treatment. Phosphorylated and unphosphorylated species of each protein are indicated. Kinase inhibitors able to reduce anisomycin or cantharidin-dependent phosphorylation of NLRP1-DR are highlighted in red. **C.** Immunoblot of GSDMD (full length and cleaved), Pro-IL-1β, cleaved IL-1β, IL-18 (full length and cleaved), and GAPDH of N/TERTs pre-treated with 1 μM of indicated kinase inhibitors before stimulation for with 1 μM anisomycin (left) or 50 μM cantharidin (right). Cells, floaters, and cell culture media were harvested after 8 hours of treatment. Kinase inhibitors able to reduce anisomycin or cantharidin-dependent activation of NLRP1 are highlighted in red. **D.** Rhodamine labelled SNAP-tagged NLRP1-DR and immunoblot of ZAK following PhosTag-SDS-Agarose-PAGE from N/TERTs pre-treated with 5 μM emricasan (pan-caspase inhibitor) and 1 μM of indicated kinase inhibitors before stimulation with 50 μM cantharidin. Cells were harvested after 3 hours of treatment. Phosphorylated and unphosphorylated species of each protein are indicated. Positive kinase inhibitors are indicated in red and negative kinase inhibitors are indicated in blue. **E.** Quantification of the intensity of phosphorylated vs unphosphorylated bands of NLRP1-DR in Figure S11D. Quantification was done with the ImageLab software. **F.** Rhodamine labelled SNAP-tagged NLRP1-DR following PhosTag-SDS-Agarose-PAGE from N/TERTs pre-treated with 5 μM emricasan (pan-caspase inhibitor) and 1 μM of indicated kinase inhibitors before stimulation with 50 μM cantharidin. Cells were harvested after 3 hours of treatment. Phosphorylated and unphosphorylated species of each protein are indicated. Positive kinase inhibitors are indicated in red and negative kinase inhibitors are indicated in blue. **G.** Immunoblot of GSDMD (full length and cleaved), Pro-IL-1β, cleaved IL-1β, IL-18 (full length and cleaved), and ponceau/ GAPDH of primary keratinocytes pre-treated with 1 μM of indicated kinase inhibitors before stimulation with 50 μM cantharidin. Cells, floaters, and cell culture media were harvested after 8 hours of treatment. Positive kinase inhibitors are indicated in red and negative kinase inhibitors are indicated in blue. Error bars represent SEM from three technical replicates, where one replicate refers to an independent sample. Significance values were calculated based on one-way ANOVA followed by Dunnett’s test (E). ns, nonsignificant; *P < 0.05; ****P < 0.0001.

**Figure S12:**
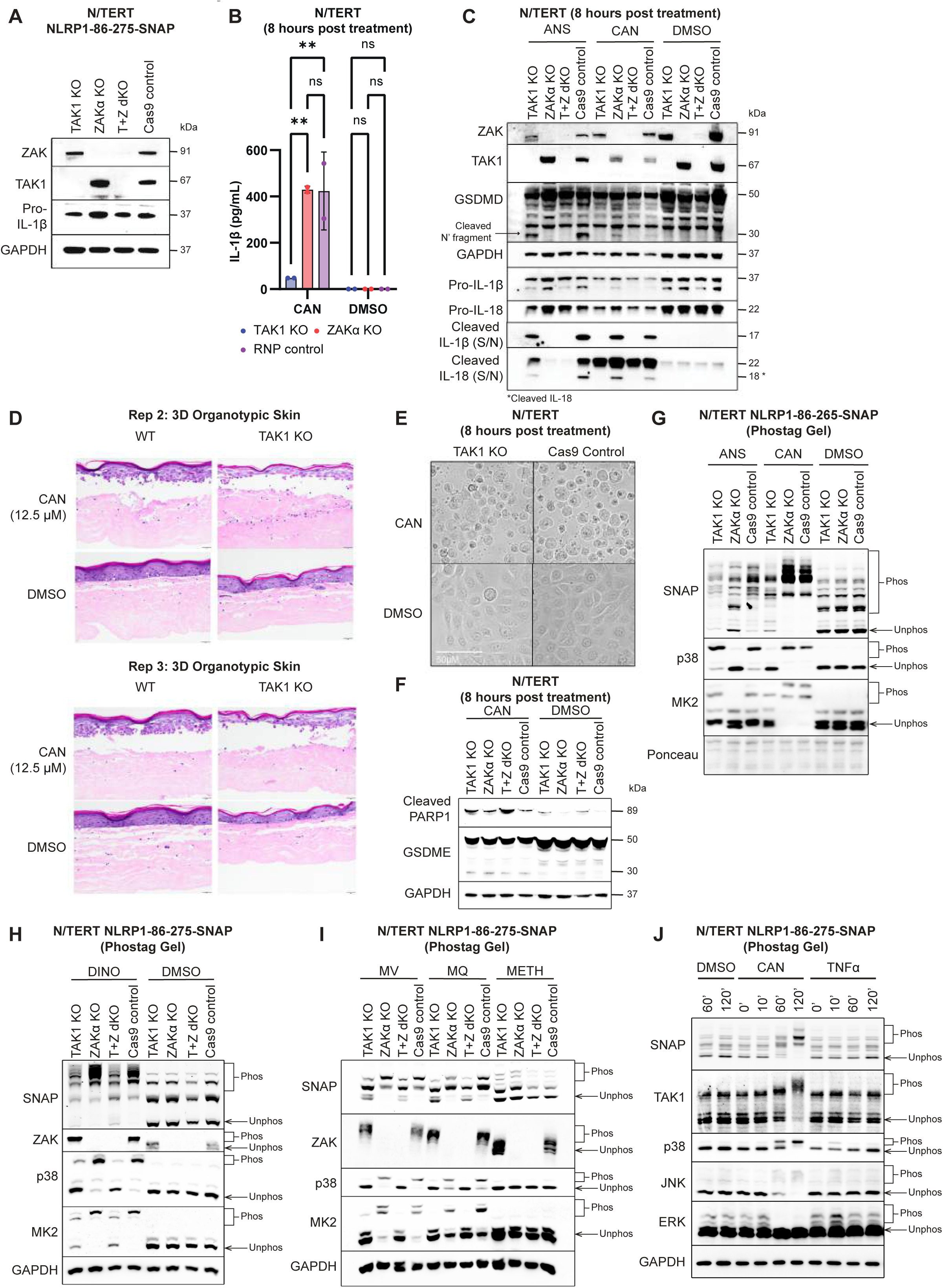
TAK1 is required for cantharidin-dependent activation of NLRP1 in immortalized keratinocytes. **A.** Immunoblot of ZAK, TAK1, Pro-IL-1β, and GAPDH from N/TERTs to verify knockout efficiency. **B.** IL-1β ELISA from TAK1 KO, ZAKα KO, and RNP control N/TERTs following treatment with 50μM cantharidin and DMSO (vehicle control). Cell culture media was harvested after 8 hours of treatment. **C.** Immunoblot of ZAK, TAK1, GSDMD (full length and cleaved), GAPDH, Pro-IL-1β, cleaved IL-1β, and IL-18 (full length and cleaved) from TAK1 KO, ZAKα KO, T+Z dKO, and RNP control N/TERTs following treatment with 1 μM anisomycin, 50 μM cantharidin, and DMSO (vehicle control). Cells, floaters, and cell culture media were harvested after 8 hours of treatment. **D.** Representative H&E images of WT and TAK1 KO 3D organotypic skin following treatment with 12.5 μM cantharidin and DMSO (vehicle control), from technical repeats. 3D organotypics were fixed after 8 hours of treatment. **E.** Representative brightfield images and **F.** immunoblot of GSDME (full length and cleaved) and GAPDH from TAK1 KO and RNP control N/TERTs following treatment with 50 μM cantharidin and DMSO (vehicle control). Cells were imaged and cells/ floaters were harvested after 8 hours of treatment. **G.** Rhodamine labelled SNAP-tagged NLRP1-DR or immunoblot of p38 and MK2 following PhosTag-SDS-Agarose-PAGE from TAK1 KO, ZAKα KO, and RNP control cells pre-treated with 5 μM emricasan (pan-caspase inhibitor) and stimulation with 1 μM anisomycin, 50 μM cantharidin, and DMSO (vehicle control). Cells were harvested after 3 hours of treatment. Phosphorylated and unphosphorylated species of each protein are indicated. **H.** Rhodamine labelled SNAP-tagged NLRP1-DR or immunoblot of ZAK, p38, and MK2 following PhosTag-SDS-Agarose-PAGE from TAK1 KO, ZAKα KO, T+Z dKO, and RNP control cells pre-treated with 5 μM emricasan (pan-caspase inhibitor) before stimulation with 12.5 nM dinophysis and DMSO (vehicle control) or **I.** X128 dilution of *mylabris variabilis* and *mylabris quadripunctata* beetle extract, and methanol (vehicle control for beetle extracts). Cells were harvested after 3 hours of treatment. Phosphorylated and unphosphorylated species of each protein are indicated. **J.** Rhodamine labelled SNAP-tagged NLRP1-DR or immunoblot of TAK1, p38, JNK, ERK, and GAPDH following PhosTag-SDS-Agarose-PAGE from N/TERTs pre-treated with 5 μM emricasan (pan-caspase inhibitor) before stimulation with DMSO (vehicle control), 50 μM cantharidin, and 50 ng/mL TNFα. Cells were harvested after the indicated amount of time. Phosphorylated and unphosphorylated species of each protein are indicated. Error bars represent SEM from three technical replicates, where one replicate refers to an independent sample. Significance values were calculated based on two-way ANOVA followed by Sidak’s test for multiple pairwise comparisons (B). ns, nonsignificant; *P < 0.05; ****P < 0.0001.

**Figure S13.**
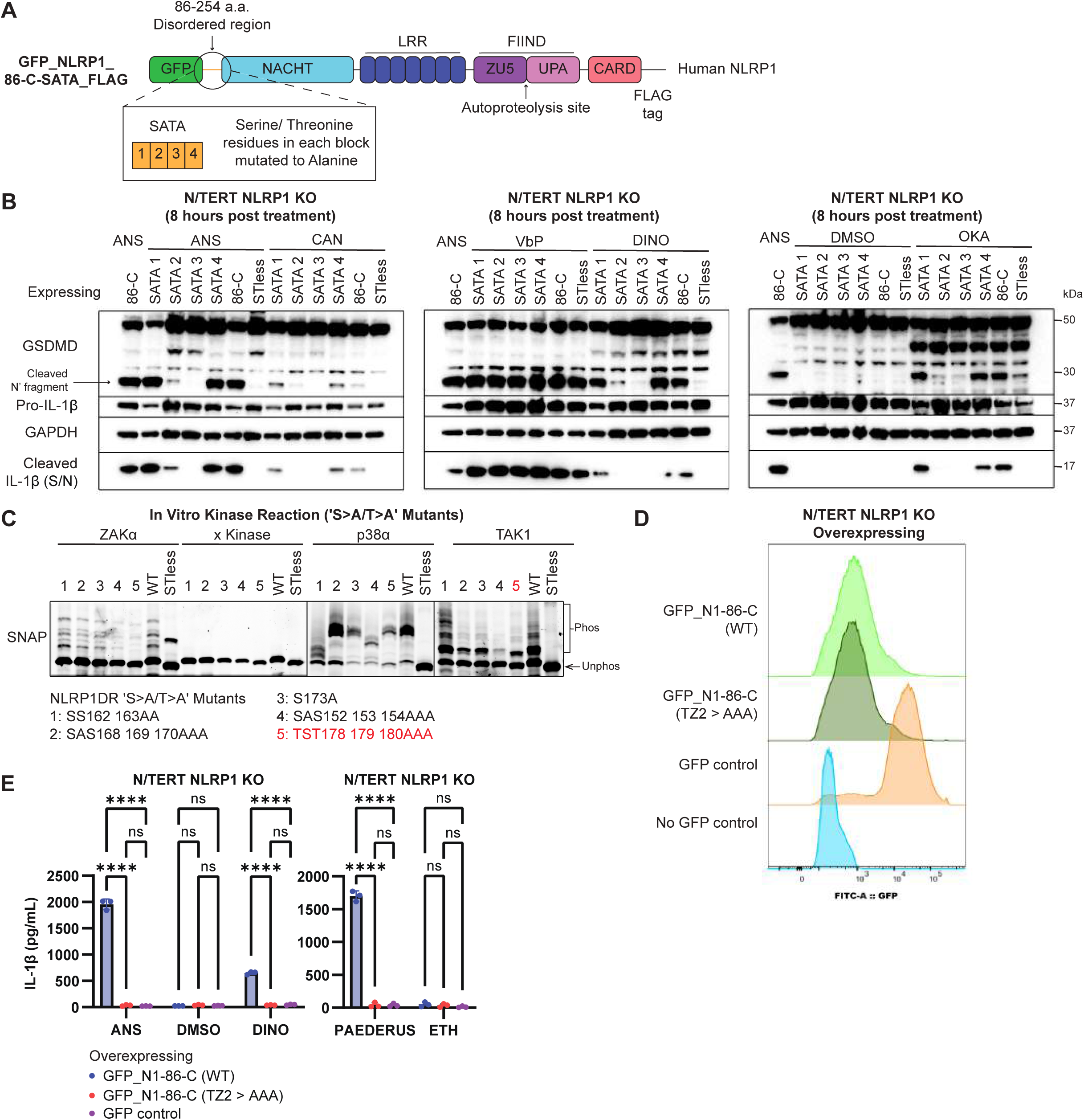
Phosphatase inhibitors require specific sections of the disordered region for activation of the NLRP1 inflammasome. **A.** Domain architecture of human NLRP1. Delta PYD NLRP1 (residues 86-1473a.a.) was cloned into a N’ terminal GFP-tagged and C’ terminal FLAG-tagged vector and overexpressed in N/TERTs. With the same NLRP1 boundaries, another construct was cloned with sections (SATA 1-4) of S/T residues in the disordered region (85-254a.a.) mutated to alanine. The delta PYD construct is referred to as “86-C”, and the disordered region mutants are labelled “SATA1-4”. **B.** Immunoblot of GSDMD (full length and cleaved), Pro-IL-1β, cleaved IL-1β, and GAPDH of NLRP1 KO N/TERTs overexpressing GFP_NLRP1 86-C_FLAG (WT), GFP_NLRP1-86-C_FLAG (SATA1-4), and GFP_NLRP1-86-C_FLAG (STless) following treatment with 1 μM anisomycin, 50 μM cantharidin, 6 μM VbP, 12.5 nM dinophysis, DMSO (vehicle control), and 62.5 nM okadaic acid. Cells, floaters, and cell culture media were harvested after 8 hours of treatment. **C.** Rhodamine labelled SNAP-tagged NLRP1-DR ‘S>A/T>A’ mutants following PhosTag-SDS-Agarose-PAGE. Each mutant had indicated serine or threonine residues mutated to alanine before recombinant expression. These recombinantly expressed mutants were incubated with kinases ZAKα, p38α, and TAK1-TAB1 fusion in a standard in vitro kinase reaction for 60 minutes. Phosphorylated and unphosphorylated species are indicated. The crucial “TZ2” motif is highlighted in red. **D.** Histograms of FITC::GFP fluorescence intensity for the live singlet population of NLRP1 KO N/TERTs overexpressing GFP_NLRP1-86-C_FLAG mutants, displayed as half-offset overlays in FlowJo. **E.** IL-1β ELISA from NLRP1 KO N/TERTs overexpressing GFP_NLRP1-86-C_FLAG (WT), TZ2 mutant, or GFP control following treatment with 1 μM anisomycin, DMSO (vehicle control), 12.5 nM dinophysis, X1500 dilution of *Paederus* beetle extract, and ethanol (vehicle control for beetle extract). Cell culture media was harvested 8 hours after treatment. Error bars represent SEM from three technical replicates, where one replicate refers to an independent sample. Significance values were calculated based on two-way ANOVA followed by Sidak’s test for multiple pairwise comparisons (E). ns, nonsignificant; *P < 0.05; ****P < 0.0001.

**Figure S14.**
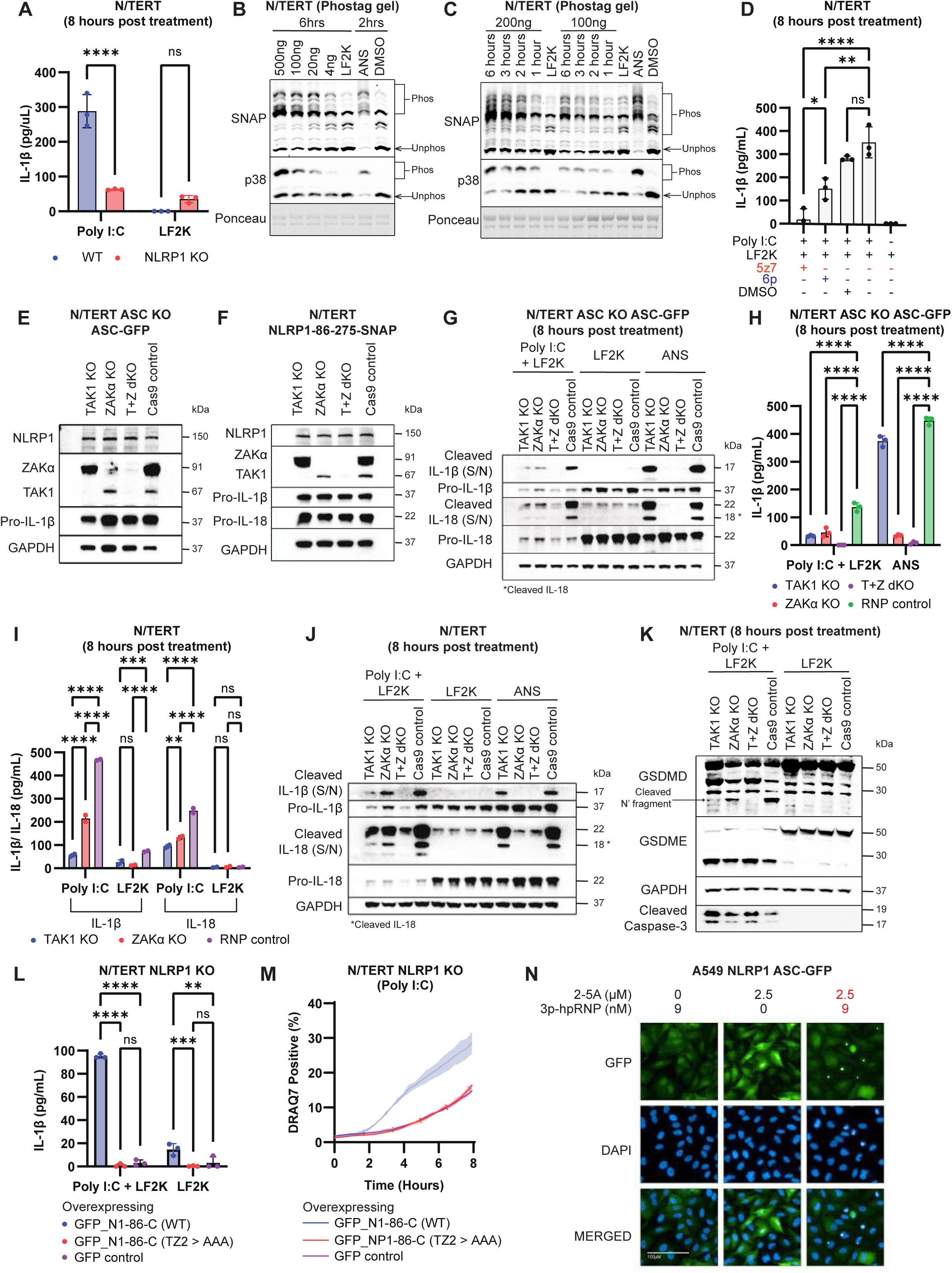
dsRNA/ CHIKV activates the NLRP1 inflammasome via combined actions of TAK1 and ZAKα in immortalized keratinocytes. **A.** IL-1β ELISA from WT and NLRP1 KO N/TERTs following treatment with 200 ng/mL Poly I:C and LF2K (transfection control). Cell culture media was harvested after 8 hours of treatment. **B.** Rhodamine labelled SNAP-tagged NLRP1-DR or immunoblot of p38 following PhosTag-SDS-Agarose-PAGE from N/TERTs pre-treated with 5 μM emricasan (pan-caspase inhibitor) before stimulation with a titration of Poly I:C, LF2K (transfection control), 1 μM anisomycin, and DMSO (vehicle control). Cells were harvested at indicated timepoints. Phosphorylated and unphosphorylated species of each protein are indicated. **C.** Rhodamine labelled SNAP-tagged NLRP1-DR or immunoblot of p38 following PhosTag-SDS-Agarose-PAGE from N/TERTs pre-treated with 5 μM emricasan (pan-caspase inhibitor) before stimulation with 200 ng/mL or 100 ng/mL of Poly I:C, LF2K (transfection control), 1 μM anisomycin, and DMSO (vehicle control). Cells were harvested at indicated timepoints. Phosphorylated and unphosphorylated species of each protein are indicated. **D.** IL-1β ELISA from N/TERTs pre-treated with 1 μM 5-z7-oxozeanol (pan-kinase inhibitor), 1 μM 6p (ZAK inhibitor), or DMSO (vehicle control) before stimulation with 200 ng/mL Poly I:C. Cell culture media was harvested after 8 hours of treatment. **E.** Immunoblot of ZAK, TAK1, Pro-IL-1β, and GAPDH from ASC KO N/TERTs overexpressing ASC-GFP (N/TERT ASC-GFP reporter cells) or **F.** WT N/TERTs overexpressing NLRP1-86-275_SNAP to verify knockout efficiency. **G.** Immunoblot of Pro-IL-1β, cleaved IL-1β, IL-18 (full length and cleaved), and GAPDH or **H.** IL-1β ELISA from TAK1 KO, ZAKα KO, T+Z dKO, and RNP control N/TERT ASC-GFP reporter cells following treatment with 200 ng/mL Poly I:C, LF2K (transfection control), and 1 μM anisomycin. Cells and cell culture media were harvested 8 hours after treatment. **I.** IL-1β ELISA or **J.** immunoblot of Pro-IL-1β, cleaved IL-1β, IL-18 (full length and cleaved), and GAPDH from TAK1 KO, ZAKα KO, T+Z dKO, and RNP control WT N/TERTs overexpressing NLRP1-86-275_SNAP following treatment with 200 ng/mL Poly I:C, LF2K (transfection control), and 1 μM anisomycin. Cells and cell culture media were harvested 8 hours after treatment. **K.** Immunoblot of GSDMD (full length and cleaved), GSDME (full length and cleaved), GAPDH, and cleaved caspase-3 from WT N/TERTs overexpressing NLRP1-86-275_SNAP following treatment with 200 ng/mL Poly I:C and LF2K (transfection control). Cells, floaters and cell culture media were harvested 8 hours after treatment. **L.** IL-1β ELISA or **M.** Quantification of percentage of DRAQ7-positive cells from NLRP1 KO N/TERTs overexpressing GFP_NLRP1-86-C_FLAG (WT), TZ2 mutant, or GFP control following treatment with 200 ng/mL Poly I:C and LF2K (transfection control). Cell culture media was harvested 8 hours after treatment.| **N.** Representative fluorescence images of A549 NLRP1 ASC-GFP reporter cells following transfection of 3p-hpRNA and 2-5A individually or in combination. Cells were fixed and stained with DAPI after 3 hours of treatment. Error bars represent SEM from three technical replicates, where one replicate refers to an independent sample. Significance values were calculated based on one-way ANOVA followed by Dunnett’s test (D) or two-way ANOVA followed by Sidak’s test for multiple pairwise comparisons (A, H, I, and L). ns, nonsignificant; *P < 0.05; ****P < 0.0001.

**Figure S15.**
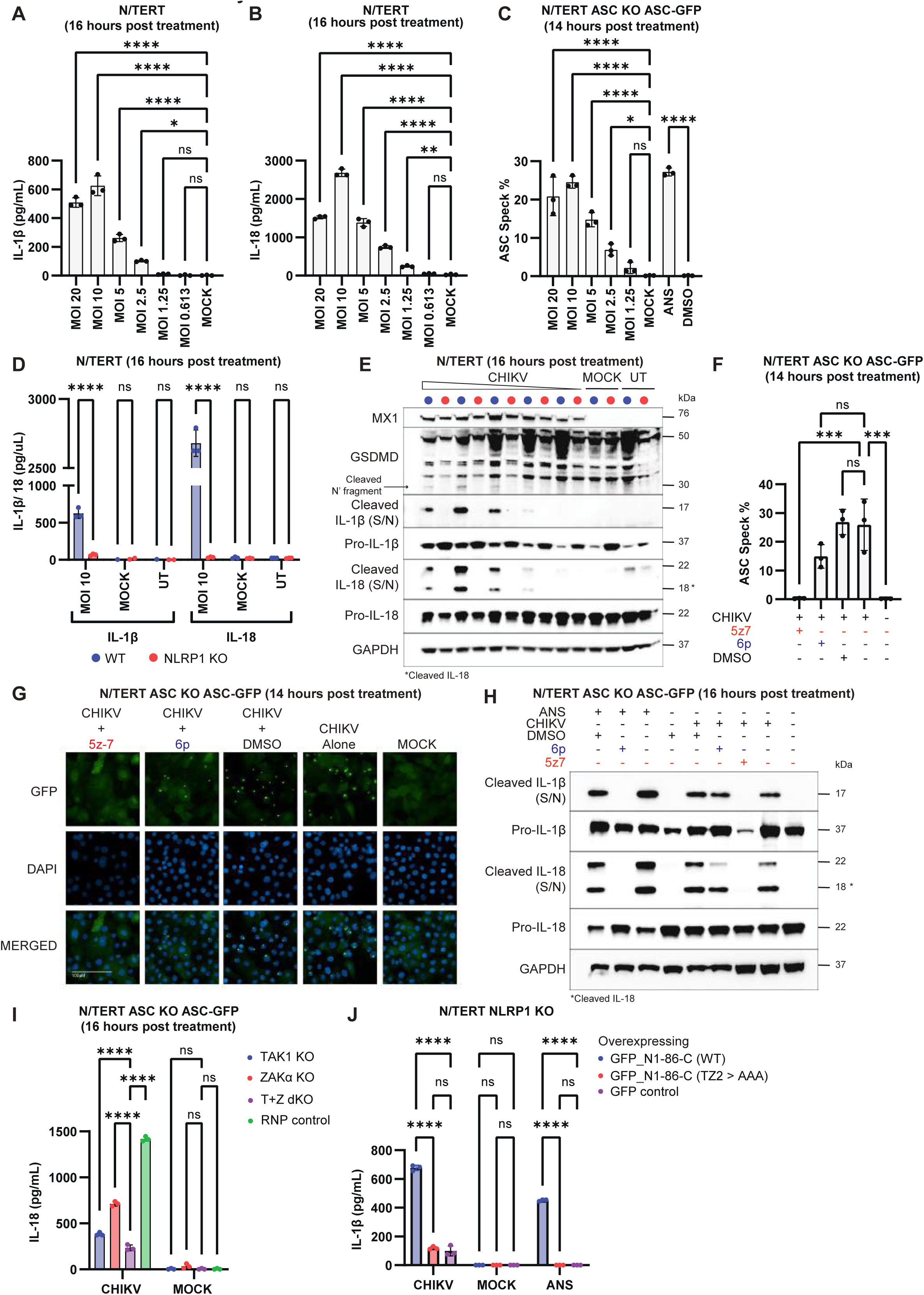
CHIKV activates the NLRP1 inflammasome in immortalized keratinocytes. **A-B.** IL-1β and IL-18 ELISA **C.** percentage of ASC-GFP speck forming N/TERTs following treatment with MOI titration of CHIKV, MOCK (vehicle control for CHIKV), 1 μM anisomycin, and DMSO (vehicle control for anisomycin). Cell culture media was harvested or cells were fixed and stained with DAPI after 16 or 14 hours of treatment. **D.** IL-1β and IL-18 ELISA and **E.** immunoblot of GSDMD (full length and cleaved), Pro-IL-1β, cleaved IL-1β, IL-18 (full length and cleaved), and GAPDH from WT and NLRP1 KO N/TERTs following treatment with CHIKV (MOI 10 or a titration) and mock (vehicle control for CHIKV). Cell culture media was harvested after 16 hours of treatment. **F.** Percentage of ASC-GFP speck forming and **G.** representative fluorescence images **H.** immunoblot of Pro-IL-1β, cleaved IL-1β, IL-18 (full length and cleaved), and GAPDH from N/TERT ASC-GFP reporter cells pre-treated with 1 μM 5-z7-oxozeanol (pan-kinase inhibitor), 1 μM 6p (ZAK inhibitor), or DMSO (vehicle control) before stimulation with CHIKV (MOI 10). Cells and cell culture media were harvested or cells were fixed and stained with DAPI after 16 or 14 hours of treatment. **I.** IL-18 ELISA from TAK1 KO, ZAKα KO, T+Z dKO, and RNP control N/TERT ASC-GFP reporter cells following treatment with CHIKV (MOI 10) and mock (vehicle control for CHIKV). Cell culture media was harvested after 16 hours of treatment. **J.** IL-1β ELISA from NLRP1 KO N/TERTs overexpressing GFP_NLRP1-86-C_FLAG (WT), TZ2 mutant, or GFP control following treatment with CHIKV (MOI 10), mock (vehicle control for CHIKV), and 1 μM anisomycin. Cell culture media was harvested after 16 hours of treatment. Error bars represent SEM from three technical replicates, where one replicate refers to an independent sample. Significance values were calculated based on one-way ANOVA followed by Dunnett’s test (A-C, and F) or two-way ANOVA followed by Sidak’s test for multiple pairwise comparisons (D, I, and J). ns, nonsignificant; *P < 0.05; ****P < 0.0001.

**Supplementary Table 1.**
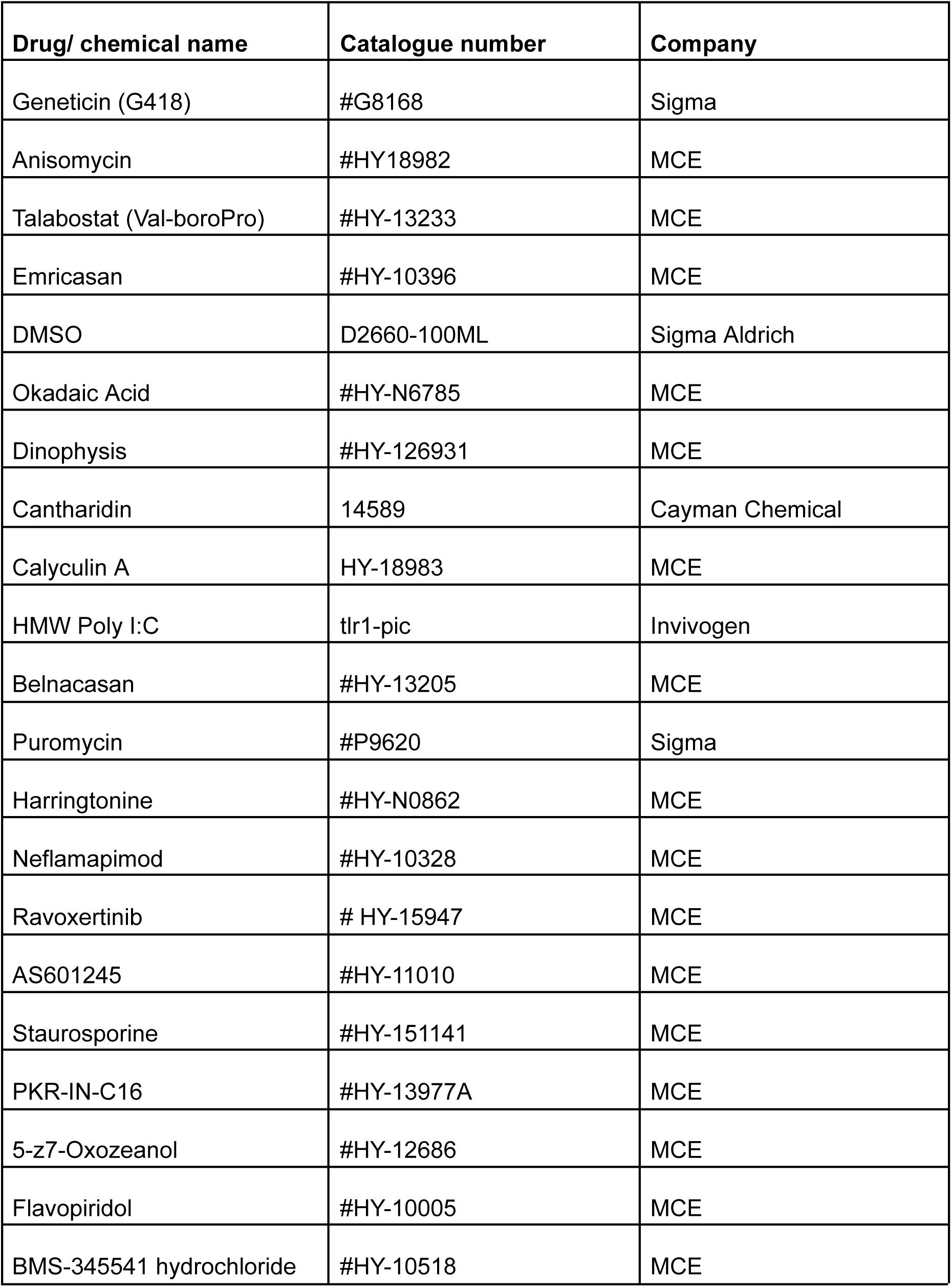
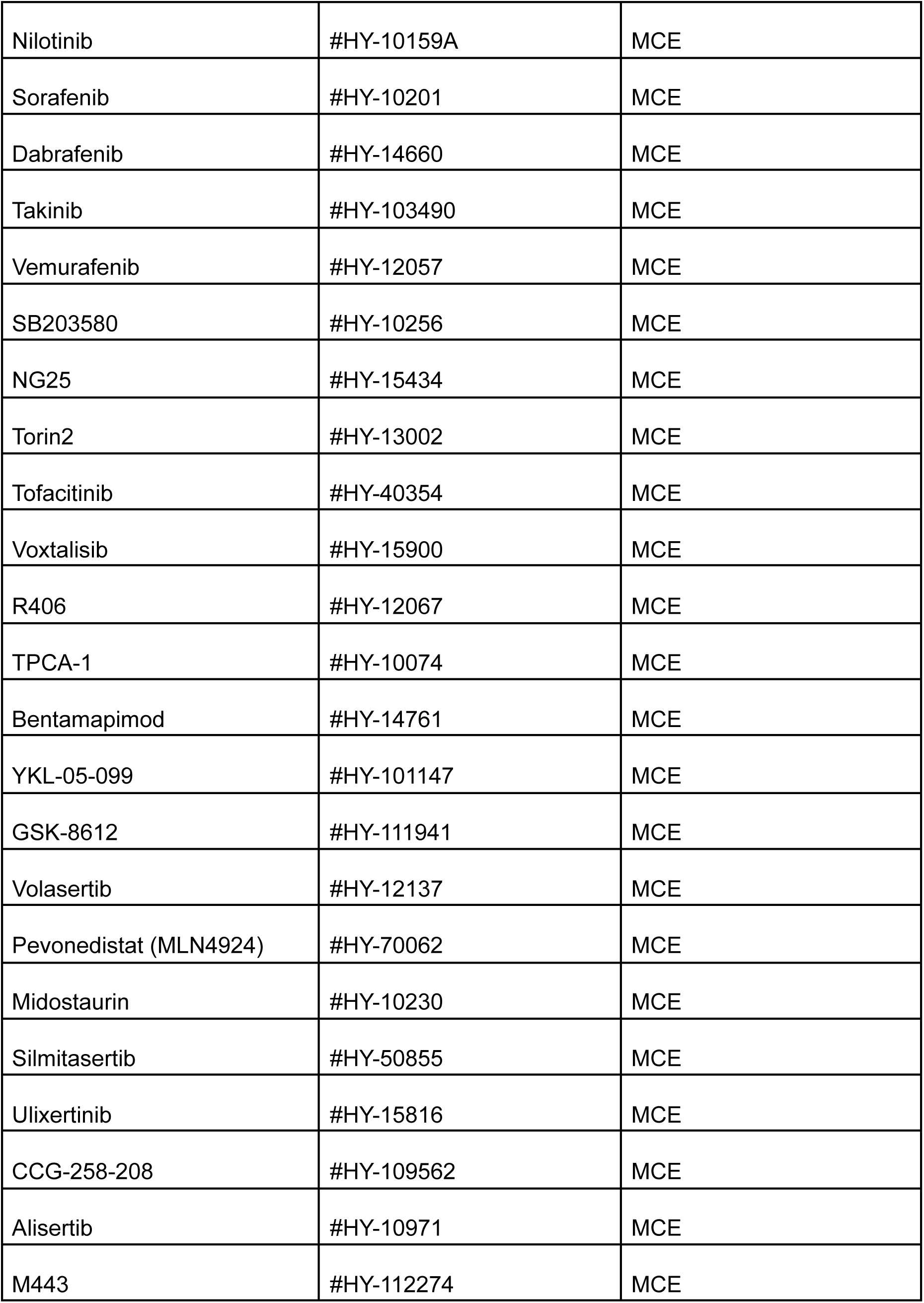
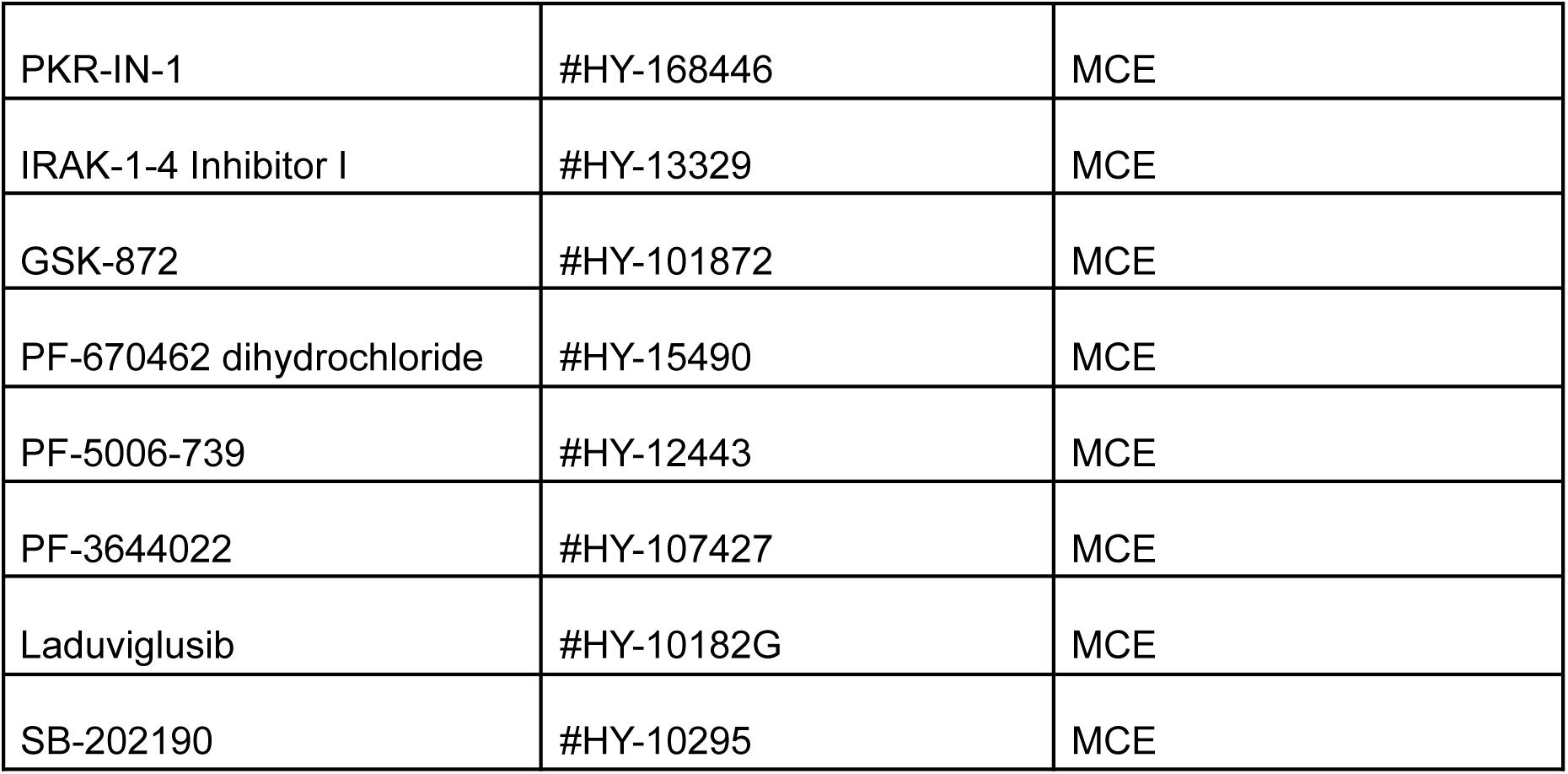
: List of Drugs and Chemicals

**Supplementary Table 2.** : List of Primary and Secondary Antibodies

| **Antibody** | **Catalogue number** |
| --- | --- |
| Cleaved PARP1 | Cell Signalling Technology, #9541 |
| Cleaved CASP3 | Cell Signalling Technology, #9664 |
| GSDME | Abcam, ab215191 |
| GSDMD-FL | Abcam, ab219800 |
| GSDMD-FL | Novus, NBP2-33422 |
| Cleaved IL1β (p17) | Cell Signalling Technology, #83186S |
| IL-1β | R&D Systems, MAB201-100 |
| IL-18 | Abcam, ab243091 |
| NLRP1 (NT) | Biolegend, 9F9B12 |
| GAPDH | Santa Cruz Biotechnology, #sc-47724 |
| p38 | Abcam, ab32142 |
| phospho-p38 | Cell Signalling Technology, #4511 |
| JNK | Cell Signalling Technology, #9252 |
| phospho-JNK | Cell Signalling Technology, #4668 |
| ZAK | Bethyl Laboratories, A301-993A |
| TAK1 | Cell Signalling Technology, #5206 |
| MX1 | Cell Signalling Technology, #37849 |
| HA | Santa Cruz Biotechnology, #sc-805 |
| GFP | Abcam, ab290 |
| FLAG | SigmaAldrich, #F3165 |
| ASC | Adipogen, #AL-177 |
| CASP1 | Santa Cruz Biotechnology, #sc-622 |
| Goat anti-mouse IgG HRP-conjugated | Jackson ImmunoResearch, 115-035-166 |
| Goat anti-rabbit IgG HRP-conjugated | Jackson ImmunoResearch, 111-035-144 |
| Donkey anti-sheep IgG HRP-conjugated | Jackson ImmunoResearch, 713-035-147 |

**Supplementary Table 3.** : MRM table for the detection of pederin, pseudopederin and verapamil.

| Metabolites | Polarity | Parent (m/z) | Product (m/z) | Quantifier/Qualifier | Collision Energy (eV) | RF Lens (V) |
| --- | --- | --- | --- | --- | --- | --- |
| Pederin<br>[M + Na]<sup>+</sup> | Positive | 526.3 | 526.3 | Quantifier | 5 | N/A |
|  |  |  | 494.27 | Qualifier | 25 | N/A |
|  |  |  | 462.25 | Qualifier | 25 | N/A |
|  |  |  | 340.17 | Qualifier | 25 | N/A |
|  |  |  | 297.17 | Qualifier | 25 | N/A |
| Pseudopederin<br>[M + Na]<sup>+</sup> | Positive | 512.3 | 512.3 | Quantifier | 5 | N/A |
|  |  |  | 494.27 | Qualifier | 25 | N/A |
|  |  |  | 480.26 | Qualifier | 25 | N/A |
|  |  |  | 462.25 | Qualifier | 25 | N/A |
|  |  |  | 340.17 | Qualifier | 25 | N/A |
|  |  |  | 297.17 | Qualifier | 25 | N/A |
|  |  |  | 282.17 | Qualifier | 25 | N/A |
| Verapamil | Positive | 455.2 | 165.089 | Quantifier | 30.02 | 207 |
|  |  |  | 303.167 | Qualifier | 26.4 | 207 |

**Supplementary Table 4.** : MRM table for the detection of cantharidin and cantharidic acid

| Metabolites | Polarity | Parent (m/z) | Product (m/z) | Quantifier / Qualifier | Collision Energy (eV) | RF Lens (V) |
| --- | --- | --- | --- | --- | --- | --- |
| Cantharidin<br>[M + H]<sup>+</sup> | Positive | 197.112 | 107.071 | Quantifier | 10.86 | 101 |
|  |  |  | 95.1 | Qualifier | 17.43 | 101 |
|  |  |  | 123.096 | Qualifier | 14.27 | 101 |
|  |  |  | 151.1 | Qualifier | 10.1 | 101 |
|  |  |  | 169.175 | Qualifier | 9.68 | 101 |
|  |  |  | 179.079 | Qualifier | 10.81 | 101 |
| Cantharidic Acid<br>[M – H<sub>2</sub>O + H]<sup>+</sup> | Positive | 197.138 | 151.125 | Quantifier | 10.39 | 99 |
|  |  |  | 95.054 | Qualifier | 17.81 | 99 |
|  |  |  | 105.071 | Qualifier | 22.27 | 99 |
|  |  |  | 107.071 | Qualifier | 11.28 | 99 |
|  |  |  | 123.125 | Qualifier | 14.56 | 99 |
|  |  |  | 169.125 | Qualifier | 10.14 | 99 |
|  |  |  | 179.079 | Qualifier | 10.81 | 99 |

